# Discovery and biosynthesis of gladiochelins: unusual lipodepsipeptide siderophores from *Burkholderia gladioli*

**DOI:** 10.1101/2020.06.16.153940

**Authors:** Yousef Dashti, Ioanna T. Nakou, Alex J. Mullins, Gordon Webster, Xinyun Jian, Eshwar Mahenthiralingam, Gregory L. Challis

**Affiliations:** Department of Chemistry, University of Warwick, Coventry CV4 7AL, United Kingdom; Microbiomes, Microbes and Informatics Group, Organisms and Environment Division, School of Biosciences, Cardiff University, Cardiff, UK; Warwick Integrative Synthetic Biology Centre, University of Warwick, Coventry CV4 7AL, United Kingdom; Department of Biochemistry and Molecular Biology, Monash University, Clayton, VIC 3800, Australia; The Centre for Bacterial Cell Biology, Biosciences Institute, Medical School, Newcastle University, Newcastle upon Tyne, NE2 4AX, UK

## Abstract

*Burkholderia* is a genus of diverse Gram-negative bacteria that includes several opportunistic pathogens. Siderophores, which transport iron from the environment into microbial cells, are important virulence factors in most pathogenic *Burkholderia* species. However, it is widely believed that *Burkholderia gladioli*, which can infect the lungs of cystic fibrosis (CF) sufferers, does not produce siderophores. *B. gladioli* BCC0238, isolated from the lung of a CF patient, produces two novel metabolites in a minimal medium containing glycerol and ribose as carbon sources. HPLC purification, followed by detailed spectroscopic analyses, identified these metabolites as unusual lipodepsipeptides containing a unique citrate-derived fatty acid and a rare dehydro-β-alanine residue. The absolute configurations of the amino acid residues in the two metabolites was elucidated using Marfey’s method and the gene cluster responsible for their biosynthesis was identified by bioinformatics and insertional mutagenesis. In-frame deletions and enzyme activity assays were used to investigate the functions of several proteins encoded by the biosynthetic gene cluster, which was found in the genomes of most *B. gladioli* isolates, suggesting that its metabolic products play an important role in the growth and/or survival of the species. The Chrome Azurol S (CAS) assay showed the metabolites bind ferric iron and that this supresses their production when added to the growth medium. Moreover, a gene encoding a TonB-dependent ferric-siderophore receptor is adjacent to the biosynthetic genes. Together, these observations suggest that these metabolites likely function as siderophores in *B. gladioli*.

## Introduction

Iron is an essential element for most organisms. In biological systems, iron is mainly present in two oxidation states: ferrous and ferric.^1^ The interconversion of the ferrous and ferric oxidation states is essential for a variety of redox reactions and electron transfer processes in cells.^2^ However, despite the relatively high abundance of iron in the Earth’s crust, it is not readily bioavailable. This is because it is predominantly in the ferric form, which has low aqueous solubility.^3^ Moreover, sequestration by proteins such as hemoglobin, ferritin, transferrin, and lactoferrin also contributes to low iron availability in mammals.^3^ As a consequence, microorganisms have evolved several strategies for obtaining iron from the environment and their hosts. One strategy involves the production of siderophores, specialised metabolites with a high affinity for ferric iron.^4^ In Gram-negative bacteria, ferric-siderophore complexes are typically transported into the periplasm by TonB-dependent outer membrane receptors. Periplasmic binding proteins then shuttle the ferric-siderophore complexes to the inner membrane, where ATP-binding cassette (ABC) transporters transfer them into the cytoplasm.^5-6^ Once in the cytoplasm, iron is released from the ferric-siderophore complex by reduction to the ferrous form and/or hydrolytic cleavage of the ligand.^5^

Several investigations of pathogenic microorganisms have identified a strong link between siderophore production and virulence.^7-8^ In the early stages of infection, siderophores play a key role in sequestering iron from host tissues. The identification of siderophores produced by pathogenic microorganisms is therefore an important field of study because interference with siderophore-mediated iron uptake is an attractive strategy for attenuating virulence.^9^ The discovery of novel siderophores is also relevant to other therapeutic applications. For example, desferrioxamine B (marketed as Desferal), a siderophore produced by numerous *Streptomyces* species,^10^ is used to treat iron overload resulting from multiple blood transfusions and aluminium toxicity due to dialysis. However, because Desferal is administered by injection and has multiple side effects, orally-available alternatives with fewer side effects are actively being sought.^11^ In addition to clinical uses, siderophores show promise for environmental and biotechnological applications.^12-13^

*Burkholderia* is a genus of highly diverse Gram-negative bacteria. Species belonging to this genus have been isolated from soil, water, plants, animals and humans.^14^ Several investigations have shown that they are prolific producers of specialised metabolites,^15-24^ and genome sequence analyses have highlighted their underexplored biosynthetic potential.^25^ To date, pathogenic *Burkholderia* species have been reported to produce siderophores belonging to three distinct structural classes (ornibactins/malleobactins, pyochelin, and cepaciachelin; Figure 1).^26-32^ Members of the *cepacia* complex and *pseudomallei* group produce one or more of these siderophores. In contrast, *Burkholderia gladioli* does not produce any of these metabolites and is believed to employ siderophore-independent mechanisms for iron acquisition.^33^

**Figure 1.**
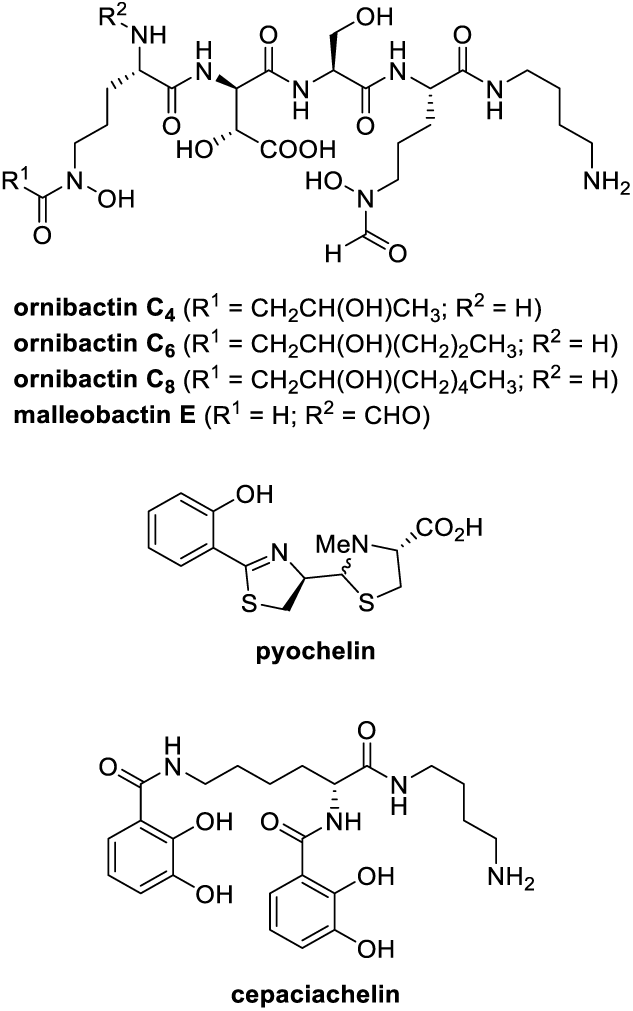
Structure of ornibactins, malleobactin E, pyochelin, and cepaciachelin identified from pathogenic strains of *Burkholderia*.

We have recently discovered that *B. gladioli* BCC0238, isolated from the sputum of a child with cystic fibrosis (CF), produces several specialised metabolites. These include gladiolin, a novel macrolide with potent activity against drug resistant *Mycobacterium tuberculosis* clinical isolates,^23^ and icosalide A1,^34^ an asymmetric lipopeptodiolide antibiotic originally isolated from an *Aureobasidium* species fungus,^35^ but subsequently shown to be produced by various strains of *B. gladioli* including one associated with the fungus.^34, 36^ Analysis of the *B. gladioli* BCC0238 complete genome sequence identified several gene clusters encoding cryptic nonribosomal peptide synthetase (NRPS), polyketide synthase (PKS) and hybrid NRPS-PKS assembly lines.^21^ Here we report the discovery of gladiochelins A and B, novel lipodepsipeptides containing a unique citrate-derived fatty acid and a rare dehydro-β-alanine residue, as the metabolic products of one of these gene clusters. A combination of comparative bioinformatics analyses, targeted gene deletions, and enzyme activity assays elucidated several key steps in the biosynthesis of these unusual natural products, which we propose based on several lines of evidence function as siderophores.

## Results and discussion

### Isolation and structure elucidation of the gladiochelins

*B. gladioli* BCC0238 produces gladiolin and icosalide A1 when grown on a solid minimal medium containing glycerol as the sole carbon source.^23, 34^ Two novel metabolites were identified by UHPLC-ESI-Q-TOF-MS analysis of extracts from cultures of a gladiolin non-producing mutant of *B. gladioli* BCC0238 (containing an in-frame deletion in the region of *gbnD1* encoding the ER domain) grown on BSM medium containing ribose as an additional carbon source alongside glycerol. The metabolites were isolated as amorphous solids from ethyl acetate extracts of 3-day cultures (1L total volume) using semi-preparative HPLC, and their planar structures were elucidated using a combination of HR-ESI-MS and 1D/2D NMR experiments (Tables S2 and S3, Figures S3-S14).

The molecular formulae of the two metabolites, which we named gladiochelins A **1** and B **2**, respectively, were established as C_39_H_61_N_5_O_14_ (calculated for C_39_H_62_N_5_O_14^+^_: 824.4288, found: 824.4284) and C_40_H_63_N_5_O_14_ (calculated for C_40_H_64_N_5_O_1_4^+^: 838.4444, found: 838.4447) using positive ion ESI-Q-TOF-MS (Figure S17). Four resonances with chemical shifts characteristic of protons attached to C-2 in α-amino acids (*δ*_H_ 4.17, 4.79, 5.36, and 5.52) correlated with signals attributable to exchangeable N-H protons (*δ*_H_ 8.17, 8.70, 9.91, and 9.99, respectively) in the COSY spectrum of gladiochelin A **1**. Further analysis of ^1^H NMR and COSY spectra identified four α-amino acid spin systems, which in conjunction with HSQC and HMBC data, were assigned to a valine residue, a homoserine (Hse) residue and two serine residues in the sequence Ser-Hse-Val-Ser (Figure 2). A ^3^*J*_CH_ correlation from the β-protons of the N-terminal Ser (N-Ser) residue (*δ*_H_ 4.73 and 4.90) to the carbonyl carbon of the C-terminal Ser (C-Ser) residue (*δ*_C_ 170.0) established the presence of a tetradepsipeptide (Figure 2). Analysis of the COSY and HMBC data also revealed a dehydro-β-alanine (Dba) residue, with a coupling constant of 14.0 Hz between the α and β protons indicating that the double bond is *E*-configured. The only other natural products reported to contain a Dba residue are the enamidonins isolated from *Streptomyces* species.^37-38 2^*J*_CH_ correlations were observed between the N-H proton of the N-Ser residue in the tetradepsipeptide (*δ*_H_ 9.99) and the carbonyl carbon of the Dba residue (*δ*_C_ 168.3), and the N-H proton of Dba residue (*δ*_H_ 11.56) and the carbonyl carbon of a fatty acid residue (*δ*_C_ 165.1). This indicates that the Dba residue links the fatty acid residue to the tetradepsipeptide. The fatty acid residue is very unusual, with Z-configured double bonds spanning C2-C3 and C11-C12 (based on CH=CH coupling constants of 11.5 and 13.5 Hz, respectively), and two carboxyl acid groups and a methoxy group appended to its tail (Figure 2).

**Figure 2.**
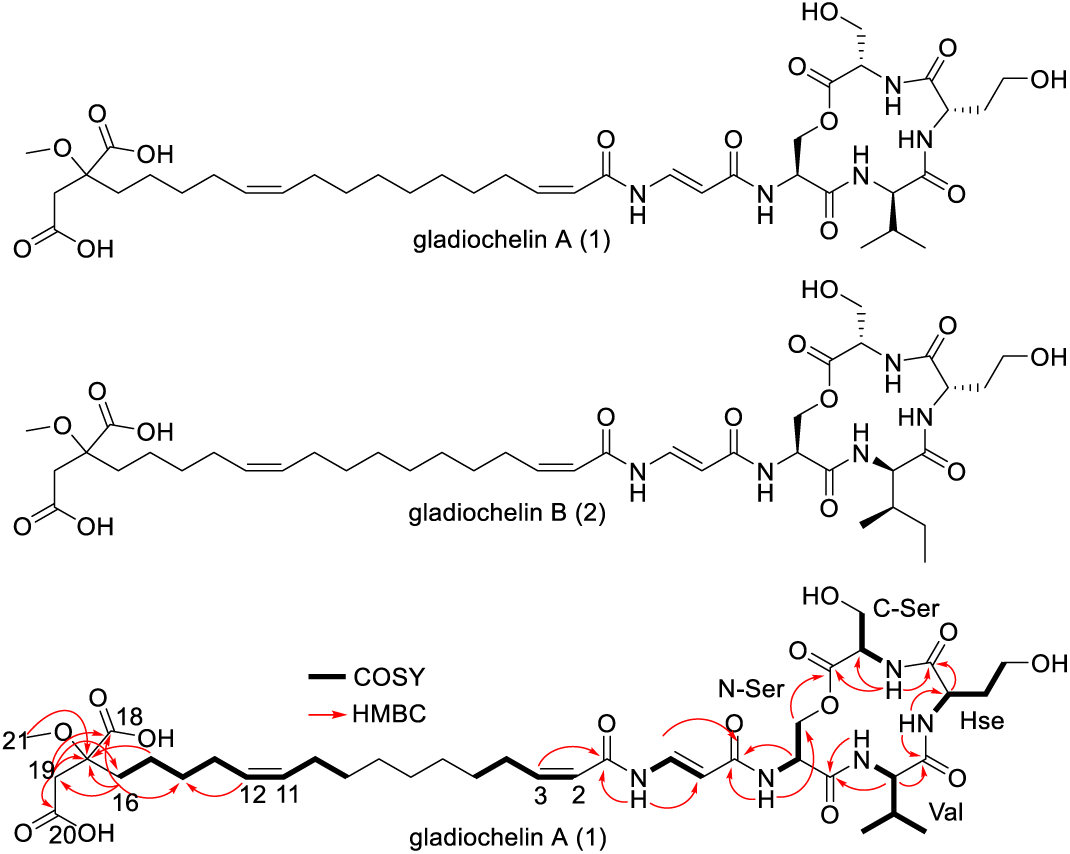
Structures of gladiochelin A (**1**) and B (**2**). COSY and key HMBC correlations are shown for gladiochelin A.

The absolute configurations of the α-amino acid residues in gladiochelin A **1** were determined as L-Ser-D-Val-L-Hse-L-Ser using Marfey’s method^39^ (Figure S15). Comparison of the NMR spectroscopic data for gladiochelins A **1** and B **2** showed that the Val residue in the latter is substituted by an Ile residue in the former, consistent with the difference of CH_2_ in the molecular formulae. The absolute configurations of the α-amino acids in gladiochelin B (**2**) were similarly shown to be L-Ser-D-*allo*-Ile-L-Hse-L-Ser by Marfey’s method (Figure S15), using a C3 column instead of a C18 column to separate the Marfey’s derivatives of D-Ile and D-*allo*-Ile (Fig. S15).^40^

Overall, the structures of gladiochelins A **1** and B **2** are highly unusual, containing a unique fatty acid residue with several polar functional groups appended to its tail, linked via a rare Dba residue to a depsitetrapeptide. We thus sought to develop an understanding of gladiochelin biosynthesis, focusing on the origin of the starter unit for the fatty acid assembly and the mechanism of Dba incorporation.

### Gladiochelin biosynthesis

Bioinformatics analysis of the *B. gladioli* BCC0238 genome sequence identified the putative galdiochelin (*gcn*) biosynthetic gene cluster (Figure 3 and Table 1), which contains a gene (*gcnH*) encoding a tetramodular NRPS. Sequence analysis of the adenylation (A) domains in this NRPS indicated that modules 1-4 activate L-Ser, L-Val/L-Ile, L-Hse and L-Ser, respectively (Table S4),^41-42^ suggesting the NRPS assembles the depsipeptide portion of the gladiochelins. Similarly, phylogenetic analysis of the GcnH condensation (C) domains indicated that they belong to the following groups.^43^ Module 1: chain initiating (C_I_ – links external acyl donor and L-configured acyl acceptor); modules 2 and 4 – chain elongating (^L^C_L_ – links L-configured acyl donor and acceptor); and module 3 – bifunctional condensation / epimerization (C_E_ – epimerizes acyl donor to D-configuration and links with L-configured acyl acceptor) (Figure 3). This is consistent with the structure elucidation data, which indicate that the gladiochelins contain L-Ser, L-Hse and D-Val/D-*allo*-Ile.

**Table 1.** Proteins encoded by the gladiochelin biosynthetic gene cluster and their predicted functions.

| Protein | Locus tag | Predicted function |
| --- | --- | --- |
| GcnA | BCC1622_03825/BCC0238_03115 | LysR family transcriptional regulator |
| GcnB | BCC1622_03826/BCC0238_03114 | acyl-CoA synthetase |
| GcnC | BCC1622_03827/BCC0238_03113 | acyl carrier protein |
| GcnD | BCC1622_03828/BCC0238_03112 | non-heme di-iron-dependent oxygenase |
| GcnE | BCC1622_03829/BCC0238_03111 | flavin-dependent oxidoreductase |
| GcnF | BCC1622_03830/BCC0238_03110 | membrane-associated fatty acid desaturase |
| GcnG | BCC1622_03831/BCC0238_03109 | MFS transporter |
| GcnH | BCC1622_03832/BCC0238_03108 | nonribosomal peptide synthetase |
| GcnI | BCC1622_03833/BCC0238_03107 | MbtH-like protein |
| GcnJ | BCC1622_03834/BCC0238_03106 | SnoaL-like protein |
| GcnK | BCC1622_03835/BCC0238_03105 | hypothetical protein |
| GcnL | BCC1622_03836/BCC0238_03104 | acyl-ACP desaturase |
| GcnM | BCC1622_03837/BCC0238_03103 | 3-oxoacyl-ACP synthase III (C→S) |
| GcnN | BCC1622_03838/BCC0238_03102 | PLP-dependent *iso*-aspartyl-ACP decarboxylase |
| GcnO | BCC1622_03839/BCC0238_03101 | *iso*-aspartyl-ACP synthetase |
| GcnP | BCC1622_03840/BCC0238_03100 | 3-oxoacyl-ACP synthase III (C→T) |
| GcnQ | BCC1622_03841/BCC0238_03099 | flavin-dependent acyl-CoA dehydrogenase |
| GcnR | BCC1622_03842/BCC0238_03098 | citrate synthase |
| GcnS | BCC1622_03843/BCC0238_03097 | O-methyltransferase |
| GcnT | BCC1622_03844/BCC0238_03096 | α-ketoglutarate and non-heme iron-dependent oxygenase |
| GcnU | BCC1622_03817/BCC0238_03123 | TonB-dependent ferric-siderophore receptor |

**Figure 3.**
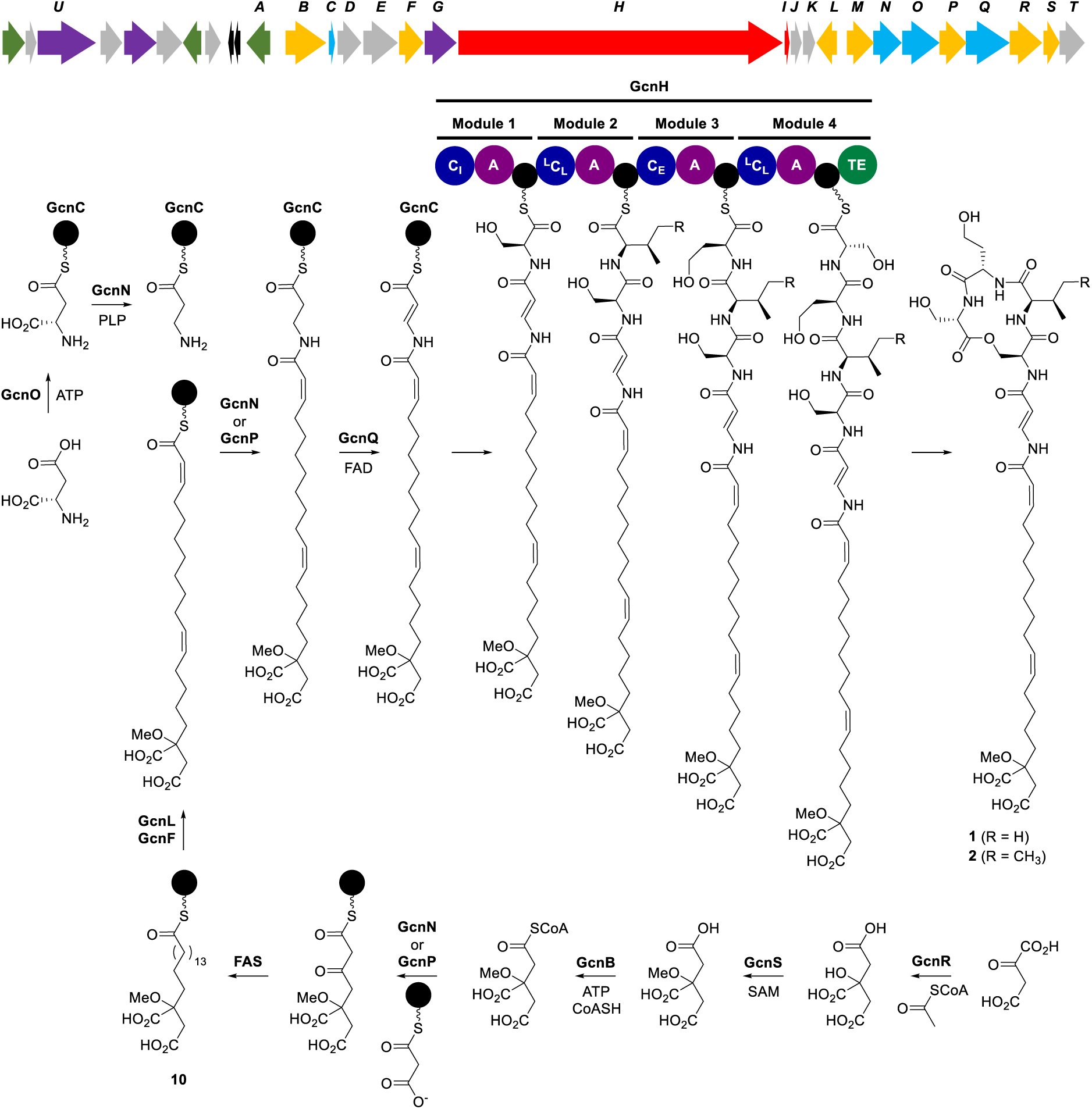
Organization of the gladiochelin biosynthetic gene cluster (*gcnA-T*) and proposed pathway for gladiochelin biosynthesis. The precise timing of O-methylation of the citrate-derived starter unit for fatty acid biosynthesis remains to be established, but is postulated to be an early step. Abbreviations are as follows: A, adenylation domain; TE, thioesterase domain; C_I_, chain initiating condensation domain; ^L^C_L_, chain elongating condensation domain utilizing two L-configured aminoacyl thioesters; C_E_, bifunctional epimerization-condensation domain; FAS, fatty acid synthase. Acyl carrier proteins (ACPs) and Peptidyl carrier protein (PCP) domains are shown as black circles.

To verify the involvement of this gene cluster in gladiochelin biosynthesis, we inactivated *gcnH* in *B. gladioli* BCC0238 Δ*gbnD1_ER* via insertional mutagenesis. LC-MS analysis confirmed that the production of gladiochelins A **1** and B **2** was abolished in the mutant (Figure 4). Putative functions were assigned to the proteins encoded by the *gcnA*-*gcnT* genes flanking *gcnH* on the basis of sequence analyses (Table 1). This enabled us to propose a plausible pathway for gladiochelin biosynthesis (Figure 3). To validate this pathway experimentally, we envisioned examining the effect of in-frame deletions in key biosynthetic genes on gladiochelin production. However, this proved to be challenging and time consuming in *B. gladioli* BCC0238 (see supporting information), so we screened other genome-sequenced *B. gladioli* isolates containing the *gcn* locus (see below) to establish whether they (i) produce the gladiochelins and (ii) are amenable to construction of in-frame deletions. *B. gladioli* BCC1622 (also isolated from a CF lung infection) was found to meet both criteria and an analogous gladiolin non-producing mutant of this strain (*B. gladioli* BCC1622 Δ*gbnD1_ER*) was created to enable in-depth functional studies of selected gladiochelin biosynthetic genes.

**Figure 4.**
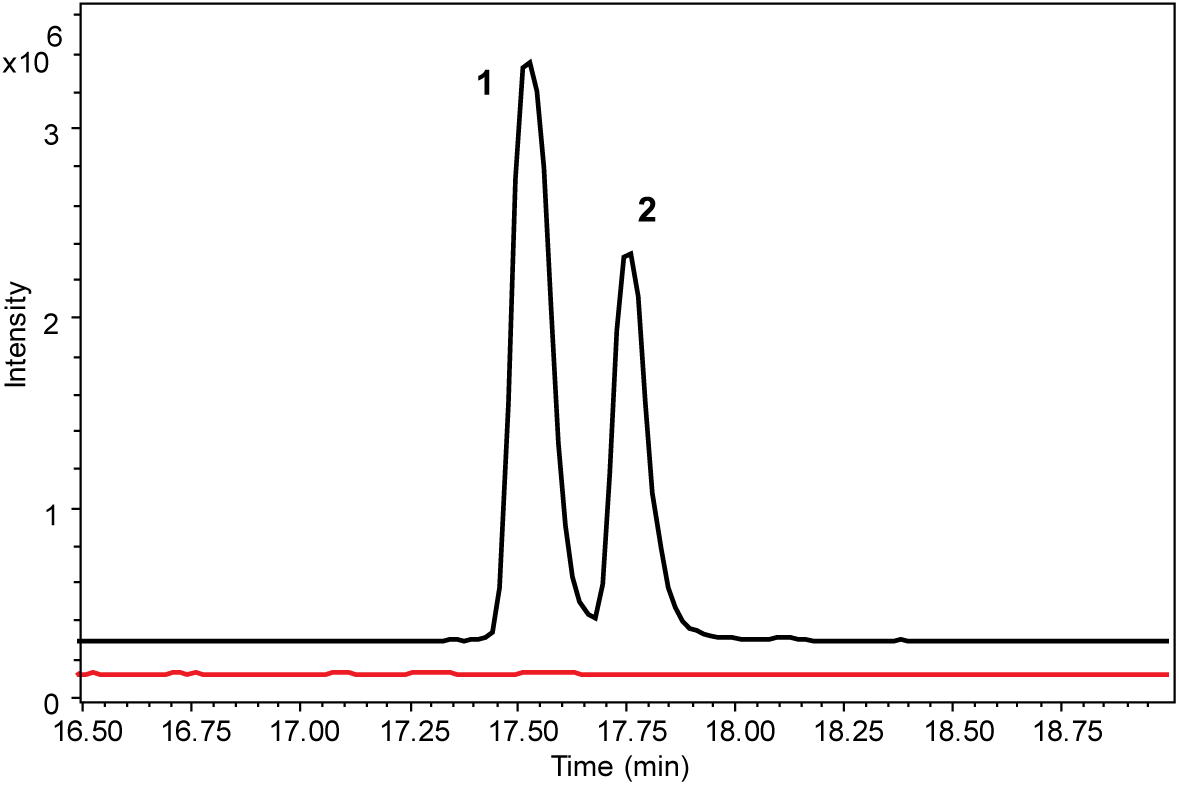
Comparison of the extracted ion chromatograms for *m/z* = 824.4288 (**1**) and 838.4444 (**2**) from UHPLC-ESI-Q-TOF MS analyses of culture extracts from *B. gladioli* BCC0238 Δ*gbnD1_ER* (black) and *B. gladioli* BCC0238 Δ*gbnD1_ER-*Ω*gcnH* (red). Insertional mutagenesis in *gcnH* confirmed the involvement of the *gcn* cluster in gladiochelin production.

GcnR shows sequence similarity to citrate synthase, a key enzyme in the Krebs cycle, which catalyzes the condensation of acetyl-CoA with oxaloactetate and hydrolysis of the resulting thioester to form citrate (Figure 5a).^44^ This suggests that citryl-CoA may serve as the starter unit for assembly of the unusual fatty acid residue incorporated into the gladiochelins (Figure 3). To verify the function of GcnR, we overproduced it in *E. coli* as an N-terminal His_6_-fusion and purified it to homogeneity (see Supporting Information). Incubation of the purified protein with oxaloacetate (**4**) and acetyl-CoA (**5**) for two hours at room temperature resulted in complete conversion to products with molecular formulae corresponding to citric acid (**6**) and coenzyme A **7** (Figure 5b).

**Figure 5.**
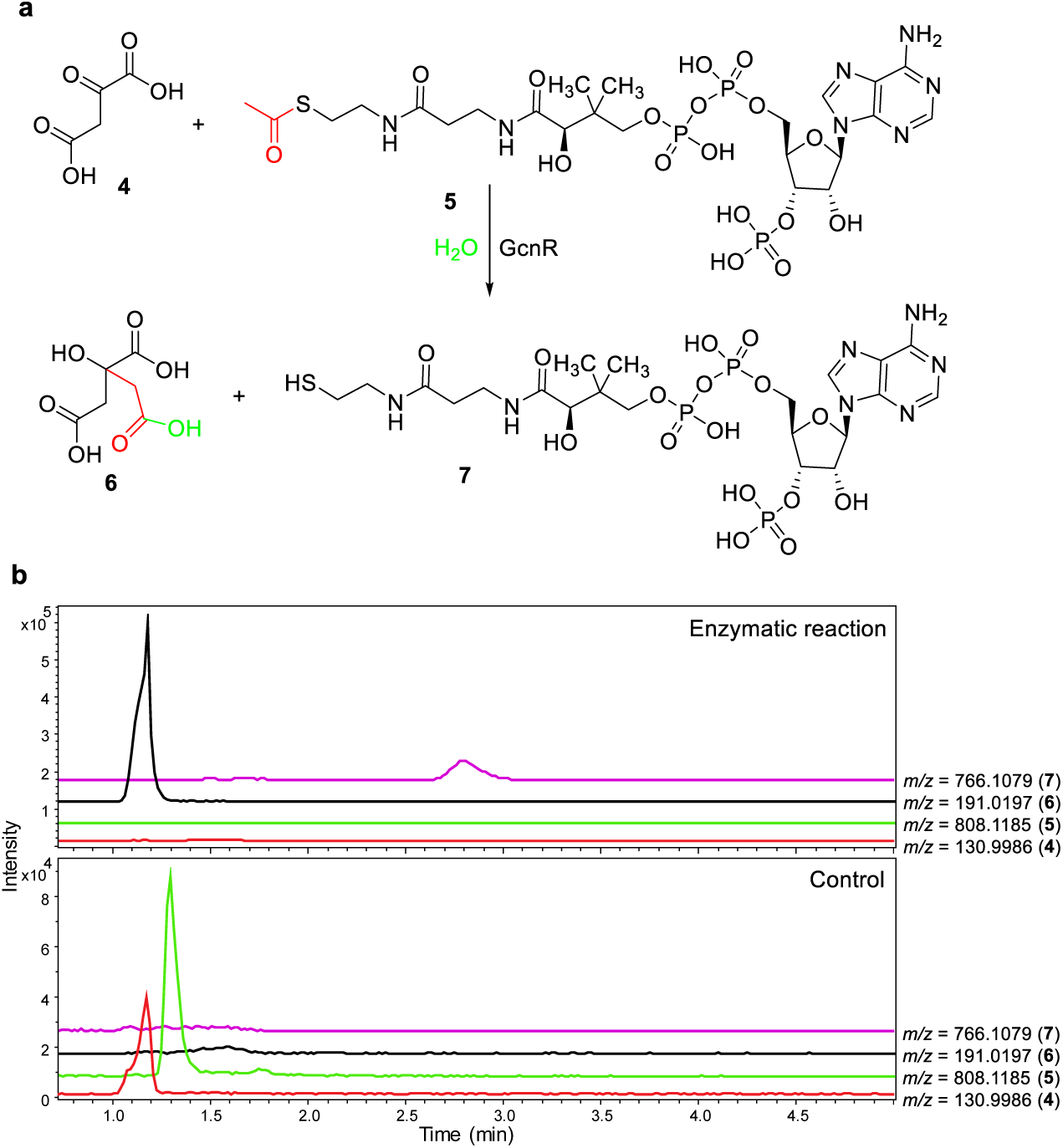
(**a**) Reaction catalyzed by GcnR. (**b)** Extracted ion chromatograms for *m/z* = 130.9986, 808.1185, 191.0197, and 766.1079 corresponding to [M-H]^−^ for **4, 5, 6** and **7** from negative ion mode UHPLC-ESI-Q-TOF-MS analyses of the GcnR-catalyzed condensation of oxaloacetate with acetyl-CoA (top panel) and a negative control reaction containing GcnR inactivated by boiling for 15 minutes prior to addition (bottom panel). The measured *m/z* values for citrate and coenzyme A were 191.0195 and 766.1076, respectively.

To probe the role played by citrate in gladiochelin biosynthesis, we constructed an in-frame deletion in *gcnR*. The resulting mutant was unable to assemble gladiochelins A and B, but produced a new metabolite with the molecular formula C_37_H_59_N_5_O_12_ (calculated for C_37_H_60_N_5_O_12_^+^: 766.4233, found: 766.4238; Figures 6 and S17). The production level of this metabolite was insufficient for NMR spectroscopic analysis. We therefore conducted LC-MS/MS analyses, which indicated the metabolite has the same depsipeptide core as gladiochelin B **2** (molecular formula C_40_H_63_N_5_O_14_; Figures 6 and S17) and that the structural differences must lie in the fatty acid and/or Dba residues. However, because the new metabolite contains three fewer carbon atoms, four fewer hydrogen atoms and two fewer oxygen atoms than gladiochelin B, but the same number of nitrogen atoms, these differences cannot be due to modification or loss of the Dba residue. The most plausible explanation is that malate, another intermediate in the Krebs cycle, is used instead of citrate (albeit inefficiently) to initiate assembly of the fatty acid residue (Figure S18). Malate does not, however, appear to be a substrate for the O-methyltransferase GcnS (see below). We thus tentatively assign structure **3** to the gladiochelin B derivative produced by the *gcnR* mutant.

**Figure 6.**
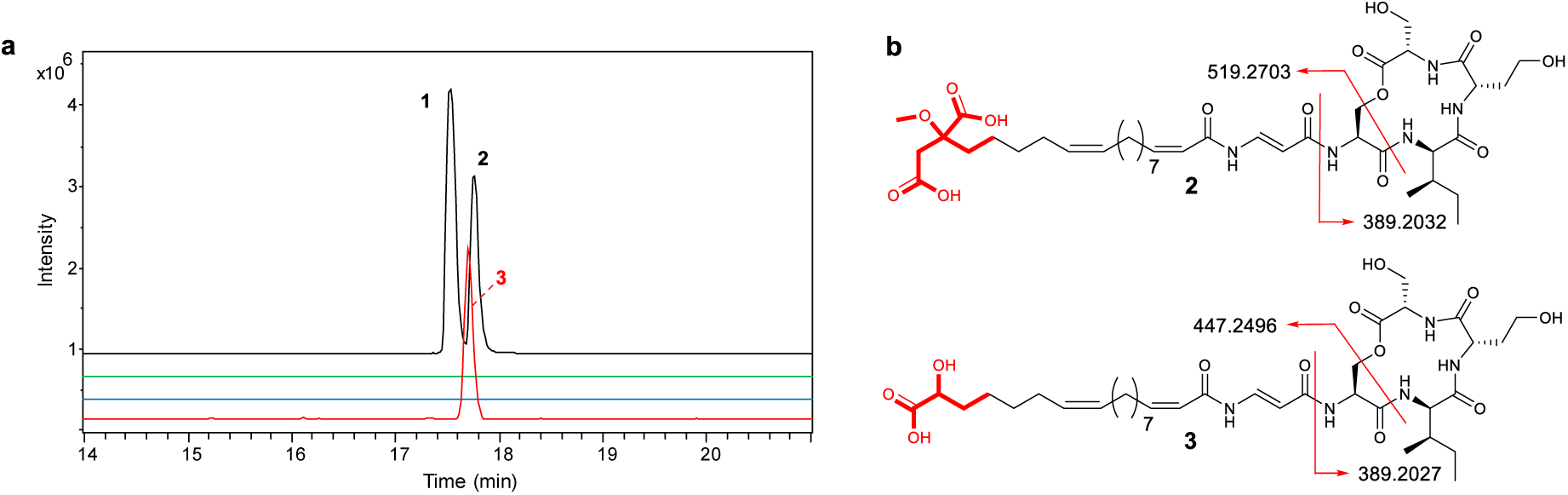
(**a**) Extracted ion chromatograms (EICs) from UHPLC-ESI-Q-TOF-MS analyses of extracts from *B. gladioli* BCC1622 Δ*gbnD1_ER* and a *gcnR* deletion mutant. From top to bottom, these are: EIC at *m/z* 824.4288 (corresponding to [M+H]^+^ for **1**) and 838.4444 (corresponding to [M+H]^+^ for **2**) for the wild type strain; EIC at *m/z* 824.4288 and 838.4444 for the mutant; EIC at *m/z* 766.4233 (corresponding to [M+H]^+^ for **3**) for the wild type strain; and EIC at *m/z* 766.4233 for the mutant. (**b**) Comparison of fragment ions observed for metabolite **3** and gladiochelin B (**2**) in MS/MS analyses.

We postulated that GcnS, which shows sequence similarity to *S*-adenosyl-L-methione (SAM) dependent methyltransferases, catalyzes methylation of the hydroxyl group in the citrate-derived starter unit at some point during gladiochelin biosynthesis. To test this hypothesis, we constructed an in-frame deletion in *gcnS*. The production of gladiochelins A **1** and B **2** was abrogated in the resulting mutant and two new metabolites with the molecular formulae C_38_H_59_N_5_O_14_ (calculated for C_38_H_60_N_5_O_14_^+^: 810.4131, found: 810.4124) and C_39_H_61_N_5_O_14_ (calculated for C_39_H_62_N_5_O_14_^+^: 824.4288, found: 824.4296) were produced (Figures 7 and S17). LC-MS/MS analyses indicated that these compounds are congeners of **1** and **2**, lacking a CH_2_ group from the fatty acid or Dba residues (Figure S17). Although production levels of these metabolites were low, we were able to purify a sufficient quantity of the gladiochelin B congener for NMR spectroscopic analysis (Table S5, Figures S19-S24). Differences were observed in the ^13^C chemical shifts of the resonances due to C-16, C-17, C-18, C-19 and C-20 in the fatty acid residue and the signals attributed to the O-methyl group in gladiochelin B were absent from the ^1^H and ^13^C spectra. We therefore conclude that the two metabolites accumulated in the *gcnS* mutant are desmethyl-gladiochelins A **8** and B **9**. The low levels of **8** and **9** produced by the mutant suggest O-methylation of the citrate-derived starter unit is an early step in gladiochelin biosynthesis, rather than a late-stage modification. However, the precise timing of this reaction remains to be determined.

**Figure 7.**
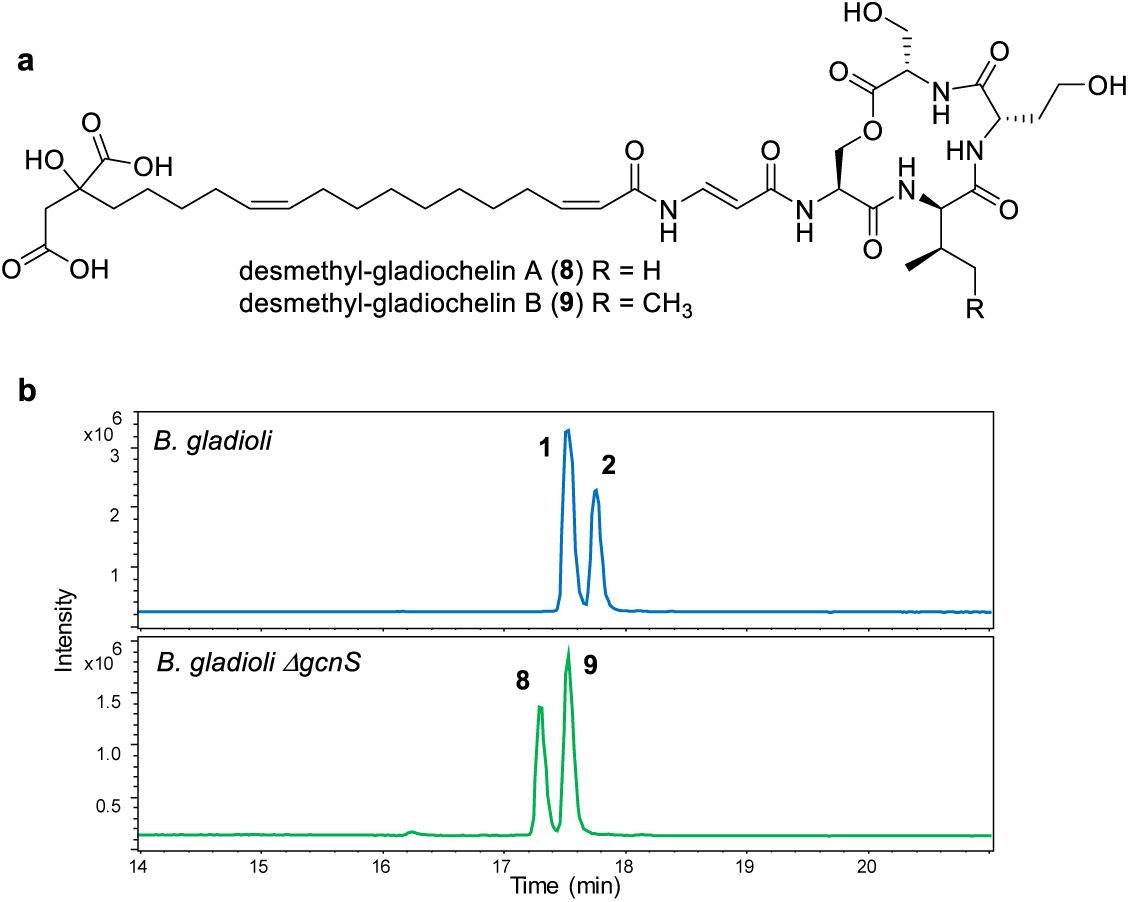
(**a**) Structures of desmethyl-gladiochelins A **8** and B **9** accumulated in the *gcnS* mutant of *B. gladioli* BCC1622 Δ*gbnD1_ER*. (**b**) Top panel: extracted ion chromatogram at *m/z* = 824.4288 (corresponding to [M+H]^+^ for **1**) and 838.4444 (corresponding to [M+H]^+^ for **2**) from UHPLC-ESI-Q-TOF MS analyses of culture extracts from *B. gladioli* BCC1622 Δ*gbnD1_ER*. Bottom panel extracted ion chromatogram at *m/z* = 810.4131 (corresponding to [M+H]^+^ for **8**) and 824.4288 (corresponding to [M+H]^+^ for **9**) from UHPLC-ESI-Q-TOF MS analyses of culture extracts from *gcnS* mutant of *B. gladioli* BCC1622 Δ*gbnD1_ER*.

Citrate (or its O-methylated derivative) is proposed to be converted to the corresponding coenzyme A thioester by GcnB, which shows similarity to acyl-CoA synthetases (Figure 3 and Table 1). Either GcnM or GcnP, both of which show similarity to β-ketoacyl-ACP synthase III (KAS III) enzymes that typically initiate fatty acid biosynthesis in bacteria (Table 1),^45^ could then catalyse the elongation of citryl-CoA (or its O-methylated derivative) with a malonyl group attached to the primary metabolic fatty acid synthase (FAS) ACP (Figure 3). Interestingly, the conserved active site Cys residue is mutated to Ser in GcnM and Thr in GcnP. The functional significance of this is unclear, but an analogous Cys to Ser mutation is observed in DpsC, a KAS III homologue that has been reported to initiate assembly of the daunorubicin polyketide chain using a propionyl-CoA starter unit.^46^ Further processing of the β-ketothioester resulting from elongation of (O-methyl)-citryl-CoA with malonyl-ACP by the primary metabolic FAS would afford the saturated O-methyl-citrate-primed fatty acyl-ACP thioester **10** (Figure 3). The 11, 12 and 2, 3 double bonds are likely introduced into this thioester by GcnF and GcnL, which are similar to membrane-associated fatty acid desaturases and acyl-ACP desaturases, respectively (Figure 3 and Table 1), completing the assembly of the unusual fatty acid residue incorporated into the gladiochelins.

The Dba residue of the gladiochelins is postulated to derive from L-aspartate, which we propose undergoes adenylation of its β-carboxyl group catalyzed by GcnO, followed by transfer onto GcnC, a putative freestanding acyl carrier protein (Figure 3). GcnO shows similarity to VinN,^47^ which catalyzes adenylation of the β-carboxyl group of β-methyl-aspartate and subsequent transfer to a freestanding acyl carrier protein VinL in the biosynthesis of vicenistatin. Moreover, the Asp230 and Ser299/Arg331 residues in VinN, which are proposed to play a key role in recognition of the α-amino and α-carboxyl groups of β-methylaspartate, respectively, are conserved in GcnO (Supplementary Fig. 25). In vicenistatin biosynthesis, the pyridoxal phosphate (PLP)-dependent enzyme VinO catalyses decarboxylation of the β-methyl-*iso*-aspartyl-VinL thioester to form the corresponding α-methyl-β-alanyl thioester.^43^ We propose that GcnN, which shows sequence similarity to BtrK, a PLP-dependent enzyme that catalyzes decarboxylation of an *iso*-glutamyl-ACP thioester in butirosin biosynthesis, plays an analogous role in gladiochelin biosynthesis (Figure 3). In-frame deletion of *gcnN* in *B. gladioli* BCC1622 Δ*gbnD1* abolished gladiochelin production and it could not be restored by feeding β-alanine to the mutant. This is consistent with the proposed roles of GcnO, GcnC and GcnN and rules out β-alanine as an intermediate in the biosynthetic pathway. The amino group of the β-alanyl-GcnC thioester is proposed to be condensed with the citrate-primed fatty acyl thioester by one of the KAS III homologues GcnM or GcnP (Figure 3).

The sequence similarity of GcnQ to acyl-CoA dehydrogenases suggested it might be responsible for formation of the Dba residue via desaturation of the N-acyl-β-alanyl-GcnC thioester. To test this hypothesis, we created an in-frame deletion in *gcnQ*, which abolished the production of gladiochelins A and B and led to the accumulation of two new metabolites with the molecular formulae C_39_H_63_N_5_O_14_ and C_40_H_65_N_5_O_14_ (Figure 7). Purification and NMR spectroscopic analysis showed these metabolites are dihydro-gladiochelins A **11** and B **12**, in which the Dba residue has been replaced by β-alanine (Tables S6 and S7, and Figures S26-S36). This is consistent with the proposed function of GcnQ as a N-acyl-β-alanyl thioester desaturase, although on the basis of these data the possibility of GcnQ converting the β-alanine residue to Dba at a later stage in gladiochelin biosynthesis cannot be ruled out.

### The *gcn* locus is widely conserved in *B. gladioli*

A local nucleotide BLAST search of 1318 genomes representing *Burkholderia, Paraburkholderia* and *Caballeronia* species showed *B. gladioli* is the only species containing the gladiochelin biosynthetic gene cluster (as indicated by the presence of *gcnH*). Read mapping of paired-end Illumina reads from 234 *B. gladioli* strains against the *gcn* gene cluster revealed that it is present in 105 strains. However, there is evidence of gene substitutions and deletions in the regions flanking the cluster (Figure 9). The *gcnU* gene, encoding the putative TonB-dependent ferric-gladiochelin outer membrane receptor, is absent from five of the 105 *B. gladioli* genomes (BCC1837, BCC1843, BCC1861, BCC1870 and BCC1871) containing the gladiochelin biosynthetic gene cluster and is present in 52% (67 of 129) of genomes lacking it.

**Figure 8.**
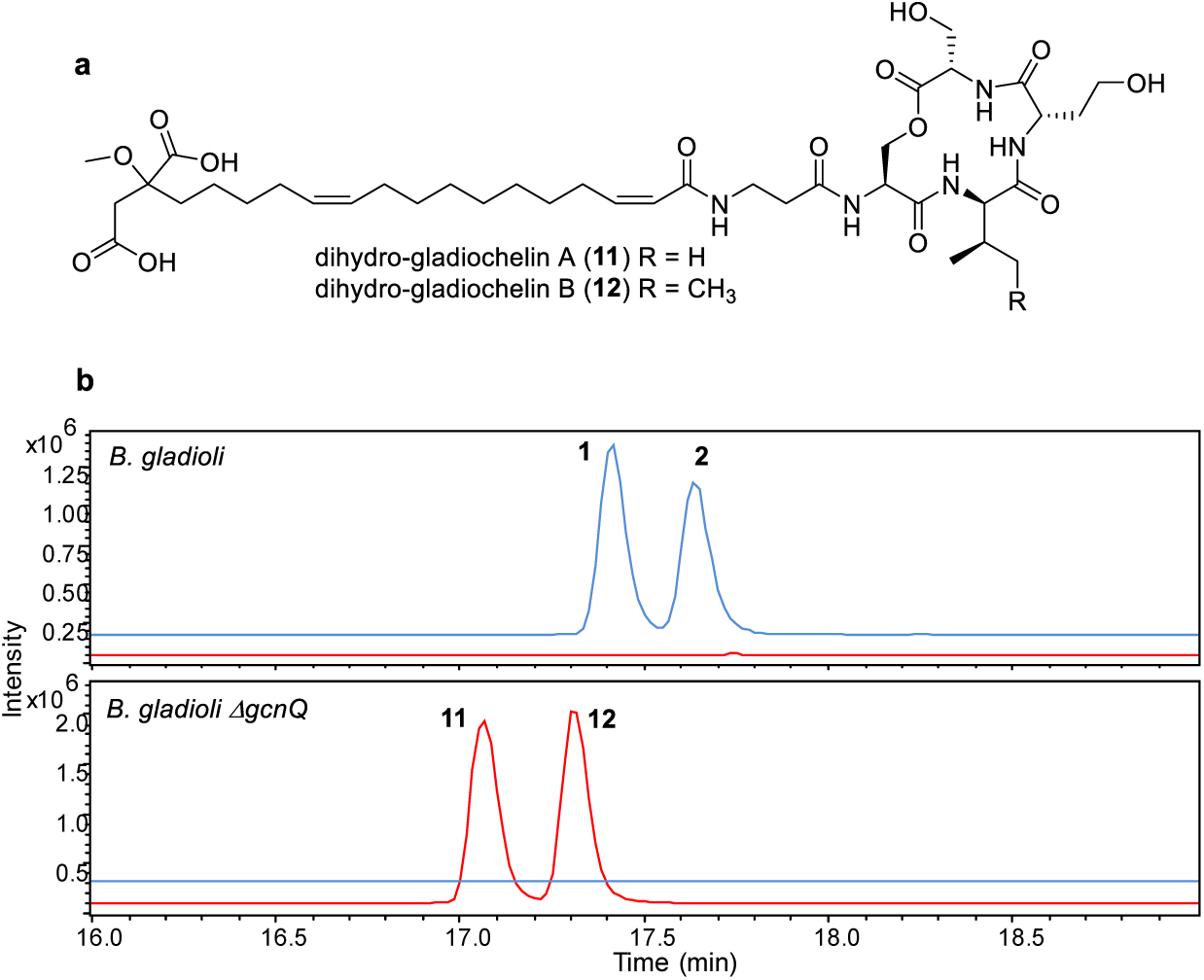
(**a**) Structures of dihydro-gladiochelins A (**11**) and B (**12**) produced by the *gcnQ* mutant of *B. gladioli* BCC1622 Δ*gbnD1_ER*. (**b**) Extracted ion chromatograms at *m/z* = 824.4288 and 838.4444 (in blue; corresponding to [M+H]^+^ for gladiochelins A (**1)** and B (**2**), respectively), and 826.4444 and 840.4601 (in red, corresponding to [M+H]^+^ for dihydro-gladiochelins A (**11**) and B (**12**), respectively) from UHPLC-ESI-Q-TOF MS analyses of culture extracts from *B. gladioli* BCC1622 Δ*gbnD1_ER* and the *gcnQ* mutant.

**Figure 9.**
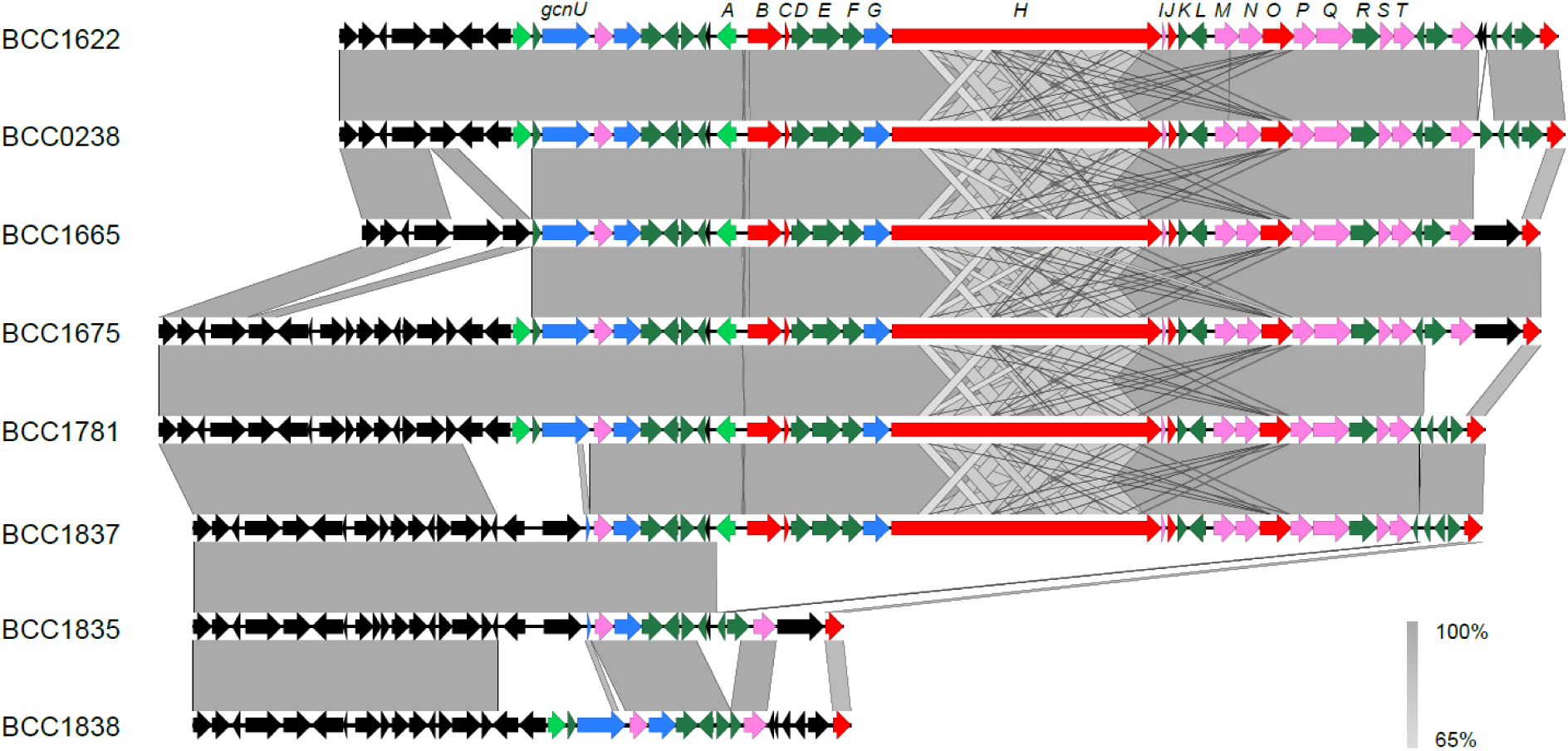
Conservation of the gladiochelin biosynthetic gene cluster in *B. gladioli.* The gene cluster was extracted from six *B. gladioli* genome assemblies (BCC1622, BCC0238, BCC1665, BCC1675, BCC1781, and BCC1837, all of which are CF lung infection isolates) that highlight the diversity of gene insertions, deletions, and substitutions flanking the cluster. *B. gladioli* BCC1835 and BCC1838 were included as examples of strains lacking the gladiochelin biosynthetic gene cluster. No variation in the *gcnA-gcnT* genes was observed for strains containing the gladiochelin biosynthetic gene cluster.

### Biological function of the gladiochelins

Gladiochelins A (**1**) and B (**2**) showed no activity against any of the ESKAPE panel of bacterial pathogens,^48^ *Mycobacterium bovis* BCG, or *Candida albicans*. The 3-methoxy-1, 4-dicarboxylate moiety of the gladiochelins is reminiscent of ferric iron-chelating citrate residues in several siderophores, such as achromobactin, vibrioferrin, and staphyloferrin B.^49^ Moreover, *gcnU* flanking the left end of the gladiochelin biosynthetic gene cluster encodes a putative TonB-dependent ferric-siderophore outer membrane receptor. We thus hypothesised that the gladiochelins function as siderophores. Although ferric complexes of the gladiochelins were to labile to observe in ESI-Q-TOF MS analyses, the chrome azurol S (CAS) assay confirmed they are able to sequester ferric iron (Figure S36).^4^

Siderophore production in microbes is usually regulated by iron availability. In iron-deficient media, production is upregulated, whereas in iron replete media it is suppressed. Accordingly, the level of gladiochelin production by *B. gladioli* BCC1622 Δ*gbnD1_ER* decreased as increasing concentrations of ferric iron were added to the medium (Figure S36). Combined with the results of the CAS assay, these data indicate that the gladiochelins likely function as siderophores in *B. gladioli*.

Siderophores often contribute to virulence in pathogenic bacteria. Thus, we used a *Galleria mellonella* wax moth infection model to investigate whether the gladiochelins are a virulence factor in *B. gladioli*. Ornibactin was shown to contribute to virulence in *Burkholderia cenocepacia* using this and other models.^33, 50^ However, no attenuation of virulence was towards the wax moth larvae was observed in the Δ*gcnN* mutant of *B. gladioli* BCC1622 Δ*gbnD1_ER*, relative to the parental strain (Figure S37).

A similar observation was reported for pyochelin deficient mutants of *B. cenocepacia* H111,^50^ leading to the hypothesis that some *Burkholderia* siderophores may play a role in metal homeostasis rather than virulence.^51^ To investigate this hypothesis, *B. cenocepacia* H111 pyochelin and ornibactin non-producing mutants were grown alongside the wild type strain in media containing various metal salts (10-50 μM). The metal salts did not affect the wild type strain, but were toxic to the siderophore non-producing mutants, indicating that pyochelin and ornibactin protect *B. cenocepacia* against metal ion toxicity.^51^ This finding was further supported by supplementing the medium with pyochelin and ornibactin, which reversed toxicity in the siderophore non-producing mutants.^51^

We thus compared the tolerance of the *B. gladioli* BCC1622 Δ*gbnD1_ER* and Δ*gbnD1_ER-* Δ*gcnN* strains to salts of aluminium, zinc, cobalt, copper, cadmium, nickel and lead at various concentrations up to 100 μM. No decrease in metal ion tolerance was observed for the gladiochelin non-producing mutant relative to parent strain and the results of these experiments indicated that *B. gladioli* is much less susceptible to metal ion toxicity than *B. cenocepacia*. Further experiments demonstrated that *B. gladioli* could readily grow in media containing 1 mM Ni^2+^, Al^2+^ and Zn^2+^. Additional work is required to establish the genetic basis for this high degree of metal ion tolerance.

## Conclusions

Bacteria belonging to the *Burkholderia* genus are increasingly being recognised as an underexplored source of novel natural products.^52^ Using a combination of carbon source modification and inactivation of the biosynthetic pathway for gladiolin, which is produced at high titre and interferes with the detection of lower abundance metabolites, we have identified gladiochelins A (**1**) and (**2**) as the metabolic product of a cryptic nonribosomal peptide biosynthetic gene cluster in *B. gladioli*. Similar approaches may prove useful for identifying the products of other cryptic *Burkholderia* biosynthetic gene clusters.

The gladiochelins have very unusual structures, consisting of a unique fatty acid residue with several polar functional groups appended to its tail, linked via a very rare Dba residue to a depsitetrapeptide. Identification of the gladiochelin biosynthetic gene cluster in *B. gladioli* BCC0238 and BCC1622 enabled us to probe the origin of these unusual structural features using a combination of detailed bioinformatics analysis, gene deletions and enzyme activity assays. Based on these studies, we propose that the fatty acid residue originates from a citryl-CoA starter unit. To our knowledge, there is no precedent for the utilisation of citryl-CoA as a starter unit in fatty acid or polyketide biosynthesis. We also hypothesise that the Dba residue derives from loading of aspartate onto a freestanding ACP. The resulting *iso*-aspartyl thioester is proposed to undergo decarboxylation and N-acylation, prior to being desaturated by a Flavin-dependent dehydrogenase, resulting in an N-acyl-Dba thioester starter unit for the NRPS that assembles the depsitetrapeptide. While loading of aspartate / β-methyl-aspartate onto a standalone ACP, followed by decarboxylation and N-acylation, is a well-established mechanism for provision of β-aminoacyl starter units to type I modular polyketide synthases,^53^ to our knowledge there is no precedent for this mechanism being employed to provide such starter units to NRPSs. Thus, our findings expand the scope of β-aminoacyl starter unit biosynthetic machinery, which may prove useful in bioengineering approaches to natural product diversification.

The gladiochelin biosynthetic gene cluster is present in approximately 45% of *B. gladioli* genomes, but was not found in other *Burkholderia* species. This suggests the gladiochelins play an important role in the adaption of *B. gladioli* to its environmental niches. Several lines of evidence indicate that gladiochelin functions as a siderophore, challenging the assumption that *B. gladioli* employs siderophore-independent mechanisms for iron acquisition.^33^ While the mode of ferric iron binding to gladiochelin remains to be established, it seems likely that the two citrate-derived carboxy and methoxy groups in the unusual fatty acid residue, and one or both of the side hydroxyl groups in the depsipetide are involved. Siderophores are known to play roles in virulence and metal ion tolerance in other *Burkholderia* species, but a *Galleria mellonella* infection model did not provide evidence that the gladiochelins contribute to virulence in *B. gladioli* and gladiochelin-deficient mutants did not show reduced tolerance towards metal ions. Further studies are therefore required to develop a better understanding of the adaptive benefit the gladiochelins confer on *B. gladioli*.

## Acknowledgments

This research was supported by grants from the BBSRC (BB/L021692/1 and BB/L023342/1) to E.M, G.L.C, T.R.C and J.P; A.J.M., G.W., E.M. and G.L.C. also acknowledge current funding from BBSRC grants BB/S007652/1 and BB/S008020/1. Y.D. was supported by a grant from the MRC (MR/N501839/1 to G.L.C). I.T.N. was supported by the BBSRC through the Midlands Integrative Bioscience Doctoral Training Partnership (BB/M01116X/1). E.M. and G.W. acknowledge funding from the Life Science Bridging Fund LSBF R2-004 for the HPLC analysis and instrumentation. The Bruker MaXis Impact UHPLC-ESI-Q-TOF-MS instrument used in this research was funded by the BBSRC (B/K002341/1 to G.L.C.). G.L.C. was the recipient of a Wolfson Research Merit Award from the Royal Society (WM130033)

