## Supporting Information for "Discovery and biosynthesis of gladiochelins: unusual lipodepsipeptide siderophores from *Burkholderia gladioli*"


| **Contents** | **Page** |
|---|---|
| Materials and methods | S3-S12 |
| Table S1. PCR primers used to create in-frame deletion and insertional inactivation constructs | S13 |
| Figure S1. Structure of gladiochelin A defining the nomenclature and numbering used in Table S2. | S14 |
| Figure S2. Structure of gladiochelin B defining the nomenclature and numbering used in Table S3. | S14 |
| Table S2. $^{1}$H, $^{13}$C and HMBC NMR data for gladiochelin A in pyridine-$d_5$. | S15 |
| Table S3. $^{1}$H, $^{13}$C and HMBC NMR data for gladiochelin B in pyridine-$d_5$. | S16 |
| Figures S3-S8. $^{1}$H NMR, $^{13}$C NMR, COSY, HSQC, HMBC, and ROESY spectra of gladiochelin A in pyridine-$d_5$. | S17-S22 |
| Figure S9-S14. $^{1}$H NMR, $^{13}$C NMR, COSY, HSQC, HMBC, and ROESY spectra of gladiochelin B in pyridine-$d_5$. | S23-S28 |
| Figure S15. LC-MS comparison of Marfey's derivatives of gladiochelin A and B hydrolysates with authentic standards. | S29 |
| Table S4: Predicted specificity pocket residues and substrates of the GcnH adenylation domains. | S30 |
| Figure S16. MS/MS spectra and proposed fragment ions for gladiochelins A and B, metabolite **3**, and desmethyl-gladiochelins A and B. | S31 |
| Figure S17. Comparison of the proposed biosynthetic pathways for galdiochelin B (**2**) and compound **3** produced by the Δ*gcnR* mutant. | S32 |
| Figure S18. Structure of desmethyl-gladiochelin B summarising COSY and key HMBC correlations, and defining the nomenclature and numbering used in Table S5. | S33 |
| Table S5. $^{1}$H, $^{13}$C and HMBC NMR data for desmethyl-gladiochelin B in pyridine-$d_5$. | S34 |
| Figure S19-S23. $^{1}$H NMR, $^{13}$C NMR, COSY, HSQC, and HMBC spectra of desmethyl-gladiochelin B in pyridine-$d_5$. | S35-S39 |
| Figure S24. Sequence alignment of VinN and GcnO. | S40 |
| Figure S25. Structure of dihydro-gladiochelins A and B summarising COSY and key HMBC correlations, and defining the nomenclature and numbering used in Tables S6 and S7. | S40 |
| Table S6. $^{1}$H, $^{13}$C and HMBC NMR data for dihydro-gladiochelin A in pyridine-$d_5$. | S41 |
| Table S7. $^{1}$H, $^{13}$C and HMBC NMR data for dihydro-gladiochelin B in pyridine-$d_5$. | S42 |
| Figures S26-S30. $^{1}$H NMR, $^{13}$C NMR, COSY, HSQC, and HMBC spectra of dihydro-gladiochelin A in pyridine-$d_5$. | S43-S47 |

| | |
|---|---|
| Figures S31-S35. $^{1}$H NMR, $^{13}$C NMR, COSY, HSQC, and HMBC spectra of dihydro-gladiochelin B in pyridine-$d_5$. | S48-S52 |
| Figure S36. Results of CAS assay with gladiochelin A and LC-MS chromatograms showing gladiochelin production is suppressed by addition of ferric iron to the growth medium. | S53 |
| Figure S37. Results of virulence assay of *B. gladioli* BCC1622 and *B. gladioli* BCC1622 Δ*gbnD1_ER1*-Δ*gcnN* using the *Galleria* wax moth larvae model. | S53 |

### Materials and methods

**General experimental procedures:** A Bruker Alpha Platinum ATR single reflection diamond ATR module was used to acquire IR spectra and a Perkin Elmer Lambda 35 UV/vis spectrophotometer was used to record UV/vis spectra. For optical rotations measurement, an Optical Activity Ltd AA-1000 millidegree auto-ranging polarimeter (589 nm) was used. Specific rotations are given in units of $10^{-1}$ deg $cm^2$ $g^{-1}$. UHPLC-ESI-Q-TOF-MS analyses were performed using a Dionex UltiMate 3000 UHPLC connected to a Zorbax Eclipse Plus C18 column (100 × 2.1 mm, 1.8 μm) coupled to a Bruker MaXis IMPACT mass spectrometer. Mobile phases consisted of water (A) and acetonitrile (B), each supplemented with 0.1% formic acid. A gradient of 5% B to 100% B over 30 minutes was employed at a flow rate of 0.2 mL/min. The mass spectrometer was operated in positive ion mode with a scan range of 50-3000 *m/z*. Source conditions were as follows: end plate offset at −500 V; capillary at −4500 V; nebulizer gas ($N_2$) at 1.6 bar; dry gas ($N_2$) at 8 L min−1; dry temperature at 180 °C. Ion transfer conditions were as follows: ion funnel RF at 200 Vpp; multiple RF at 200 Vpp; quadrupole low mass at 55 m/z; collision energy at 5.0 eV; collision RF at 600 Vpp; ion cooler RF at 50–350 Vpp; transfer time at 121 μs; pre-pulse storage time at 1 μs. Calibration was performed with 1 mM sodium formate through a loop injection of 20μL at the start of each run. Analytical HPLC-ESI-Ion Trap-MS analyses were performed on a Bruker amaZon X connected to a ZORBAX 300SB-C3 column (150 × 4.6 mm, 5 μm). A gradient of 15% to 100% acetonitrile over 50 minutes was employed at a flow rate of 1 mL/min. NMR spectra of the gladiochelins and derivative were recorded in pyridine-$d_5$. NMR spectra were recorded on a Bruker 500 MHz spectrometer equipped with a DUI cryoprobe at 25 °C. The $^1$H and $^{13}$C NMR chemical shifts were referenced to the solvent peaks at $\delta_H$ 7.58 and $\delta_C$ 135.91 for pyridine-$d_5$. All HPLC and LC-MS experiments were performed with MeCN-$H_2O$ gradient solvent system. HPLC grade solvents were used for chromatography.

**Production, extraction and HPLC purification of gladiochelins A and B:** *B. gladioli* BCC0238 Δ*gbnD1_ER1* was grown on 2 L solid Basal Salts Medium (BSM),[1] containing glycerol (4 g/L) and ribose (4 g/L) as carbon sources, for 3 days at 30 °C. The agar was extracted with ethyl acetate and the resulting solution was evaporated to dryness. The residue was resuspended in 1.5 mL of methanol, pre-adsorbed to C18-bonded silica and packed into a stainless steel HPLC guard cartridge (10 × 30 mm), which was connected to a semi-preparative reverse-phase C18 Betasil column (21.2 mm × 150 mm). The column was eluted as follows: 5% MeCN / 95% $H_2O$ for 5 min; linear gradient from 5 to 100% MeCN over 45 min; 100% MeCN for 10 min. The flow rate was 9 mL/min. 120 fractions of equal volume were collected over 60 min. Gladiochelins A and B were obtained as amorphous solids by evaporation of fractions 61 and 63, respectively.

*Gladiochelin A*: (1.5 mg from 2 L culture media); $[\alpha]_D^{28}$ -17 (*c* 0.05, MeOH); UV (MeOH) $\lambda_{max}$ (log ε) 227 (4.98), 288 (5.26) nm; IR $\nu_{max}$ 3284, 2918, 2849, 1735, 1670, 1642, 1602, 1551, 1461, 1409, 1239, 1187, 1059, 724 $cm^{-1}$; $^1H$ NMR (500 MHz, pyridine-$d_5$) and $^{13}C$ NMR (125 MHz, pyridine-$d_5$), see Table S2; HRESIMS *m/z* 824.4285 $[M + H]^+$ (calcd for $C_{39}H_{62}N_5O_{14}$, 824.4288).

*Gladiochelin B*: (1.7 mg from 2 L culture media); $[\alpha]_D^{28}$ -10 (*c* 0.05, MeOH); UV (MeOH) $\lambda_{max}$ (log ε) 227 (4.97), 285 (5.22) nm; IR $\nu_{max}$ 3278, 2917, 2849, 1736, 1669, 1642, 1602, 1459, 1403, 1239, 1189, 1061, 716 $cm^{-1}$; $^1H$ NMR (500 MHz, pyridine-$d_5$) and $^{13}C$ NMR (125 MHz, pyridine-$d_5$), see Table S3; HRESIMS *m/z* 838.4445 $[M + H]^+$ (calcd for $C_{40}H_{64}N_5O_{14}$, 838.4444).

**Identification and *in silico* analysis of the gladiochelin biosynthetic gene cluster:** antiSMASH v3.0[2] was used to identify putative specialised metabolite biosynthetic gene clusters in the genomes of *Burkholderia gladioli* BCC0238 and BCC1622. The functions of proteins encoded by genes in the gladiochelin biosynthetic gene cluster were assigned via comparative sequence analyses. The substrate specificity of the A domains in GcnH were predicted as described (Table S4).[3, 4] The specificity conferring residues for Hse-activating A domains was determined by analysing the A domain in module 4 of the nunapeptin synthetase NunD.[5] The substrate specificity of GcnO was assigned by comparing its sequence with the β-methyl-*iso*-aspartyl-ACP synthetase VinN involved in vicenistatin biosynthesis.[6]

**Genetic manipulation of *B. gladioli* strains:** The insertional inactivation and in-frame deletion mutants were created using a pGPI-*Sce*I-based homologous recombination mutagenesis system.[7] Plasmids were mobilised into *B. gladioli* by tri-parental mating.

#### *Tri-parental mating procedures*

The parental *B. gladioli* strain was grown in 5 mL of LB medium (supplemented with 150 μg/mL of trimethoprim (Tp) when the strain is a single crossover mutant) at 30 °C overnight. Donor *E. coli* SY327, carrying the pGPI-*Sce*I insertional inactivation/in-frame deletion construct or pDAI-*Sce*I, and helper *E. coli* HB101, carrying pRK2013, were grown in 5mL of LB medium supplemented with the appropriate antibiotic at 37 °C overnight (pGPI-*Sce*I constructs: Tp 50 μg/mL, pDAI-*Sce*I: tetracycline (Tc) 20 μg/mL, pRK2013: kanamycin (Km) 50 μg/mL). The overnight cultures of *B. gladioli* and *E. coli* donor/helper strains were centrifuged, and the cell pellets were washed with LB medium and resuspended in 5 mL of LB medium containing 10 mM $MgCl_2$. 100 μL of each re-suspended mixture was combined and 100 μL of the resulting mixture was spread onto a nitrocellulose membrane (Millipore, pore size 0.22 μm) placed on a LB agar plate. After overnight incubation at 30 °C, the cells were washed off the membrane with 1mL of sterile 0.9% NaCl and 100 μL of the mixture was spread on an LB agar plate containing appropriate antibiotics (for donor *E. coli* strain carrying pGPI-*Sce*I constructs: 600 U/mL polymyxin B (Pm) and 150 μg/mL Tp; for the donor *E. coli* strain carrying pDAI-*Sce*I: 600 U/mL Pm and 200 μg/mL Tc). The plate was incubated at 30 °C for 72_h, and colonies were picked and grown in LB medium containing the same antibiotics as the agar plates to confirm the antibiotic resistance phenotype. Colonies with the correct phenotype were screened for the desired genotype using colony PCR.

***Construction of gladiolin non-producing mutants of* B. gladioli *BCC0238 and* B. gladioli *BCC1622***

Because our originally reported gladiolin non-producing mutant of *B. gladioli* BCC0238 contains and insertion in *gbn*D1,[8] it was not suitable for further genetic manipulation using the same mutagenesis system. Thus, we created a mutant of *B. gladioli* BCC0238 with an in-frame deletion in *gbnD1*, which was used as the parental strain for subsequent genetic modifications. To do this, regions of *gbnD1* flanking the region encoding the ER domain (see table S1 for the sequences of the PCR primers used) were ligated into pGPI-*Sce*I via *Xba*I and *Eco*RI restriction sites, and the resulting construct (pGPI-*gbnD1_ER1*) was used to transform *E. coli* SY327, then mobilized into *B. gladioli* BCC0238 using tri-parental mating, as described above. The $Tp^R$ phenotype of exconjugants was confirmed by growing them in LB medium containing Tp (150 μg/mL) at 30 °C overnight and colony PCR was used to screen for the desired single crossover mutants. One correct mutant was transformed with pDAI-*Sce*I by tri-parental mating. pDAI-SceI constitutively expresses the I-*Sce*I nuclease, which creates double strand breaks at the *Sce*I site in the backbone of pGPI-*Sce*I constructs, necessitating a second crossover to repair the chromosome. Exconjugants with a $Tc^RTp^S$ phenotype were anticipated to have the desired genotype. However, the introduction of pDAI-*Sce*I into the *B. gladioli* BCC0238 single crossover mutant was found to be very inefficient. Only a few colonies were obtained, all of which were $Tc^RTp^R$. This made the creation of in-frame deletions in *B. gladioli* BCC0238 very challenging using this mutagenesis system. Therefore, we switched to a different approach.

The *B. gladioli* BCC0238 *gbnD1_ER1* single crossover mutant was grown without antibiotic selection on a LB agar plate for several generations to promote a second crossover and colonies were screened for $Tp^S$. Sensitive colonies were further screened using colony PCR to identify those with the desired genotype. DNA sequencing was used to further confirm the genotype of one of the mutants and it was named *B. gladioli* BCC0238 Δ*gbnD1_ER1*.

Because the construction of in-frame deletions in *B. gladioli* BCC0238 proved to be challenging, we identified *B. gladioli* BCC1622 as an alternative producer of gladiochelin (and gladiolin) that is amenable to genetic manipulation using the pGPI-*Sce*I / pDAI-*Sce*I-based mutagenesis system. We therefore introduced the *gbnD1_ER1* deletion into *B. gladioli* BCC1622. The pGPI-*gbnD1_ER1* construct used to create the single crossover mutant of *B. gladioli* BCC0238 was introduced into *B. gladioli* BCC1622 using the procedure described above. After introducing pDAI-*Sce*I into the resulting single crossover mutant of *B. gladioli* BCC1622 by tri-parental mating, $Tc^R Tp^S$ exconjugants were selected and screened for the Δ*gbnD1_ER1* genotype using colony PCR. The resulting mutants were grown on an M9 agar plate at 30 °C for 72 hours to promote loss of the pDAI-*Sce*I plasmid, allowing the construction of double gene deletion mutants. Colonies were picked and screened for $Tc^S Tp^S$ (indicative of pDAI-*Sce*I loss). DNA sequencing was used to further confirm the genotype of one of the resulting mutants and it was named *B. gladioli* BCC1622 Δ*gbnD1_ER1*.

***Insertional inactivation of gcnH in* B. gladioli *BCC0238* ΔgbnD1_ER1**

The involvement of the *gcn* locus in gladiochelin biosynthesis was initially investigated via insertional inactivation of *gcn*H in *B. gladioli* BCC0238 Δ*gbnD1_ER1*. A 930 bp internal fragment of *gcnH* was cloned into pGPI-*Sce*I via *Eco*RI and *Xba*I restriction sites (the sequences of the PCR primers used are shown in Table S1), and the resulting construct was used to transform *E. coli* SY327, then mobilized into *B. gladioli* BCC0238 Δ*gbnD1_ER1* by tri-parental mating, as described above. Exconjugants were confirmed to be $Tp^R$ and screened for the correct genotype using colony PCR. One correct mutant was selected and named *B. gladioli* BCC0238 Δ*gbnD1_ER1* Ω*gcnH.*

***Construction of in-frame deletions in gladiochelin biosynthetic genes in* B. gladioli *BCC1622* ΔgbnD1_ER1**

In-frame deletions in *gcnR, gcnS, gcnN, gcnQ* and *gcnT* were constructed by cloning regions flanking the targeted regions into pGPI-*Sce*I via *Xba*I and *Eco*I restriction sites (the sequences of the PCR primers used are shown in Table S1). The resulting constructs were used to transform *E. coli* SY327 and mobilized into *B. gladioli* BCC1622 Δ*gbnD1_ER1* by triparental mating. The remaining steps in construction and confirmation of the mutants were as described above for construction of the *B. gladioli* BCC1622 Δ*gbnD1_ER1* mutant.

**Purification of desmethyl-gladiochelin B, dihydro-gladiochelin A, and dihydro-gladiochelin B:** The culture conditions and HPLC method used for purification of desmethyl-gladiochelin B, dihydro-gladiochelin A, and dihydro-gladiochelin B were similar to the those used for gladiochelins A and B. Desmethyl-gladiochelin B, dihydro-gladiochelin A, and dihydro-gladiochelin B eluted in fractions 64, 61, and 63, respectively.

*Desmethyl-gladiochelin B*: amorphous solid (0.5 mg from 8 L culture media); $[\alpha]_D^{30}$ -14 (*c* 0.05, MeOH); UV (MeOH) $\lambda_{max}$ (log ε) 228 (4.97), 287 (5.25) nm; IR $\nu_{max}$ 3276, 2918, 2850, 1736, 1670, 1642, 1601, 1458, 1403, 1238, 1188, 1060 $cm^{-1}$; $^1H$ NMR (500 MHz, pyridine-$d_5$) and $^{13}C$ NMR (125 MHz, pyridine-$d_5$), see Table S5; HRESIMS *m/z* 824.4293 $[M + H]^+$ (calcd for $C_{39}H_{62}N_5O_{14}$, 824.4288).

*dihydro-gladiochelin A*: amorphous solid (1.1 mg from 4 L culture media); $[\alpha]_D^{32}$ -16 (*c* 0.05, MeOH); UV (MeOH) $\lambda_{max}$ (log ε) 234 (3.39) nm; IR $\nu_{max}$ 3314, 2927, 2854, 1733, 1655, 1534, 1458, 1376, 1234, 1195, 1058 $cm^{-1}$; $^1H$ NMR (500 MHz, pyridine-$d_5$) and $^{13}C$

NMR (125 MHz, pyridine-$d_5$), see Table S6; HRESIMS $m/z$ 826.4447 $[M + H]^+$ (calcd for $C_{39}H_{64}N_5O_{14}$, 826.4444).

*dihydro-gladiochelin B*: amorphous solid (1.4 mg from 4 L culture media); $[\alpha]_D^{32}$ -15 ($c$ 0.05, MeOH); UV (MeOH) $\lambda_{max}$ (log $\varepsilon$) 235 (3.39) nm; IR $\nu_{max}$ 3314, 2929, 2854, 1728, 1713, 1658, 1537, 1457, 1378, 1227, 1060 cm$^{-1}$; $^1$H NMR (500 MHz, pyridine-$d_5$) and $^{13}$C NMR (125 MHz, pyridine-$d_5$), see Table S7; HRESIMS $m/z$ 840.4602 $[M + H]^+$ (calcd for $C_{40}H_{66}N_5O_{14}$, 840.4601).

**Overproduction and purification of GcnR:** *gcnR* was amplified from genomic DNA of *Burkholderia gladioli* BCC1622 using Phusion DNA polymerase (NEB) with the following primers; CitSyn-F: (5′-CAGCATATGATGGAGACGAACCCCAGCAGCACG-3′) and CitSyn-R: (5′-GACAAGCTTTTACAGCCAGCCCGCCGGCAC-3′). Following separation of PCR products on a 1% agarose gel, the desired bands were excised and purified with a GeneJET Gel Extraction Kit (Thermo Scientific). The insert was digested with *Nde*I and *Hin*dIII and ligated to *Nde*I and *Hin*dIII-digested pET28a using T4 DNA ligase. The resulting vector was used to transform chemically competent *E. coli* TOP10 cells (Invitrogen) and was plated on LB agar containing Km (50 μg/mL). Colonies were picked and grown overnight in LB medium supplemented with Km (50 μg/mL). Plasmids were extracted using a GeneJET Plasmid Miniprep Kit (Thermo Scientific), digested with *Nco*I and analysed by agarose gel electrophoresis to identify those containing the correct insert. The inserts of positive clones were sequenced to verify their integrity. One correct clone was used to transform *E. coli* BL21(DE3) and a single transformant was inoculated into LB medium containing Km (50 μg/mL) and incubated overnight at 37 °C and 180 rpm. The resulting culture was used to inoculate 2 L of LB medium containing Km (50 μg/mL), which was incubated at 37 °C and 180 rpm until the $OD_{595}$ of the culture reached 0.6. IPTG (0.5 mM) was added and the culture was incubated overnight at 15 °C and 180 rpm. Cells were harvested by centrifugation (4,500 rpm, 20 min, 4 °C), resuspended in 20 mM Tris-HCl, 100 mM NaCl, pH 7.8 at 20 mL/L of growth medium, and lysed by sonication. The lysate was centrifuged (17,000 rpm, 60 min, 4 °C) and the supernatant was passed through a 0.45 μm filter, then loaded onto a HiTrap Chelating Column (GE Healthcare), which had been equilibrated with the lysis buffer. Proteins were eluted in a stepwise manner using increasing concentrations of imidazole (50, 100, 200, and 300 mM) in lysis buffer. SDS-PAGE was used to identify fractions containing recombinant GcnR. These fractions were combined and concentrated to ~100 μM using a 10 kDa MWCO Vivaspin centrifugal concentrator (Sartorius). Aliquots of 50 μL were snap-frozen in liquid $N_2$ and stored at -80 °C.

**Citrate Synthase Assay:** A 100 μL of reaction mixture containing oxaloacetic acid (100 μM), Acetyl-CoA (100 μM), and purified recombinant GcnR (50 μM) in Tris-HCl (20 mM, pH 7.8) was incubated at room temperature for 2 hours. The reaction was stopped by adding 100 μL of methanol and the precipitated protein was pelleted by centrifugation. The supernatant was analysed by negative ion mode UHPLC-ESI-Q-TOF-HRMS on a Zorbax Eclipse Plus C18 column (100 × 2.1 mm, 1.8 μm) at a flow rate of 0.2 mL/min. The column was eluted with a combination of 0.1% aqueous ammonia solution (eluent A) and acetonitrile (eluent B) using the following profile: from 100% A to 95% A / 5% B over 20 minutes; from 95% A / 5% B to 100% B over three minutes; isocratic 100% B. The negative control contained GcnR that had been inactivated by incubating at 100 °C for 15 min before addition to the reaction mixture.

**Analysis of the distribution of the gladiochelin biosynthetic gene cluster across *Burkholderia* and related genera:** A local nucleotide BLAST search v2.7.1+ was performed to screen for the presence of the gladiochelin biosynthetic gene cluster in 1318 genomes representing *Burkholderia*, *Paraburkholderia* and *Caballeronia* species. The presence/absence and completeness of the gene cluster in *B. gladioli* genomes was assessed by a read mapping approach. Paired-end Illumina reads associated with *B. gladioli* and subordinate taxa were downloaded from the European Nucleotide Archive. These reads were mapped against the *B. gladioli* BCC1622 gladiochelin biosynthetic gene cluster using Snippy v3.2-dev (https://github.com/tseemann/snippy). The coverage per base of the *gcn* gene cluster was extracted with bedtools and visualised in a read coverage plot. The gladiochelin biosynthetic gene clusters in different strains were visually compared using Easyfig v2.1.[9]

**Antimicrobial activity assays:** The antimicrobial activity of gladiochelins A and B were tested against the ESKAPE pathogens,[10] *Mycobacterium bovis* BCG and *Candida albicans* using disc diffusion assays. The ESKAPE pathogens, *M. bovis* and *C. albicans* were grown on Mueller-Hinton, 7H10 and YPD agar plates, respectively. 5 μL of a 64 μg/mL DMSO solution of each gladiochelin was placed on a filter paper disc. After allowing the discs to dry, they were placed on the agar plates, which were incubated at 37 °C for 24 h for ESKAPE pathogens and *Candida albicans*, and 72 h for *Mycobacterium bovis* BCG. No zones of growth inhibition around the discs were observed.

***Galleria mellonella* virulence assay:** Infection of *G. mellonella* larvae (TruLarv; BioSystems Technology Ltd., UK) was performed as described previously.[11] Briefly, bacterial cultures of *B. gladioli* BCC1622 Δ*gbnD1_ER1* and BCC1622 Δ*gbnD1_ER1* Ω*gcnH* were grown overnight in TSB at 37°C, washed and resuspended in phosphate-buffered saline PBS, and adjusted to approximately $1 \times 10^5$ colony forming units (CFU) $mL^{-1}$. Larvae were injected with 10 μL aliquots of bacterial suspension or PBS (as a negative control). Each treatment included ten larvae, and the experiment was performed in triplicate over a period of 5 days. All treated larvae were incubated separately at 37°C for 72 h, and the percentage survival was recorded at 18, 21, 24, 42, 45, 48, 66 and 72 h post-inoculation. Total viable counts for the bacterial inoculum were calculated after each experiment by drop-count and the mean inoculum viability ranged from 4.2 to 5.4 × $10^2$ CFU per larvae across the replicate experiments.

**Metal toxicity assays:** To test the protective effect of gladiochelin against different metals, *B. gladioli* BCC1622 Δ*gbnD1_ER1* and BCC1622 Δ*gbnD1_ER1* Ω*gcnH* were grown in BSM containing 4 g/L glycerol (BSMG) supplemented with 0, 6.2, 12.5, 25, 50 and 100 μM metals ($AlCl_2$, $ZnCl_2$, $CoCl_2.6H_20$, $CuCl_2.2H_2O$, $CdCl_2$, $NiCl_2.6H_2O$, $PbCl_2$) in a 96 well plate. After 24 h incubation at 30$^o$C, growth was measured by absorbance at 600 nm. Additional high exposure experiments were carried out in BSMG supplemented with 1 mM metal chlorides.

**Table S1.** PCR Primers used to create in-frame deletion and insertional inactivation constructs.

| Target gene | Parental strain | Primers |
|---|---|---|
| Region of *gbnD1* encoding ER domain | *B. gladioli* BCC0238/ *B. gladioli* BCC1622 | 5′-flank_Forward: CTGTCTAGATTGTCGTCCTCGTCGC<br>5′-flank_Reverse: CAGAAGCTTGAGCGCGTGCACGGTA<br>3′-flank_Forward: GCAAAGCTTGTGGTGATCCGCCATCG<br>3′-flank_Reverse: GCCGAATTCCACGCCATGCACGGGA |
| *gcnH* | *B. gladioli* BCC0238 Δ*gbnD1_ER1* | Forward: ACGTCTAGACACGGCATCCTCGATCAGGC<br>Reverse: CTAGAATTCACCGATCTCACGCGCGAC |
| *gcnR* | *B. gladioli* 1622 Δ*gbnD1_ER1* | 5′-flank_Forward: GACTCTAGACTCACGCCGATCTGCAAG<br>5′-flank_Reverse: GACAAGCTTCCGGCCGGTCATGTTCCG<br>3′-flank_Forward: GACAAGCTTGCCTTCGGCGTCAACAGC<br>3′-flank_Reverse: CTAGAATTCCAGCCTGCCTTCGACCAG |
| *gcnS* | *B. gladioli* 1622 Δ*gbnD1_ER1* | 5′-flank_Forward: GCATCTAGAGAGCAGACCATGGACGCG<br>5′-flank_Reverse: GACAAGCTTCCGGCCGATGAACATCAG<br>3′-flank_Forward: GACAAGCTTCCCTACGCGACCCTCTTC<br>3′-flank_Reverse: CTAGAATTCCGGGTTGGGATGCTGGGC |
| *gcnN* | *B. gladioli* 1622 Δ*gbnD1_ER1* | 5′-flank_Forward: GACTCTAGAGATCGCGGCGTGCAATGC<br>5′-flank_Reverse: CAGAAGCTTCCAGGCATGGTCGGCATC<br>3′-flank_Forward: GACAAGCTTTTCCTGTCGCACCAGCCC<br>3′-flank_Reverse: GCAGAATTCCGCCTCGCAGTATTTCGG |
| *gcnQ* | *B. gladioli* 1622 Δ*gbnD1_ER1* | 5′-flank_Forward: GACTCTAGAGTGGCCAGTTGCGCGCTC<br>5′-flank_Reverse: CGAAAGCTTCTGCTCGGTCTCGAACAG<br>3′-flank_Forward: CAGAAGCTTGGCGACGAACTTGCACAG<br>3′-flank_Reverse: CTAGAATTCCGCGTCCATGGTCTGCTC |
| *gcnT* | *B. gladioli* 1622 Δ*gbnD1_ER1* | 5′-flank_Forward: GACTCTAGAACCTGAGACCGGCGAACG<br>5′-flank_Reverse: GACAAGCTTCGCGCATTGCACGGCATC<br>3′-flank_Forward: CAGAAGCTTCATCGCGACACGCAGGTG<br>3′-flank_Reverse: CTAGAATTCCGAATAGGCGTCCCAGGC |

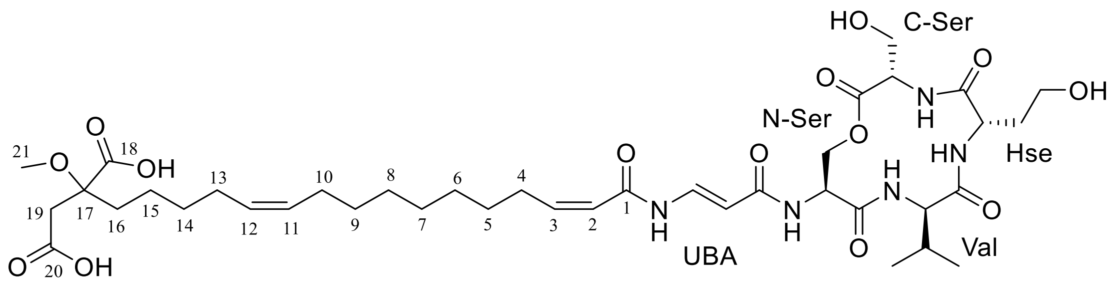


**Figure S1.** Structure of gladiochelin A defining the nomenclature and numbering used in Table S2.

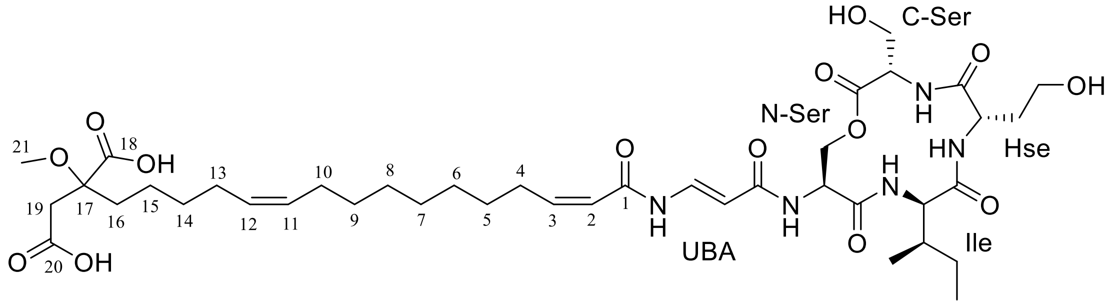


**Figure S2.** Structure of gladiochelin B defining the nomenclature and numbering used in Table S3.

**Table S2.** $^{1}H$, $^{13}C$ and HMBC NMR data for gladiochelin A in pyridine-$d_5$.

| Position | | $\delta_C$ | $\delta_H$ | HMBC |
|---|---|---|---|---|
| C-Ser | C=O | 170.03 | | |
| | α-C | 56.51 | 4.17 (m) | C-Ser(C=O) |
| | β-C | 62.26 | 4.24 (ddd, *J* = 26.8, 10.8, 3.8 Hz) | C-Ser(α-C, C=O) |
| | NH | | 8.17 (d, *J* = 6.9 Hz) | C-Ser(α-C, C=O), Hse (C=O) |
| Hse | C=O | 172.47 | - | |
| | α-C | 53.22 | 5.36 (m) | Hse (C=O, β-C, γ-C) |
| | β-C | 35.13 | 2.22 (m)/2.51 (m) | Hse (C=O, α-C, γ-C) |
| | γ-C | 59.18 | 4.11 (m) | Hse (α-C, β-C) |
| | NH | | 9.91 (d, *J* = 7.5 Hz) | Hse (α-C, β-C), Val (C=O) |
| Val | C=O | 173.93 | - | |
| | α-C | 61.09 | 4.79 (t, *J* = 10.0 Hz) | Val (C=O, β-C, γ-C), Ser2(C=O) |
| | β-C | 29.40 | 2.26 (m) | Val (C=O, α-C, γ-C) |
| | γ-C | 19.33<br>19.81 | 0.97 (d, *J* = 6.6 Hz)<br>0.91 (d, *J* = 6.5 Hz) | Val (α-C, β-C, γ-C)<br>Val (α-C, β-C, γ-C) |
| | NH | | 8.70 (*ol* with solvent peak) | Ser2(C=O) |
| N-Ser | C=O | 171.65 | - | |
| | α-C | 54.13 | 5.52 (m) | N-Ser(C=O), UBA (C=O) |
| | β-C | 67.78 | 4.73 (d, *J* = 10.5 Hz)<br>4.90 (dd, *J* = 10.8, 3.7 Hz) | N-Ser(C=O, α-C), C-Ser(C=O) |
| | NH | | 9.99 (d, *J* = 7.0 Hz) | N-Ser(α-C, β-C), UBA (C=O) |
| UBA | C=O | 168.33 | - | |
| | α-C | 103.95 | 6.00 (d, *J* = 13.9 Hz) | UBA (C=O, β-C) |
| | β-C | 136.80 | 8.83 (dd, *J* = 13.5, 11.4 Hz) | UBA (C=O) |
| | NH | | 11.56 (d, *J* = 11.2 Hz) | UBA (α-C), FA (C=O) |
| Fatty acid (FA) | 1 | 165.14 | - | |
| | 2 | 122.10 | 6.07 (d, *J* = 11.3 Hz) | FA (C=O, C3, C4) |
| | 3 | 150.13 | 6.20 (m) | FA (C=O, C2, C5) |
| | 4 | 29.81 | 2.95 (dd, *J* = 14.6, 7.3 Hz) | FA (C2, C3, C5) |
| | 5 | 29.86 | 1.42 (m) | FA (C3, C4) |
| | 6 | 29.84-30.01 | 1.22-1.32 | |
| | 7 | 29.84-30.01 | 1.22-1.32 | |
| | 8 | 29.84-30.01 | 1.22-1.32 | |
| | 9 | 30.33 | 1.22-1.32 (m) | |
| | 10 | 27.85 | 2.03 (dd, *J* = 13.3, 6.8 Hz) | FA (C9, C11) |
| | 11 | 130.72 | 5.43 (m) | FA (C13) |
| | 12 | 130.32 | 5.43 (m) | FA (C10) |
| | 13 | 27.88 | 2.12 (m) | FA (C12, C14, C15) |
| | 14 | 30.83 | 1.48 (m) | FA (C12, C13, C15, C16) |
| | 15 | 24.15 | 1.77 (m) | FA (C14, C16) |
| | 16 | 35.94 | 2.42 | FA (C14, C15, C17, C18, C19) |
| | 17 | 82.0 | - | |
| | 18 | 175.79 | - | |
| | 19 | 40.69 | 3.41 (d, *J* = 15.1)/ 3.47 (d, *J* = 15.1) | FA (C16, C17, C18, C20) |
| | 20 | 173.51 | - | |
| | 21 | 52.34 | 3.66 (s) | FA (C17) |

**Table S3.** Summary of $^{1}$H, $^{13}$C and HMBC NMR data for gladiochelin B in pyridine-$d_5$.

| Position | | $\delta_C$ | $\delta_H$ | HMBC |
|---|---|---|---|---|
| C-Ser | C=O | 170.0 | - | |
| | α-C | 56.50 | 4.14 (m) | |
| | β-C | 62.20 | 4.25 (ddd, *J* = 24.1, 10.9, 3.9 Hz) | C-Ser(C=O) |
| | NH | | 8.12 (d, *J* = 7.0 Hz) | C-Ser(α-C, C=O), Hse (C=O) |
| Hse | C=O | 172.42 | - | |
| | α-C | 53.15 | 5.38 (m) | Hse (C=O, β-C, γ-C) |
| | β-C | 35.18 | 2.23 (m)/2.55 (m) | Hse (C=O, α-C, γ-C) |
| | γ-C | 59.23 | 4.12 (m) | Hse (α-C, β-C) |
| | NH | | 9.99 (d, *J* = 7.4 Hz) | Hse (α-C, β-C), Ile (C=O) |
| Ile | C=O | 174.06 | - | |
| | α-C | 59.47 | 4.91 (*ol*) | Ile (C=O, β-C, γ1-C, γ2-C), Ser2(C=O) |
| | β-C | 34.93 | 2.12 (m) | Ile (γ1-C) |
| | γ1-C | 25.42 | 1.18 (m)/1.70 (m) | Ile (α-C, β-C, γ2-C, δ-C) |
| | γ2-C | 15.98 | 0.89 (d, *J* = 6.7 Hz) | Ile (α-C, β-C, γ1-C) |
| | δ-C | 10.69 | 0.66 (t, *J* = 7.4 Hz) | Ile (β-C, γ1-C) |
| | NH | | 8.83 (*ol*) | Ser2(C=O) |
| N-Ser | C=O | 171.68 | - | |
| | α-C | 53.99 | 5.57 (m) | N-Ser(C=O), UBA (C=O) |
| | β-C | 67.99 | 4.71 (d, *J* = 10.7 Hz)/4.93 (*ol*) | N-Ser(C=O, α-C), C-Ser(C=O) |
| | NH | | 10.04 (d, *J* = 7.3 Hz) | UBA (C=O) |
| UBA | C=O | 168.13 | - | |
| | α-C | 104.07 | 5.98 (d, *J* = 13.9 Hz) | UBA (C=O, β-C) |
| | β-C | 136.68 | 8.79 (*ol*) | UBA (C=O), FA (C=O) |
| | NH | | 11.54 (d, *J* = 11.2 Hz) | UBA (α-C), FA (C=O) |
| Fatty acid (FA) | 1 | 165.13 | - | |
| | 2 | 122.11 | 6.03 (d, *J* = 11.2 Hz) | FA (C=O, C4) |
| | 3 | 150.06 | 6.18 (m) | FA (C=O, C2, C5) |
| | 4 | 29.81 | 2.95 (dd, *J* = 14.6, 7.3 Hz) | FA (C2, C3, C5) |
| | 5 | 29.86 | 1.42 (m) | FA (C3, C4) |
| | 6 | 29.84-30.01 | 1.23-1.32 | |
| | 7 | 29.84-30.01 | 1.23-1.32 | |
| | 8 | 29.84-30.01 | 1.23-1.32 | |
| | 9 | 30.34 | 1.29 (m) | |
| | 10 | 27.86 | 2.03 (dd, *J* = 13.3, 6.8 Hz) | FA (C11, C12) |
| | 11 | 130.72 | 5.43 (m) | |
| | 12 | 130.33 | 5.47 (m) | |
| | 13 | 27.89 | 2.12 (m) | FA (C11, C12, C14, C15) |
| | 14 | 30.84 | 1.48 (m) | FA (C12, C13, C15, C16) |
| | 15 | 24.16 | 1.77 (m) | FA (C13, C14, C16, C17) |
| | 16 | 35.94 | 2.42 (m) | FA (C14, C15, C17, C18, C19) |
| | 17 | 82.0 | - | |
| | 18 | 175.82 | - | |
| | 19 | 40.71 | 3.41 (d, *J* = 15.1)/ 3.47 (d, *J* = 15.1) | FA (C16, C17, C18, C20) |
| | 20 | 173.53 | - | |
| | 21 | 52.33 | 3.66 (s) | FA (C17) |

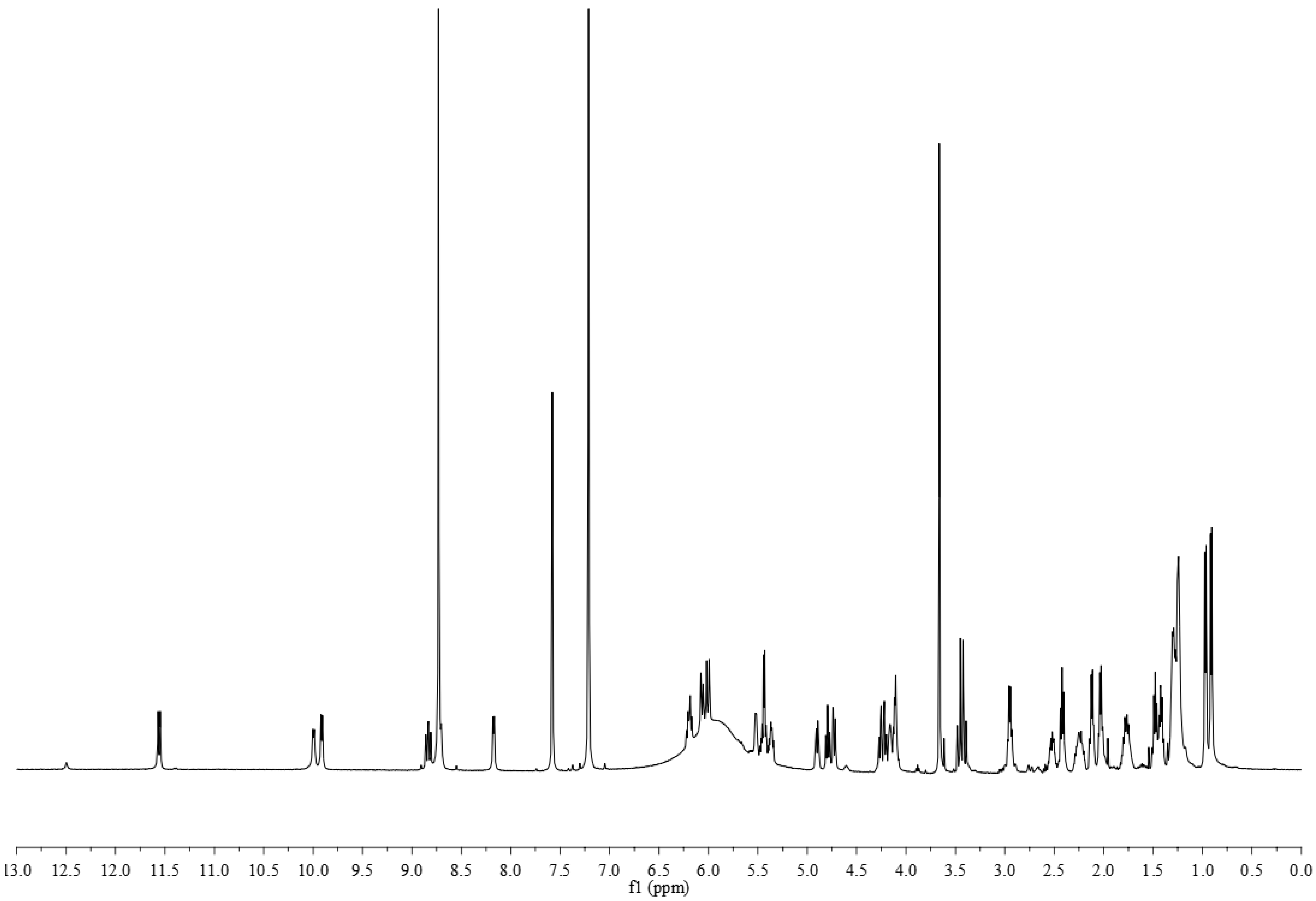


**Figure S3.** $^{1}$H NMR spectrum of gladiochelin A in pyridine-$d_5$.

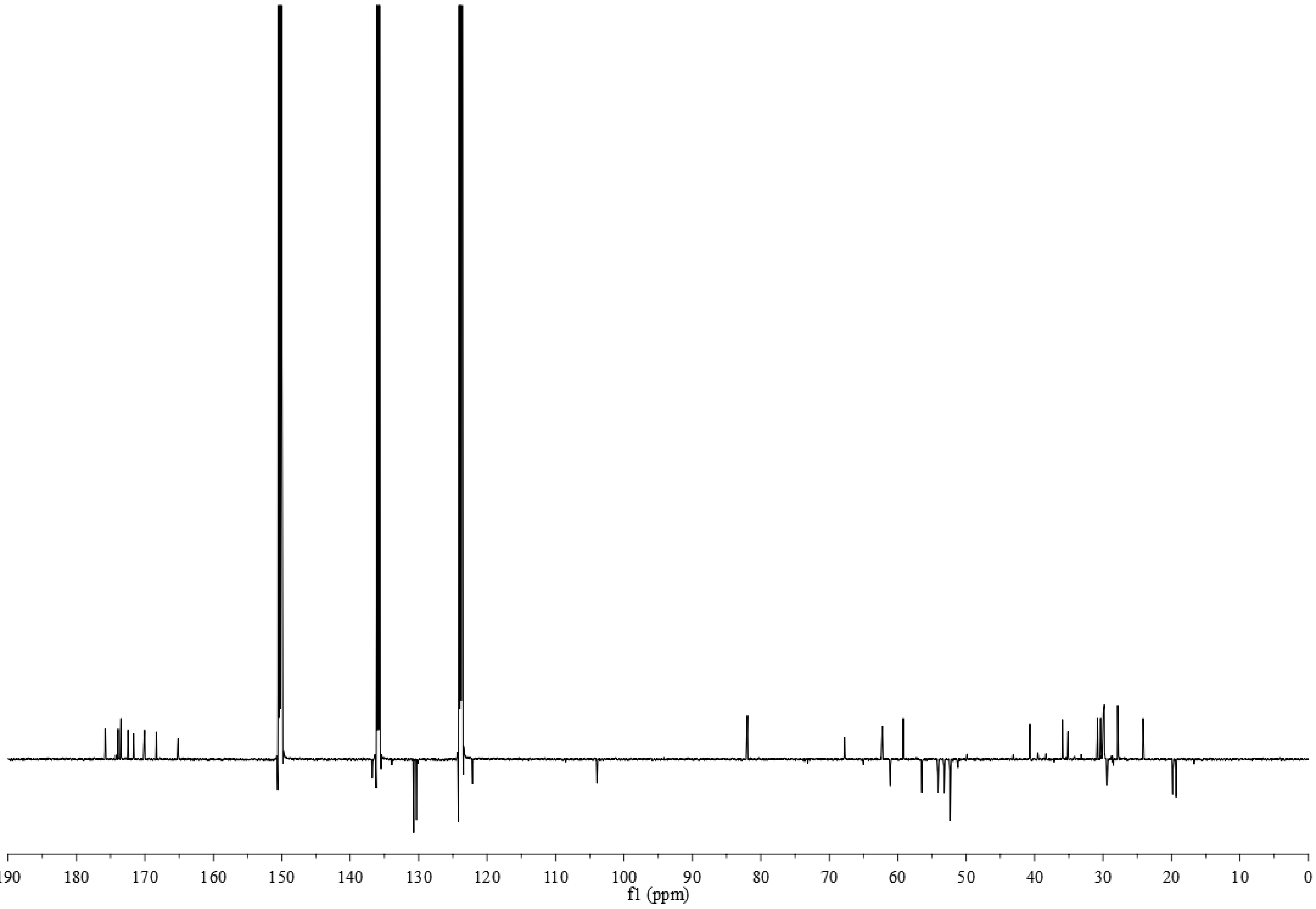


**Figure S4.** $^{13}$C NMR spectrum of gladiochelin A in pyridine-$d_5$.

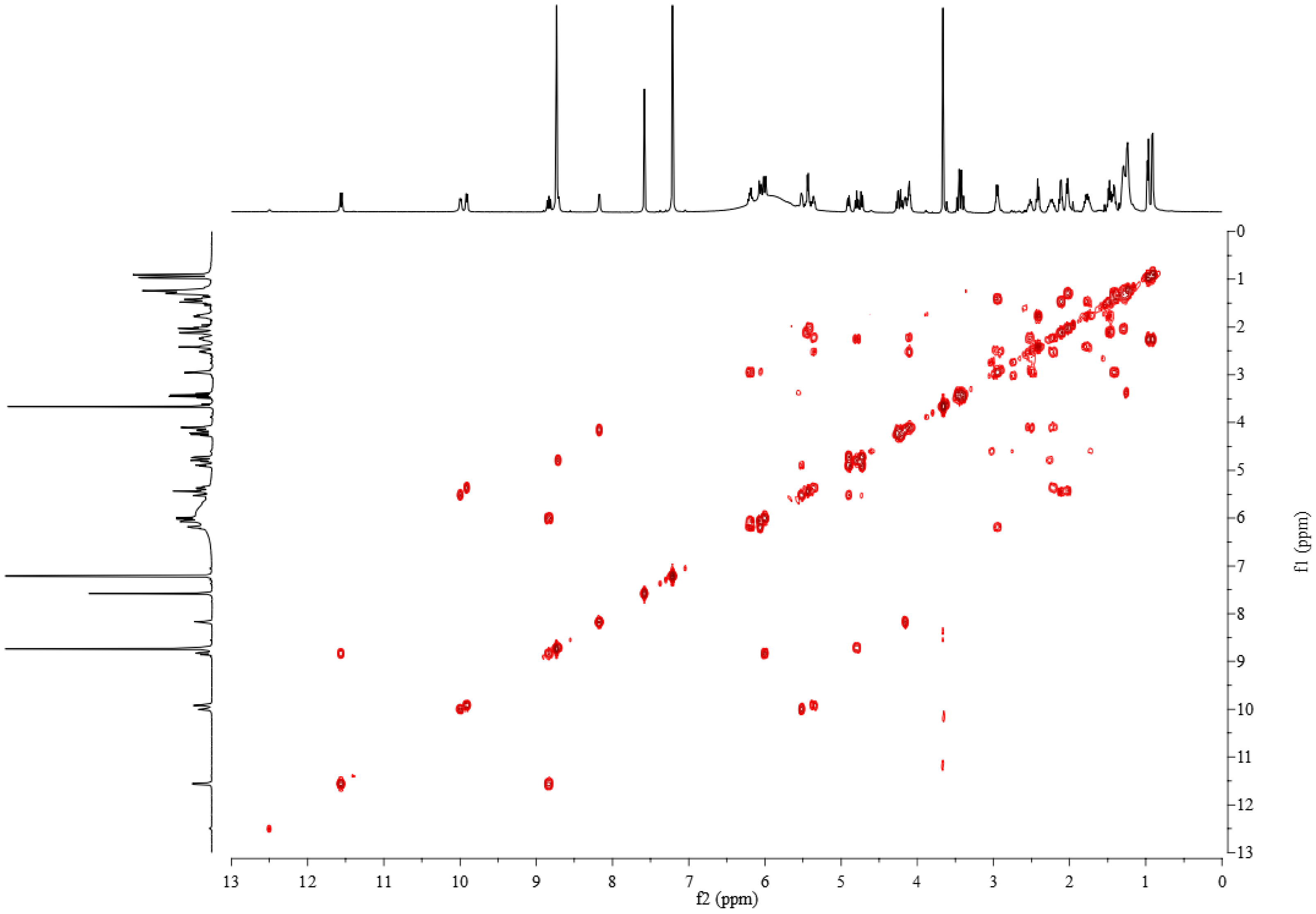


**Figure S5.** COSY spectrum of gladiochelin A in pyridine-$d_5$.

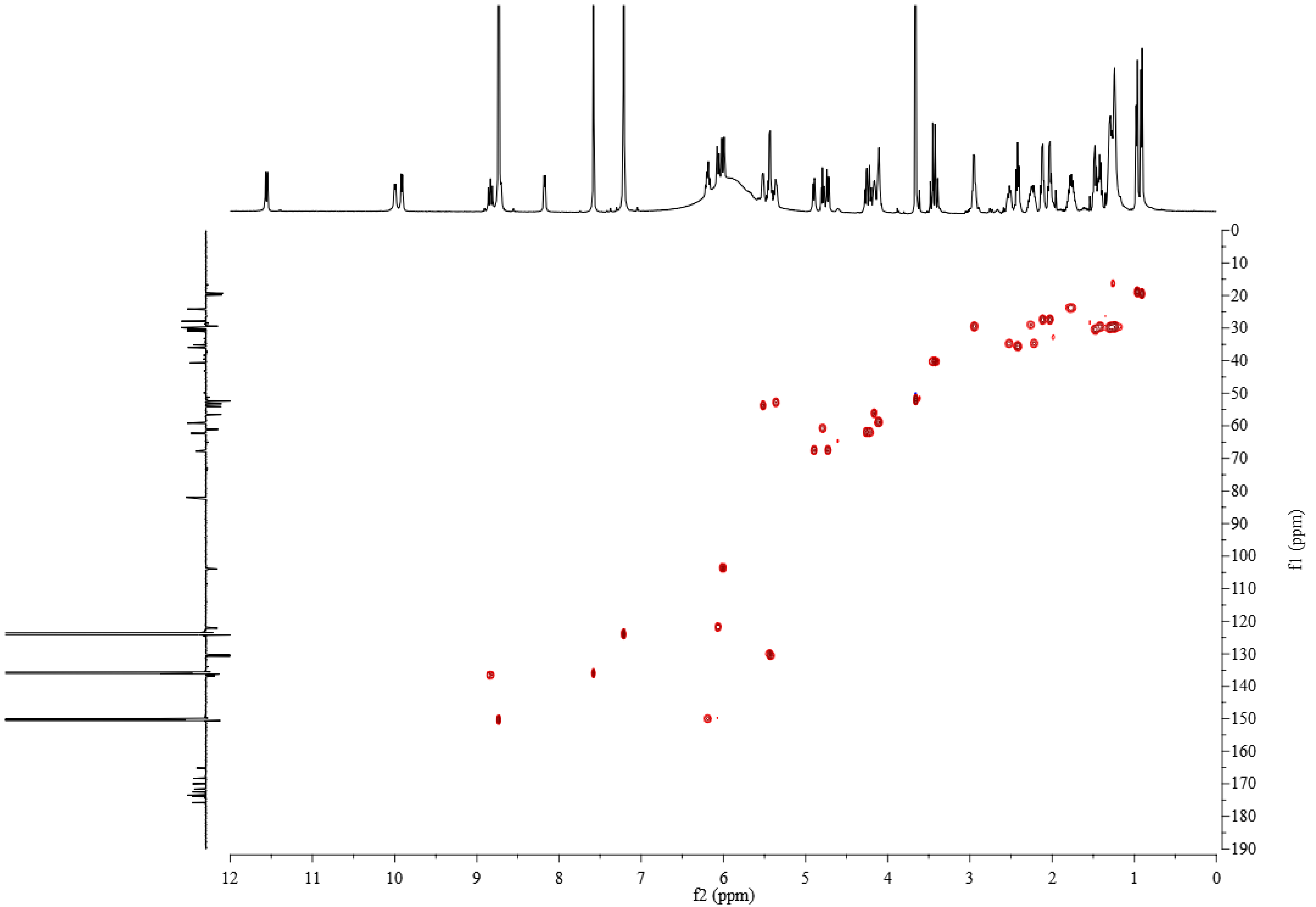


**Figure S6.** HSQC spectrum of gladiochelin A in pyridine-$d_5$.

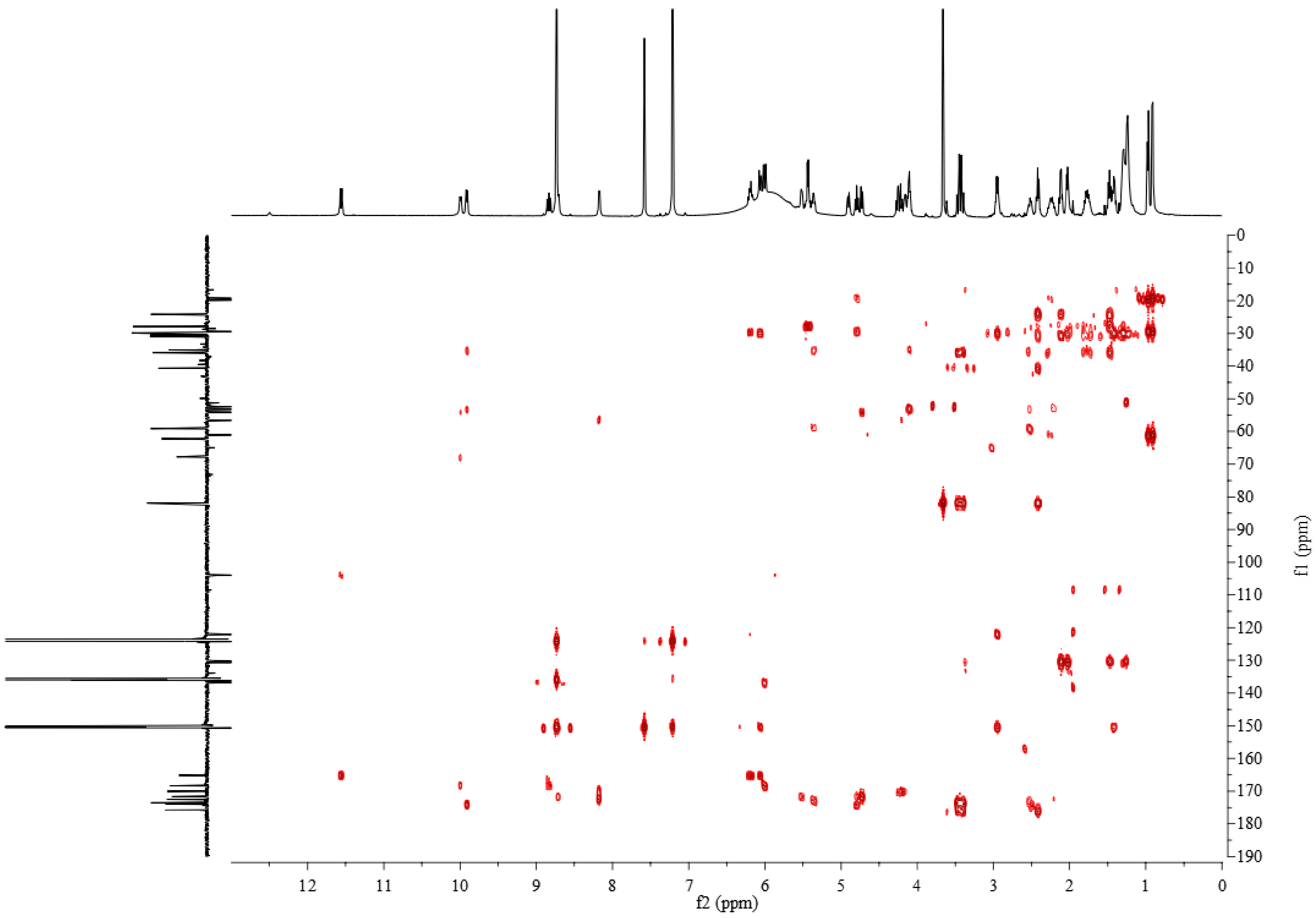


**Figure S7.** HMBC spectrum of gladiochelin A in pyridine-$d_5$.

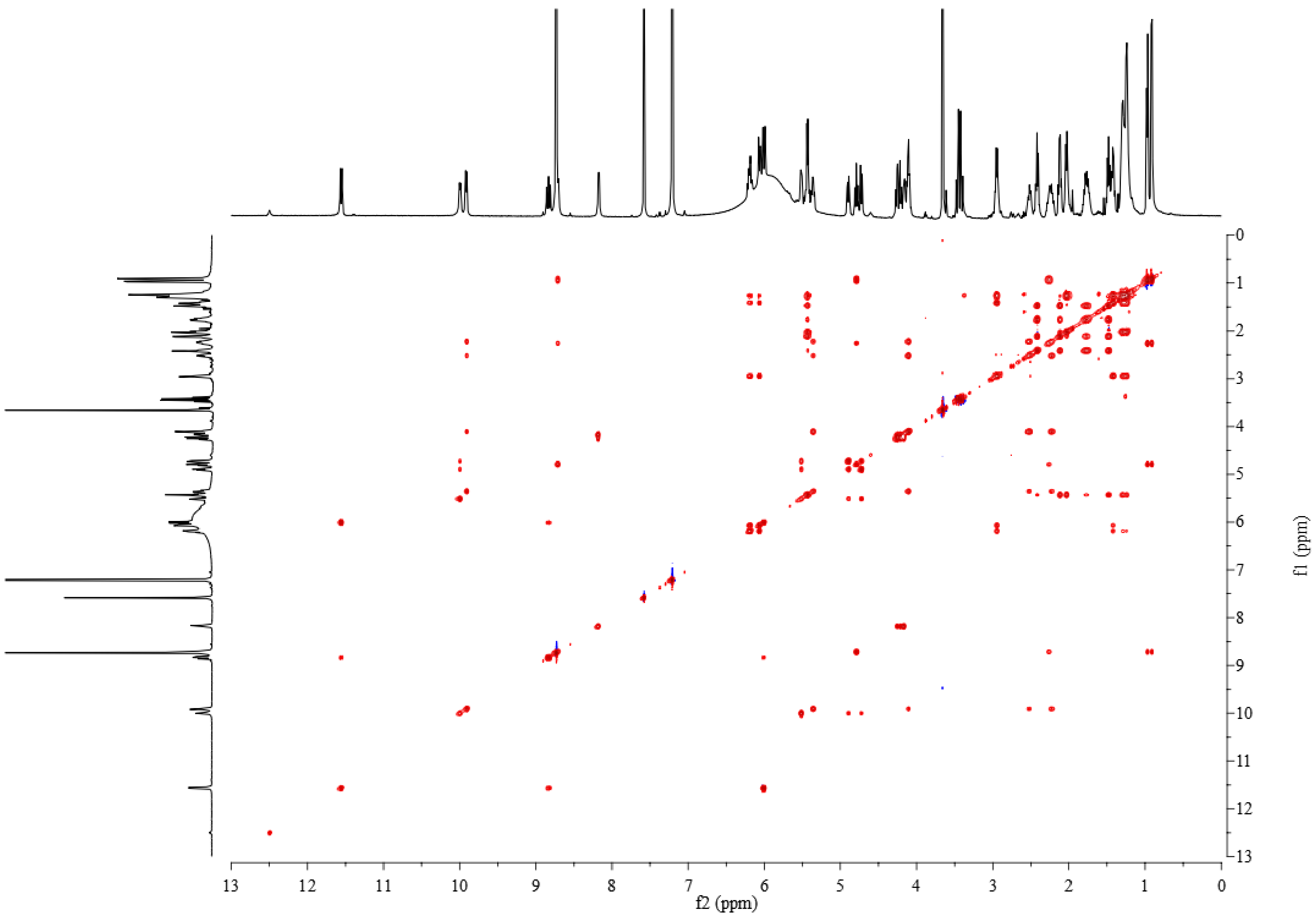


**Figure S8.** ROESY spectrum of gladiochelin A in pyridine-$d_5$.

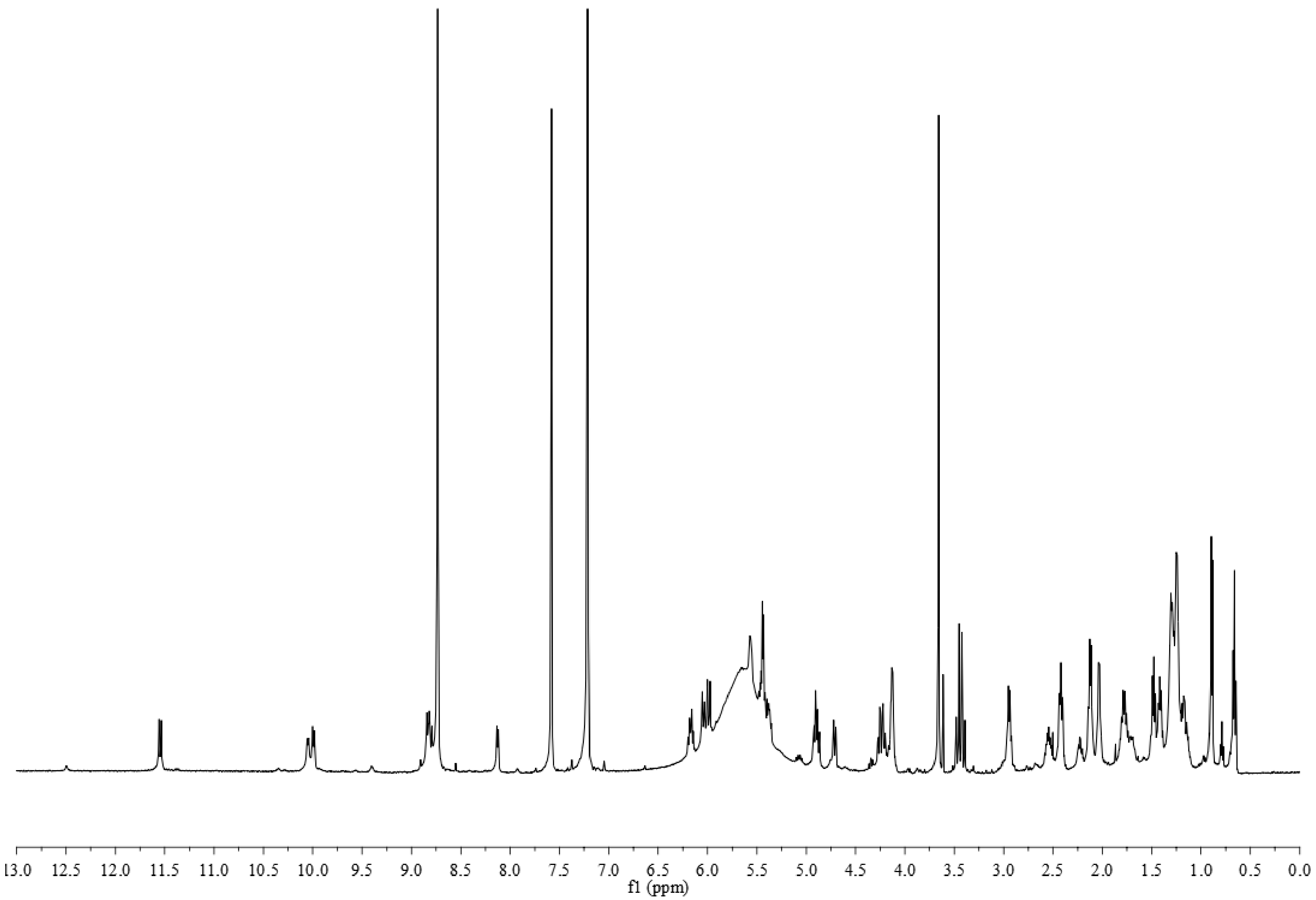


**Figure S9.** $^{1}$H NMR spectrum of gladiochelin B in pyridine-$d_5$.

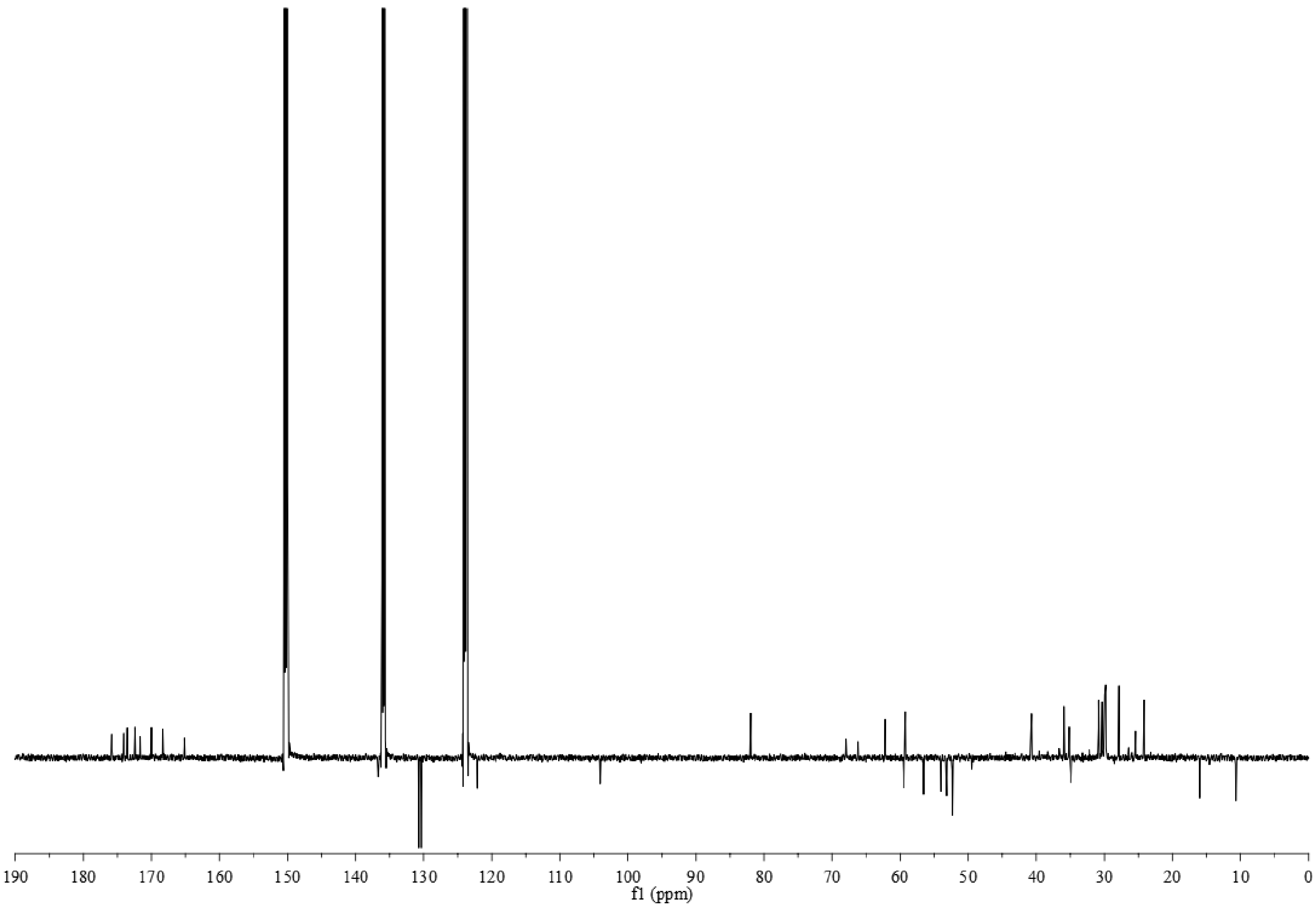


**Figure S10.** $^{13}C$ NMR spectrum of gladiochelin B in pyridine-$d_5$.

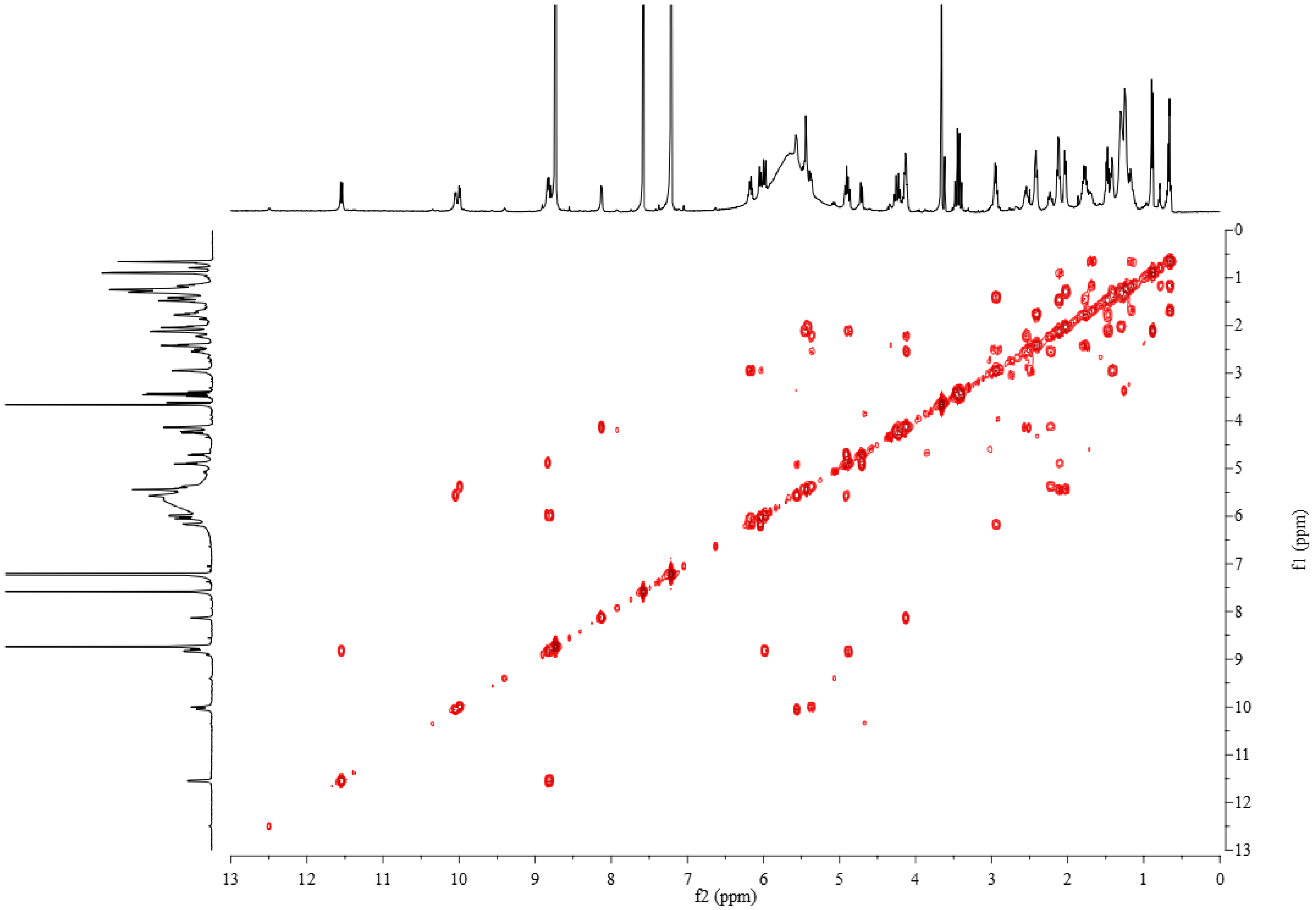


**Figure S11.** COSY spectrum of gladiochelin B in pyridine-$d_5$.

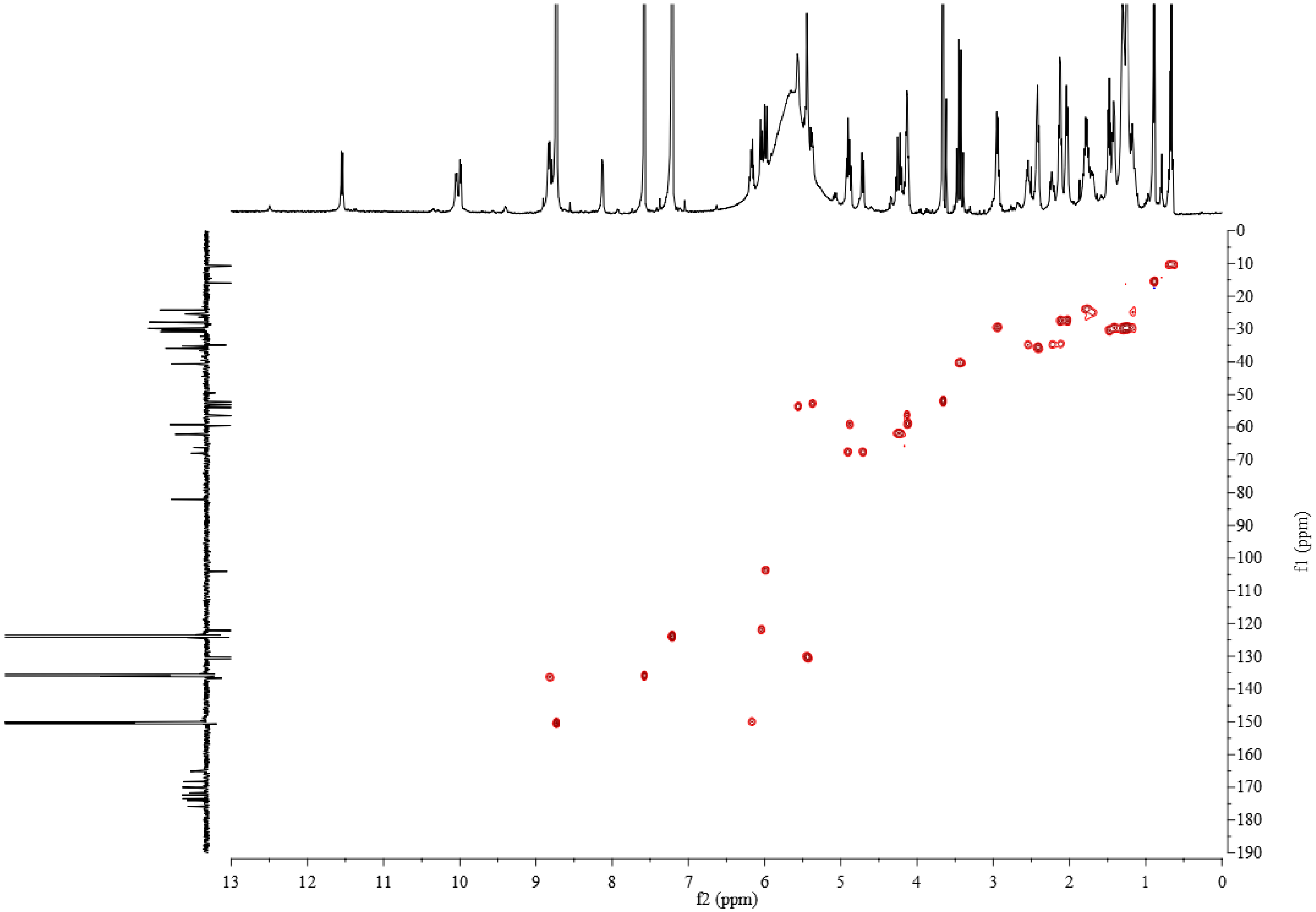


**Figure S12.** HSQC spectrum of gladiochelin B in pyridine-*$d_5$*.

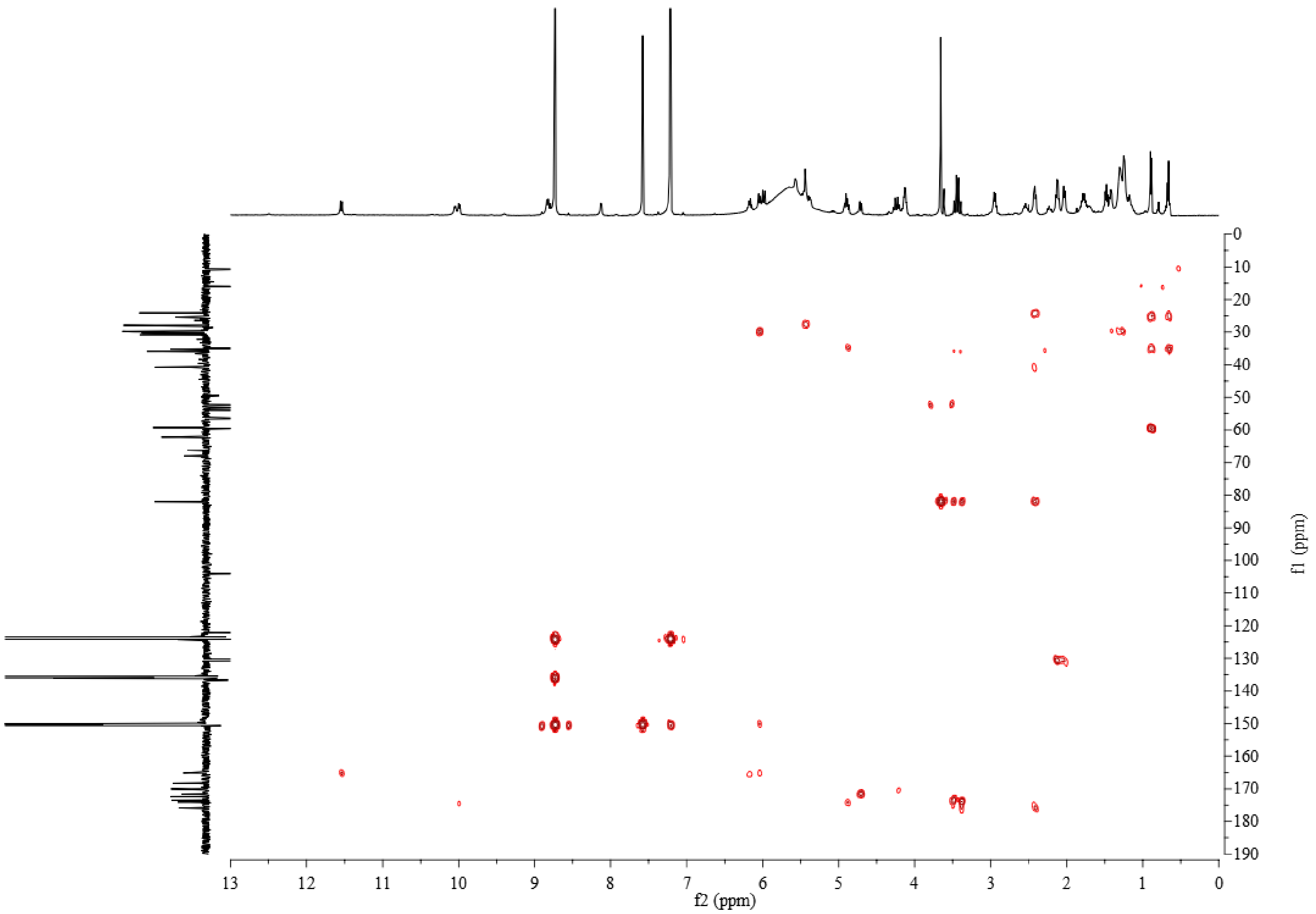


**Figure S13.** HMBC spectrum of gladiochelin B in pyridine-$d_5$.

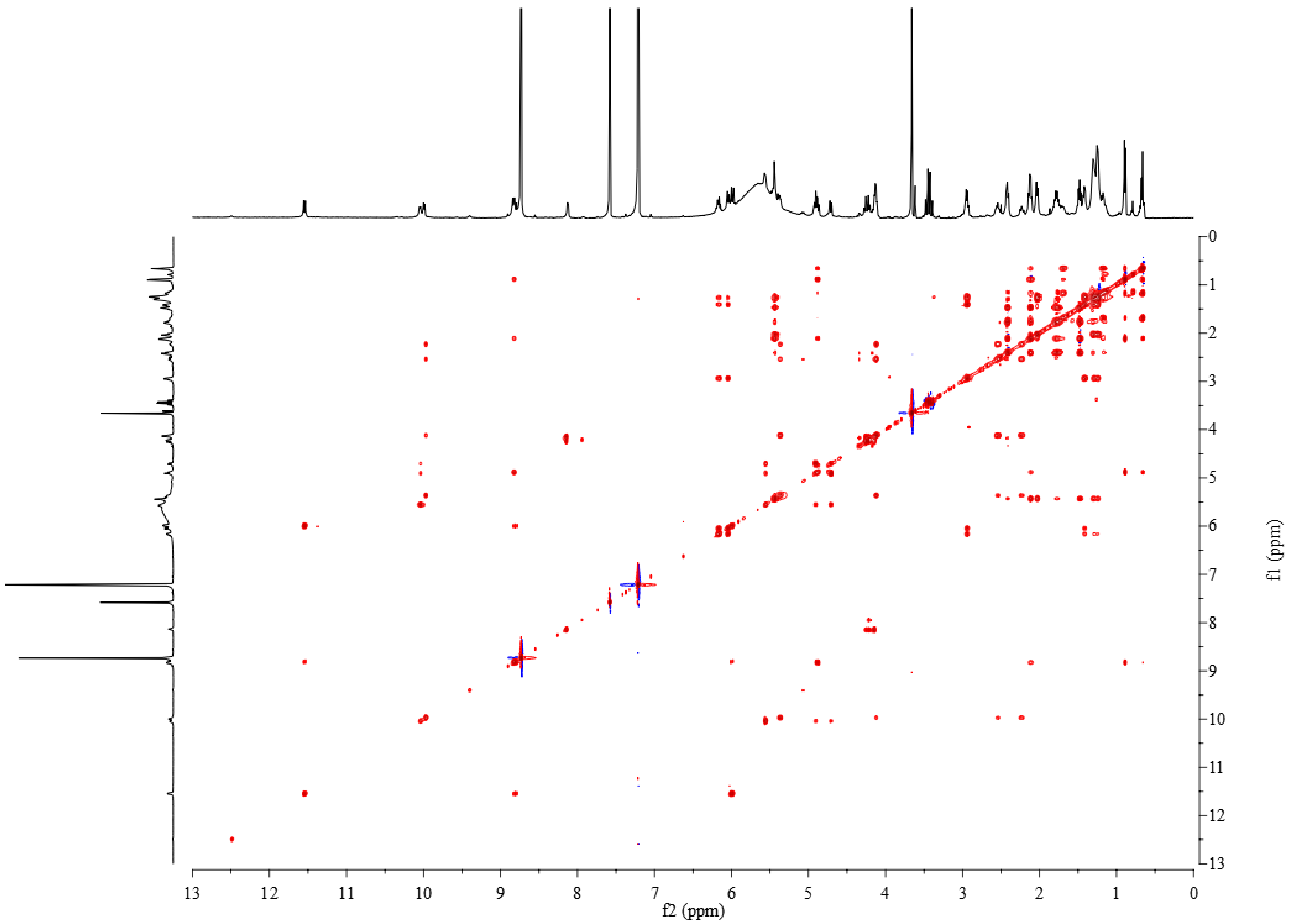


**Figure S14.** ROESY spectrum of gladiochelin B in pyridine-$d_5$.

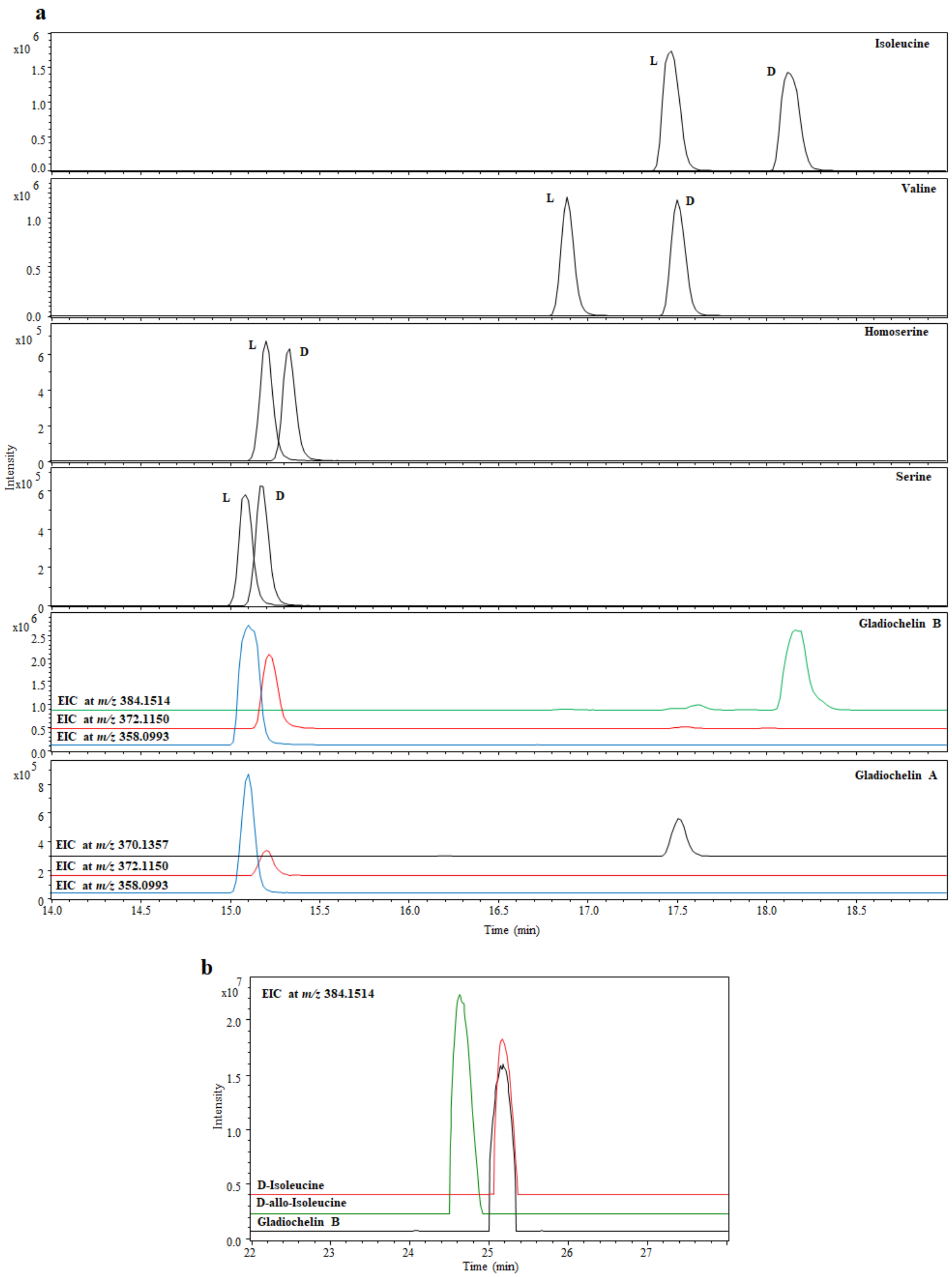


**Figure S15.** LC-MS comparison of Marfey's derivatives of gladiochelin A and B hydrolysates with authentic standards. **a**) Extracted ion chromatograms at *m/z* 358.0993, 372.1150, 370.1357, and 384.1514 corresponding to $[M+H]^+$ for Marfey's derivatives of Ser, Hse, Val, and Ile, respectively, from analyses comparing the derivatives from gladiochelin A and B hydrolysates with the derivatives of authentic standards L/D-Ser, L/D-Hse, L/D-Val, and L/D-Ile. **b**) Extracted ion chromatogram at *m/z* 384.1514, corresponding to $[M+H]^+$ for the Marfey's derivative Ile from analyses comparing the derivatives from gladiochelin B hydrolysate with D-isoleucine and D-*allo*-isoleucine.

**Table S4.** Predicted specificity pocket residues and substrates of the GcnH adenylation domains.

| A domain | Putative Specificity Pocket Residues | Predicted Substrate |
|---|---|---|
| Module 1 | D V W H M S L V (D) | L-Ser |
| Module 2 | D A L W M G G V (F) | L-Val/L-Ile |
| Module 3 | D L K N V G S D (V) | Hse |
| Module 4 | D V W H V S L I (D) | L-Ser |

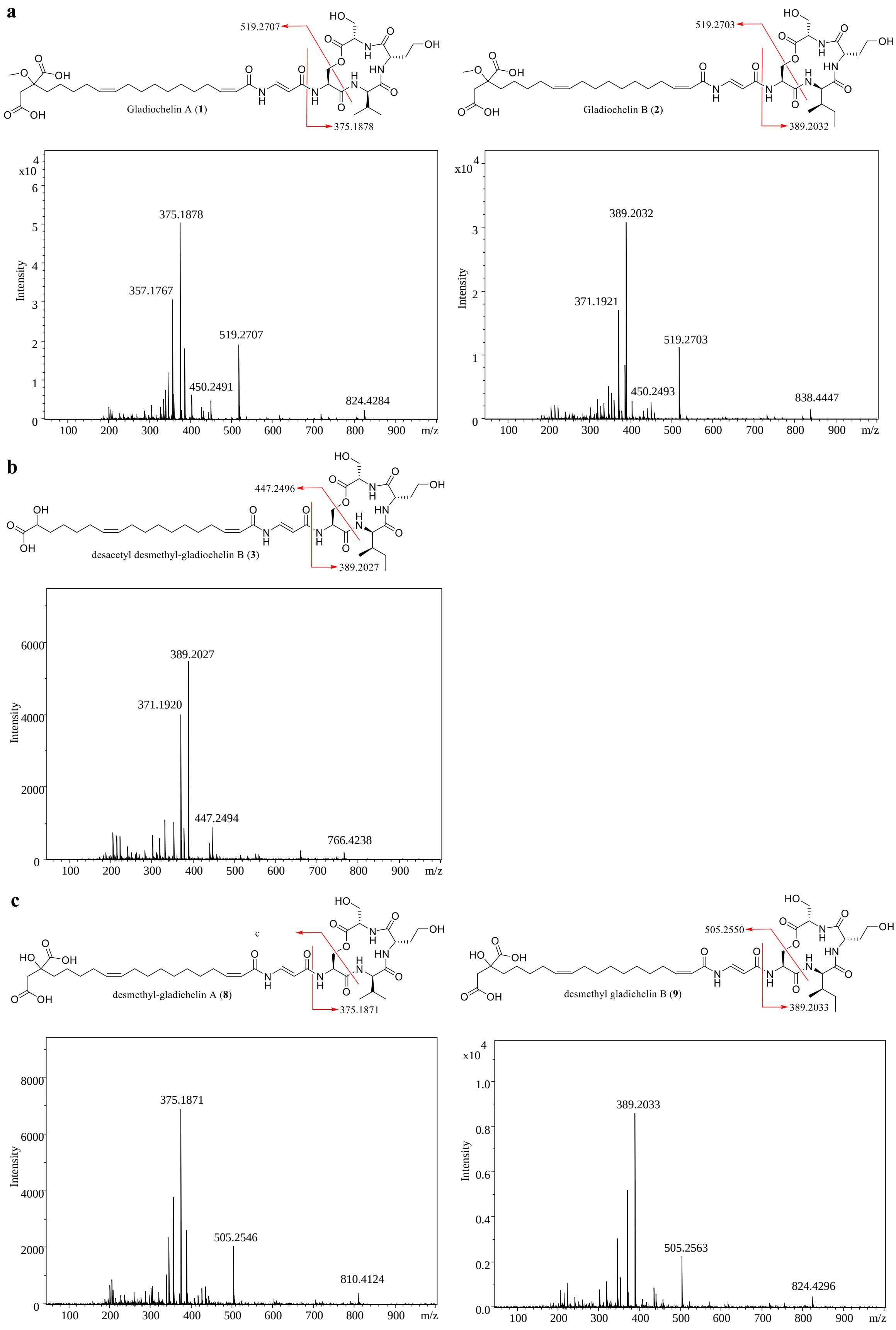


**Figure S16.** MS/MS spectra and proposed fragment ions for gladiochelins A and B, metabolite **3**, and desmethyl-gladiochelins A and B. **a)** gladiochelins A (**1**) and B (**2**); **b)** metabolite **3** accumulated by the Δ*gcnR* mutant; and **c)** desmethyl-gladiochelins A (**8**) and B (**9**) produced by the Δ*gcnS* mutant.

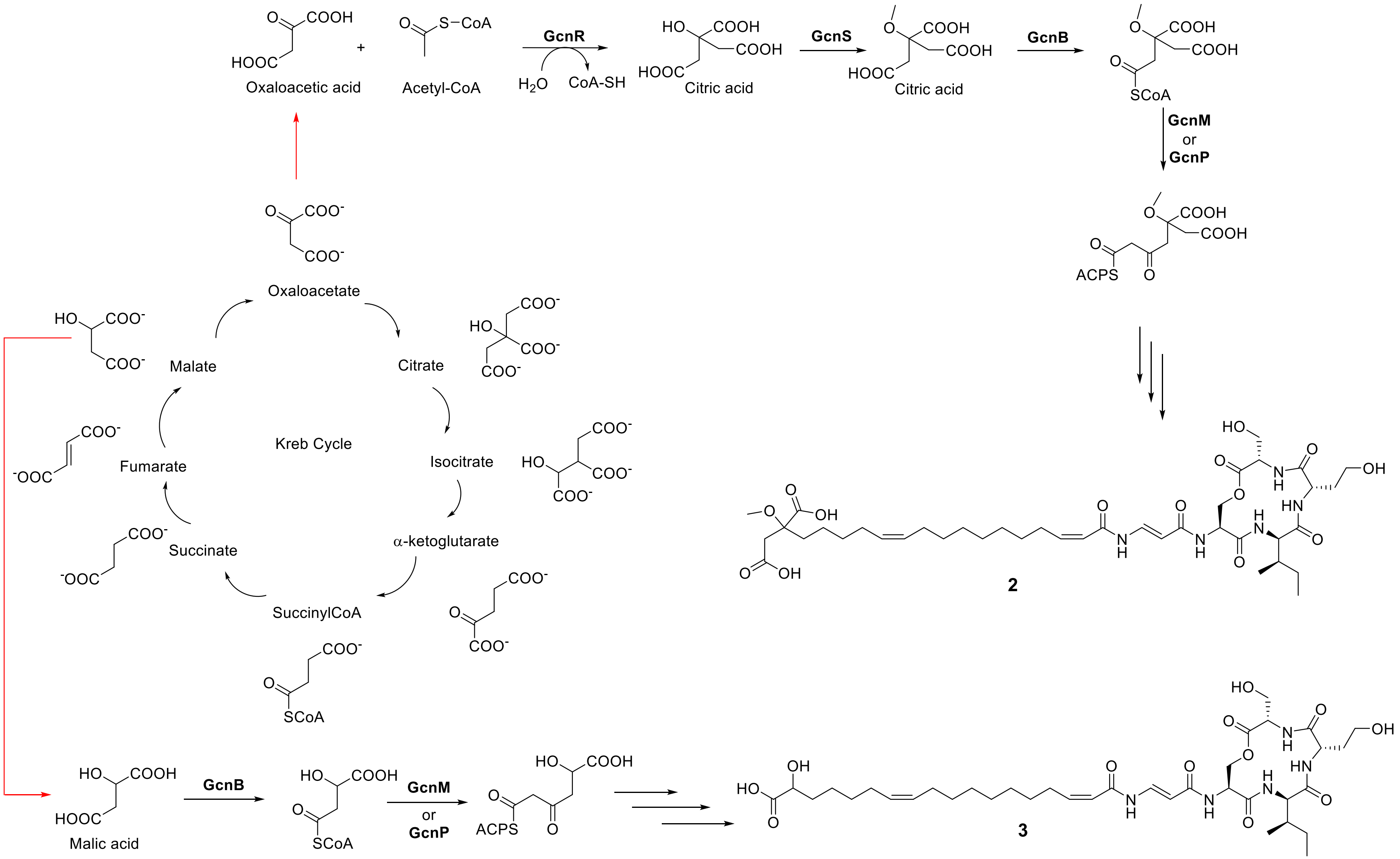


**Figure S17.** Comparison of the proposed biosynthetic pathways for galdiochelin B (**2**) and compound **3** produced by the Δ*gcnR* mutant.

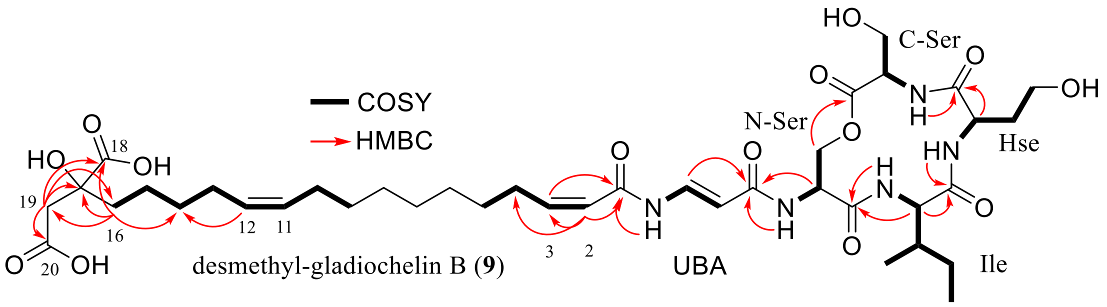


**Figure S18.** Structure of desmethyl-gladiochelin B summarising COSY and key HMBC correlations, and defining the nomenclature and numbering used in Table S5.

**Table S5.** $^{1}H$, $^{13}C$ and HMBC NMR data for desmethyl-gladiochelin B in pyridine-$d_5$.

| Position | | $\delta_C$ | $\delta_H$ | HMBC |
|---|---|---|---|---|
| C-Ser | C=O | 170.0 | - | |
| | α-C | 56.6 | 4.13 (m) | |
| | β-C | 62.20 | 4.24 (ddd, *J* = 24.4, 10.8, 3.7 Hz) | C-Ser (C=O) |
| | NH | | 8.12 (d, *J* = 7.0 Hz) | Hse (C=O) |
| Hse | C=O | 172.4 | - | |
| | α-C | 53.1 | 5.38 (m) | Hse (C=O, β-C, γ-C) |
| | β-C | 35.2 | 2.22 (m)/2.55 (m) | Hse (C=O, α-C, γ-C) |
| | γ-C | 59.2 | 4.13 (m) | Hse (α-C, β-C) |
| | NH | | 9.99 (d, *J* = 7.5 Hz) | Ile (C=O) |
| Ile | C=O | 174.0 | - | |
| | α-C | 59.5 | 4.89 (*ol*) | Ile (C=O, β-C, γ1-C, γ2-C), N-Ser (C=O) |
| | β-C | 34.9 | 2.12 (m) | Ile (γ1-C) |
| | γ1-C | 25.4 | 1.18 (m)/1.70 (m) | Ile (α-C, β-C, γ2-C, δ-C) |
| | γ2-C | 16.0 | 0.89 (d, *J* = 6.7 Hz) | Ile (α-C, β-C, γ1-C) |
| | δ-C | 10.7 | 0.66 (t, *J* = 7.4 Hz) | Ile (β-C, γ1-C) |
| | NH | | 8.82 (*ol*) | N-Ser (C=O) |
| N-Ser | C=O | 171.6 | - | |
| | α-C | 54.0 | 5.56 (m) | N-Ser (C=O), UBA (C=O) |
| | β-C | 68.0 | 4.71 (d, *J* = 10.7 Hz)/4.92 (*ol*) | N-Ser (C=O, α-C), C-Ser (C=O) |
| | NH | | 10.02 (d, *J* = 7.5 Hz) | UBA (C=O) |
| UBA | C=O | 168.3 | - | |
| | α-C | 104.1 | 5.97 (d, *J* = 13.9 Hz) | UBA (C=O, β-C) |
| | β-C | 136.7 | 8.82 (*ol*) | UBA (C=O) |
| | NH | | 11.54 (d, *J* = 11.3 Hz) | FA (C=O) |
| Fatty acid (FA) | 1 | 165.1 | - | |
| | 2 | 122.1 | 6.04 (d, *J* = 11.4 Hz) | FA (C=O, C3, C4) |
| | 3 | 150.1 | 6.17 (m) | FA (C=O) |
| | 4 | 29.8 | 2.95 (dd, *J* = 14.3, 7.1 Hz) | FA (C5) |
| | 5 | 29.9 | 1.42 (m) | FA (C4) |
| | 6 | 29.8-30.0 | 1.23-1.32 | |
| | 7 | 29.8-30.0 | 1.23-1.32 | |
| | 8 | 29.8-30.0 | 1.23-1.32 | |
| | 9 | 30.34 | 1.23-1.32 | |
| | 10 | 27.9 | 2.03 (dd, *J* = 13.0, 6.6 Hz) | FA (C11, C12) |
| | 11 | 130.7 | 5.41 (m) | |
| | 12 | 130.3 | 5.45 (m) | |
| | 13 | 27.9 | 2.11 (m) | FA (C11, C12, C15) |
| | 14 | 30.8 | 1.45 (m) | FA (C12, C13, C15, C16) |
| | 15 | 24.2 | 1.75 (m), 1.96 (m) | FA (C13, C14, C16, C17) |
| | 16 | 40.7 | 2.11 (m), 2.21 (m) | FA (C14, C15, C17, C18, C19) |
| | 17 | 76.1 | - | |
| | 18 | 178.9 | - | |
| | 19 | 45.6 | 3.25 (d, *J* = 15.7)/ 3.57 (d, *J* = 15.7) | FA (C16, C17, C18, C20) |
| | 20 | 174.4 | - | |

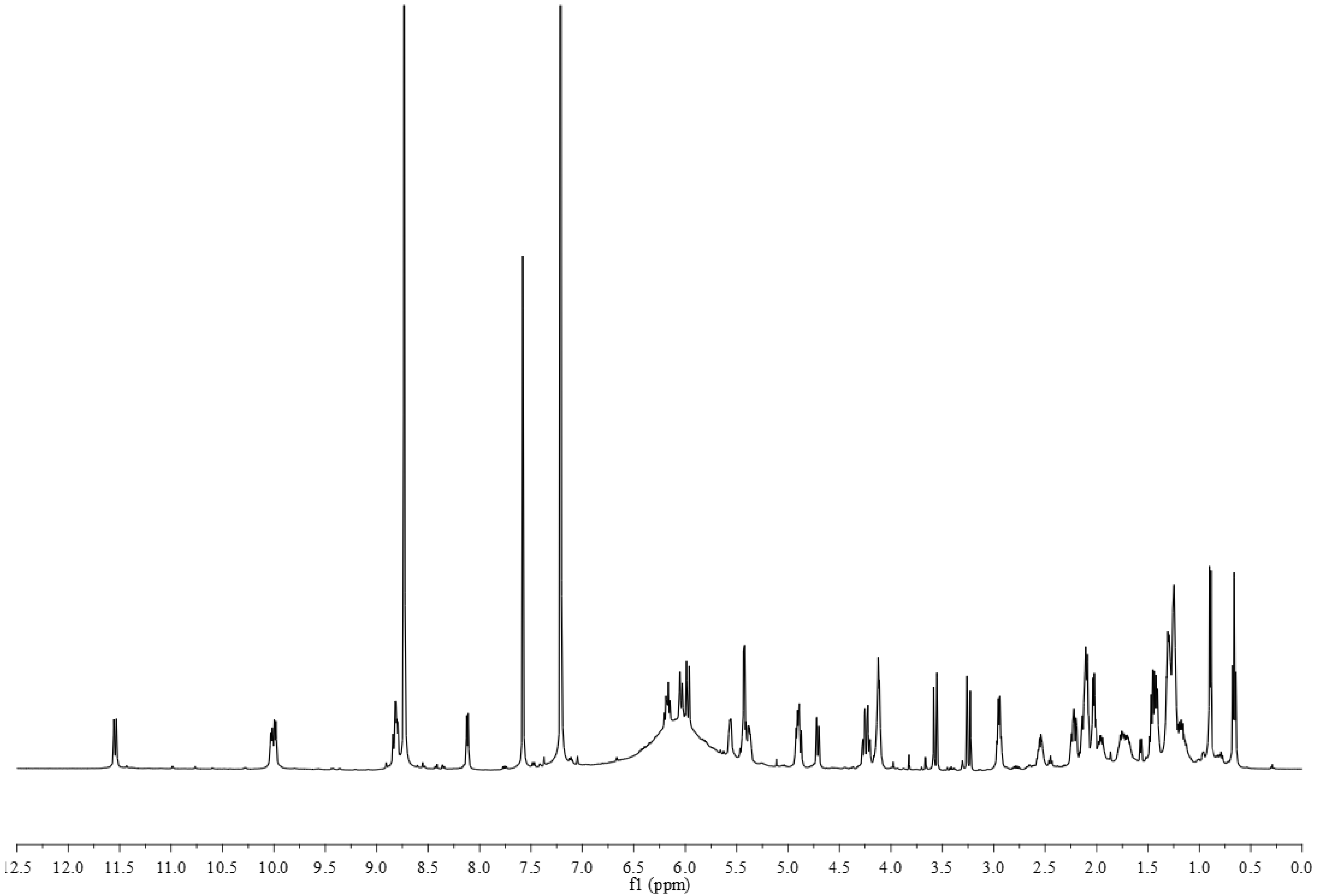


**Figure S19.** $^1$H NMR spectrum of desmethyl-gladiochelin B in pyridine-$d_5$.

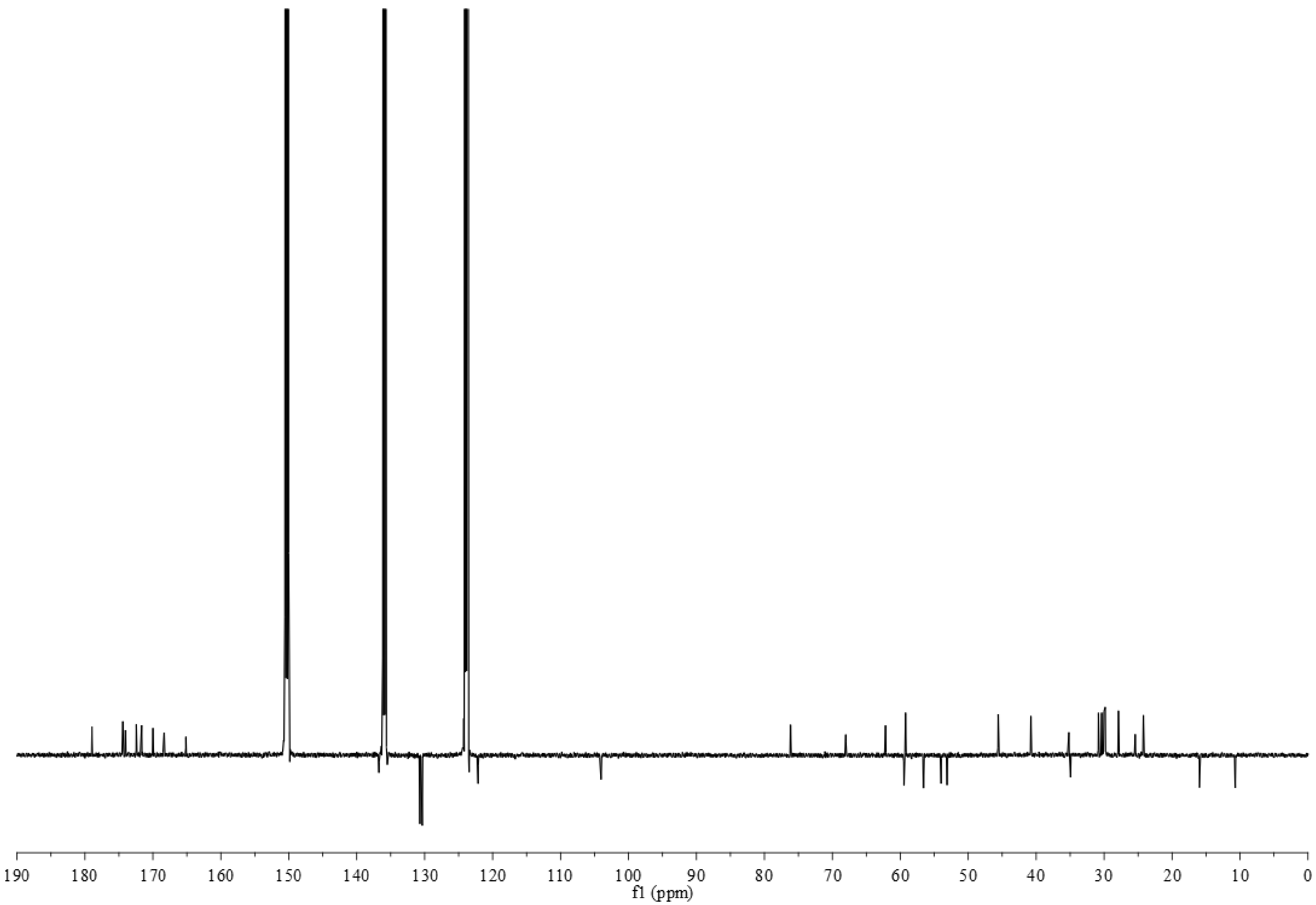


**Figure S20.** $^{13}$C NMR spectrum of desmethyl-gladiochelin B in pyridine-$d_5$.

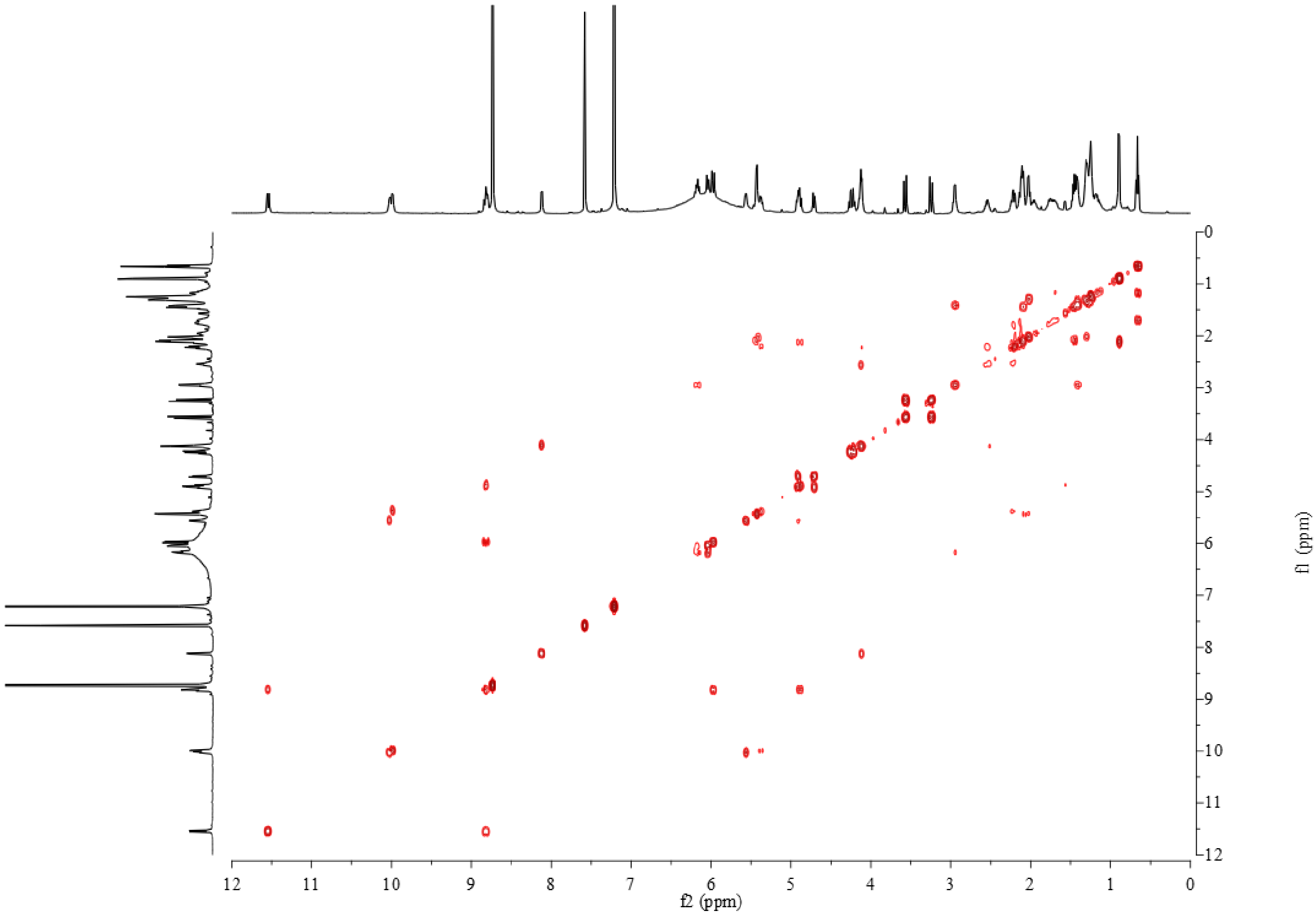


**Figure S21.** COSY spectrum of desmethyl-gladiochelin B in pyridine-*d*₅.

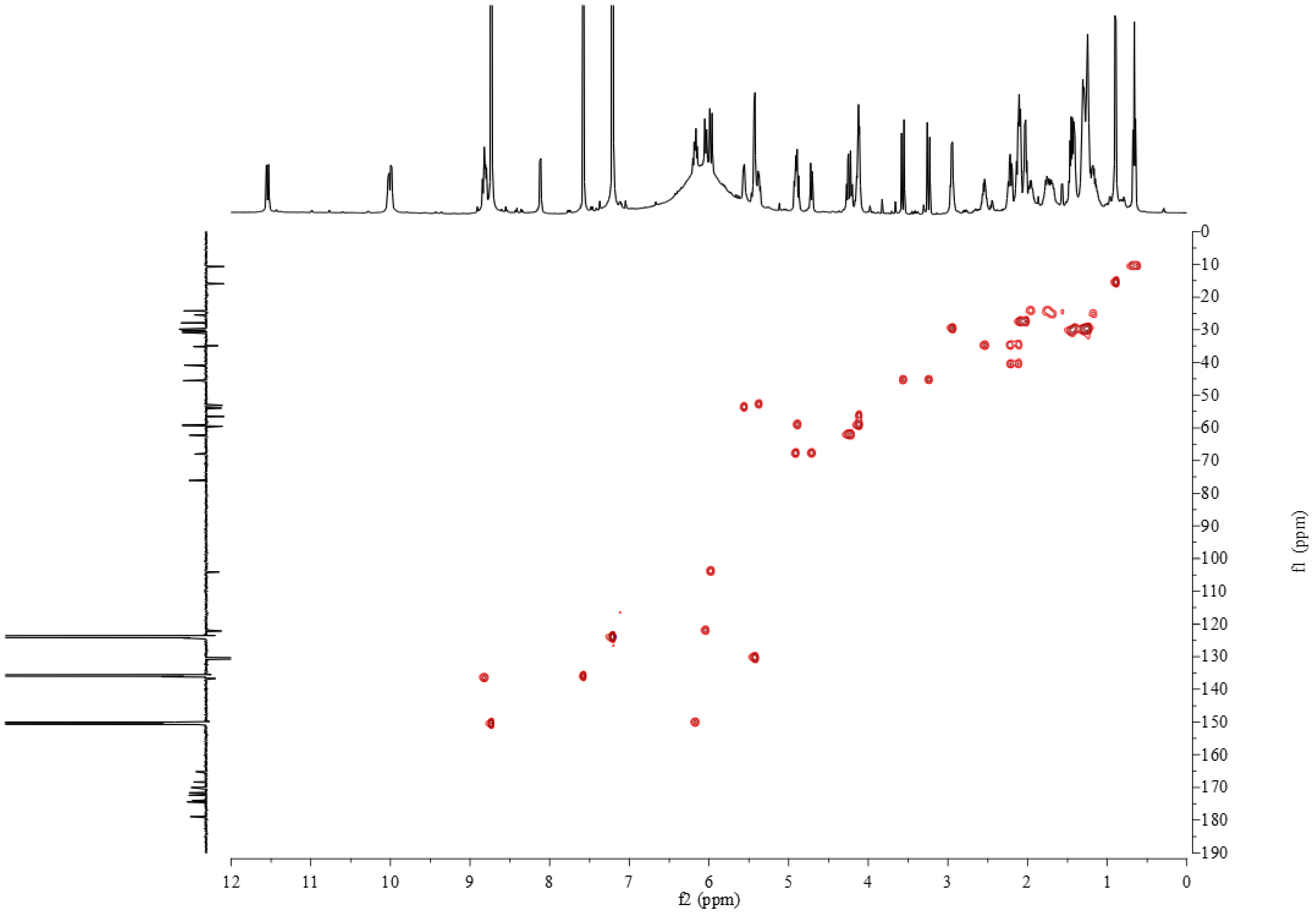


**Figure S22.** HSQC spectrum of desmethyl-gladiochelin B in pyridine-$d_5$.

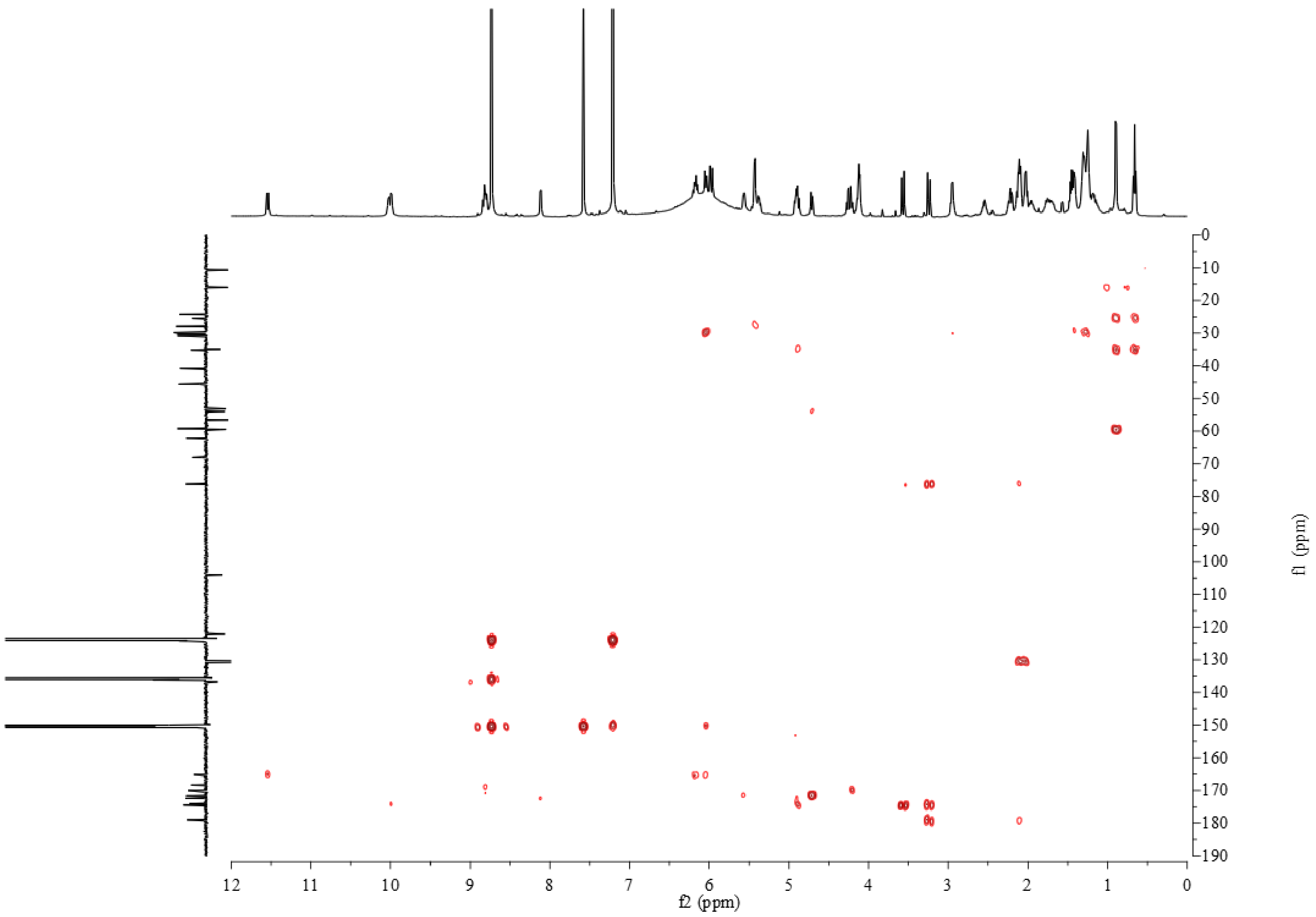


**Figure S23.** HMBC spectrum of desmethyl-gladiochelin B in pyridine-$d_5$.

```
                                                          230
VinN            QQIVFAASSINAVLGYHAEDIVFCRMSVSWDFGLYKVLISTLTGAKLVLAGGEPDIALVK
GcnO            RNLVSGAFSVAAYQGLEAHDVILGVLPLSFDAGLSQLTTALVSGACYAPLDFLRPSEVPK
                :::* .* *: *  * .*.*:::  :.:*:* ** ::  : ::**  .  .      : *
                                                                   299
VinN            SLRESGATMMPIVPSLASMLTTLIRRDPEGAPTLRMFTNSAAALPQVTIDALRSAFPGAQ
GcnO            YCEARGVTSITAVPPLWMQLASVEWS-EAARQRIRRFANTGGHLATPLLNRLEALYPRAA
                  .  *.* :. **.*   *:::      .   :* *:*:.. *.   :: *.: :* *
                          331
VinN            VVRMYGQTECKRISIMPPHLEHERPDSVGLPLPGTTIEILDEDGTLLPPGEPGEITVTGP
GcnO            PYLMYGLTEAFRSTYLPPAEARQRPSSIGKAVPNAEILVLRADGSECGVDEPGELVHRGA
                   *** **. * : :**   ::**.*:* .:*.: * :*  **:    .****:.  *.
```

**Figure S24.** Sequence alignment of VinN and GcnO. Residues known (VinN) and predicted (GcnO) to make polar contacts with the α-carboxylate group in the β-methyl-Asp and Asp substrates, respectively, of these adenylating enzymes are highlighted in yellow.

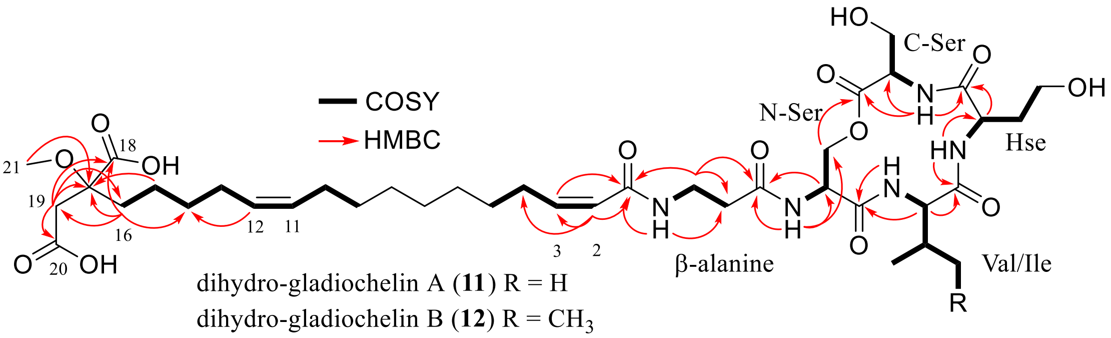


dihydro-gladiochelin A (**11**) R = H
dihydro-gladiochelin B (**12**) R = $CH_3$

**Figure S25.** Structure of dihydro-gladiochelins A and B summarising COSY and key HMBC correlations, and defining the nomenclature and numbering used in tables S6 and S7.

**Table S6.** $^{1}$H, $^{13}$C and HMBC NMR data for dihydro-gladiochelin A in pyridine-$d_5$.

| Position | | $\delta_C$ | $\delta_H$ | HMBC |
|---|---|---|---|---|
| C-Ser | C=O | 170.17 | - | |
| | α-C | 56.45 | 4.33 (m, 1H) | C-Ser (C=O) |
| | β-C | 62.08 | 4.27 (m, 1H) | C-Ser (C=O) |
| | NH | | 8.30 (dd, *J* = 7.1 Hz, 1H) | C-Ser (α-C, C=O), Hse (C=O) |
| Hse | C=O | 172.48 | - | |
| | α-C | 53.10 | 5.34 (qd, *J* = 7.6, 6.8, 3.8 Hz, 1H) | Hse (β-C) |
| | β-C | 35.07 | 2.23(m, 1H)/ 2.51 (m, 1H) | Hse (α- C, γ-C, C=O) |
| | γ-C | 59.20 | 4.10 (m, 1H) | Hse (α- C) |
| | NH | | 9.89 (d, *J* = 7.6 Hz, 1H) | Hse (α-C, β-C, C=O) |
| Val | C=O | 173.75 | - | |
| | α-C | 61.18 | 4.71 (t, *J* = 10.1 Hz, 1H) | Val (β-C, γ-C), N-Ser (C=O) |
| | β-C | 29.39 | 2.23 (ddd, *J* = 15.2, 10.2, 5.2 Hz, 1H) | |
| | γ-C | 19.44<br>19.79 | 0.95 (d, *J* = 6.6 Hz, 2H)<br>0.92 (d, *J* = 6.6 Hz, 2H) | Val (α-C, β-C) |
| | NH | | 8.75 (*ol* with solvent peak) | N-Ser (C=O) |
| N-Ser | C=O | 171.52 | - | |
| | α-C | 54.14 | 5.34 (m, 1H) | N-Ser (C=O) |
| | β-C | 67.36 | 4.62 (dd, *J* = 10.9, 2.5 Hz, 1H)/ 4.90 (dd, *J* = 10.9, 4.2 Hz, 1H) | C-Ser (C=O), N-Ser (α-C) |
| | NH | | 10.06 (d, *J* = 7.4 Hz, 1H) | N-Ser (α-C, β-C, C=O) |
| β–alanine | C=O | 173.45 | - | |
| | α-C | 36.69 | 2.81 (m, 1H)/ 2.63 (m, 1H) | β-ala (β-C, C=O) |
| | β-C | 36.32 | 3.89 (q, *J* = 6.6 Hz, 2H) | β-ala (β-C), FA (C1), N-Ser (C=O) |
| | NH | | 8.77 (*ol* with solvent peak) | |
| Fatty acid (FA) | 1 | 167.41 | - | |
| | 2 | 123.75 | 6.11 (d, *J* = 11.0 Hz, 1H) | FA (C=O, C3, C4) |
| | 3 | 145.45 | 6.0 (dt, *J* = 11.5, 7.3 Hz, 1H) | FA (C=O, C2, C4) |
| | 4 | 29.39 | 2.96 (q, *J* = 6.7 Hz, 2H) | FA (C2, C3, C5) |
| | 5 | 29.80 | 1.42 (m, 1H) | FA (C7) |
| | 6 | 30.00 | 1.23-1.30 | |
| | 7 | 30.04 | 1.23-1.30 | |
| | 8 | 30.11 | 1.23-1.30 | |
| | 9 | 30.31 | 1.23-1.30 | |
| | 10 | 27.82 | 2.09 (q, *J* = 6.8 Hz, 2H) | FA (C9, C12) |
| | 11 | 130.71 | 5.42 (m, 1H) | FA (C13) |
| | 12 | 130.25 | 5.42 (m, 1H) | |
| | 13 | 27.84 | 2.11 (q, *J* = 7.0 Hz, 2H) | FA (C11, C14, C15) |
| | 14 | 30.79 | 1.45 (m, 1H) | FA (C13, C15, C16) |
| | 15 | 24.11 | 1.76 (m, 1H) | |
| | 16 | 35.89 | 2.41 (t, *J* = 8.4 Hz, 2H) | FA (C15, C17, C19) |
| | 17 | 81.93 | - | |
| | 18 | 175.75 | - | |
| | 19 | 40.64 | 3.43 ( q, *J* = 15.1 Hz, 2H) | FA (C16, C17, C20) |
| | 20 | 173.19 | - | |
| | 21 | 52.31 | 3.65 (s, 3H) | FA (C17) |

**Table S7.** $^{1}H$, $^{13}C$ and HMBC NMR data for dihydro-gladiochelin B in pyridine-$d_5$.

| Position | | $\delta_C$ | $\delta_H$ | HMBC |
|---|---|---|---|---|
| C-Ser | C=O | 170.12 | - | |
| | α-C | 56.45 | 4.10 (m, 1H) | |
| | β-C | 62.07 | 4.26 (td, *J* = 11.6, 10.5, 3.8 Hz, 1H) | C-Ser (C=O) |
| | NH | | 8.26 (d, *J* = 6.7 Hz, 1H) | C-Ser (α-C) Hse (C=O) |
| Hse | C=O | 172.47 | - | |
| | α-C | 53.00 | 5.34 (m, 1H) | |
| | β-C | 35.09 | 2.22 (m, 1H)/ 2.52 (m, 1H) | Hse (α-C, C=O) |
| | γ-C | 59.19 | 4.10 (q, *J* = 5.1 Hz, 1H) | Hse (α-C) |
| | NH | | 10.10 (d, *J* = 5.1 Hz, 1H) | Hse (α-C) Ile (C=O) |
| Ile | C=O | 173.86 | - | |
| | α-C | 59.42 | 4.83 (t, *J* = 10.1 Hz, 1H) | Ile (β-C, γ1-C, γ2-C, C=O) |
| | β-C | 35.09 | 2.20 (dd, *J* = 13.8, 6.8 Hz, 1H) | |
| | γ1-C | 25.41 | 1.15 (m, 1H) / 1.60 (m, 1H) | Ile (α-C, β-C, γ2-C, δ-C) |
| | γ2-C | 15.94 | 0.90 (d, *J* = 6.6 Hz, 1H) | Ile (α-C, β-C, γ1-C) |
| | δ-C | 10.60 | 0.67 (t, *J* = 7.4 Hz, 1H) | Ile (β-C, γ1-C) |
| | NH | | 8.80 (d, *J* = 9.3 Hz, 1H) | N-Ser (59.42, C=O) |
| N-Ser | C=O | 171.48 | - | |
| | α-C | 54.06 | 5.35 (m, 1H) | |
| | β-C | 67.51 | 4.64 (dd, *J* = 10.9, 2.1 Hz, 1H)/ 4.90 (dd, *J* = 10.8, 4.1 Hz, 1H) | N-Ser (α-C, C=O) |
| | NH | | 9.89 (d, *J* = 7.6 Hz, 1H) | N-Ser (α-C) β-ala(α-C, β-C) |
| β–alanine | C=O | 173.45 | - | |
| | α-C | 36.65 | 2.65 (m, 1H)/ 2.81 (m, 1H) | β-ala (β-C) |
| | β-C | 36.29 | 3.89 (q, *J* = 6.5 Hz, 2H) | β-ala (α-C, C=O) FA(C1) |
| | NH | | 8.75 (*ol* with solvent peak) | β-ala (α-C, β-C) FA (C1) |
| Fatty acid (FA) | 1 | 166.57 | - | |
| | 2 | 123.73 | 6.11 (d, *J* = 11.5 Hz, 1H) | FA (C1, C4) |
| | 3 | 145.44 | 6.00 (dt, *J* = 11.6, 7.3 Hz, 1H) | FA (C1, C5) |
| | 4 | 29.78 | 2.96 (q, *J*= 7.5 Hz, 1H) | |
| | 5 | 29.83 | 1.42 (m, 1H) | |
| | 6 | 29.98 | 1.20-1.33 | |
| | 7 | 30.02 | 1.20-1.33 | |
| | 8 | 30.08 | 1.20-1.33 | |
| | 9 | 30.29 | 1.20-1.33 | |
| | 10 | 27.80 | 2.02 (q, *J* = 6.8 Hz, 2H) | FA (C9, C11, C12) |
| | 11 | 130.69 | 5.42 (m, 1H) | FA (C10) |
| | 12 | 130.23 | 5.42 (m, 1H) | |
| | 13 | 27.82 | 2.10 (q, *J* = 6.8 Hz, 2H) | FA (C11, C14, C15) |
| | 14 | 30.77 | 1.47 (m, 1H) | FA (C10, C11) |
| | 15 | 24.09 | 1.77 (m, 1H) | FA (C12) |
| | 16 | 35.87 | 2.40 (t, *J* = 8.4 Hz, 1H) | FA (C15, C14, C17, C18) |
| | 17 | 81.91 | - | |
| | 18 | 175.75 | - | |
| | 19 | 40.62 | 3.42 (q, *J* = 15.1 Hz, 2H) | FA (C16, C17, C20, C21) |
| | 20 | 173.19 | - | |
| | 21 | 52.29 | 3.65 (s, 3H) | FA (C17) |

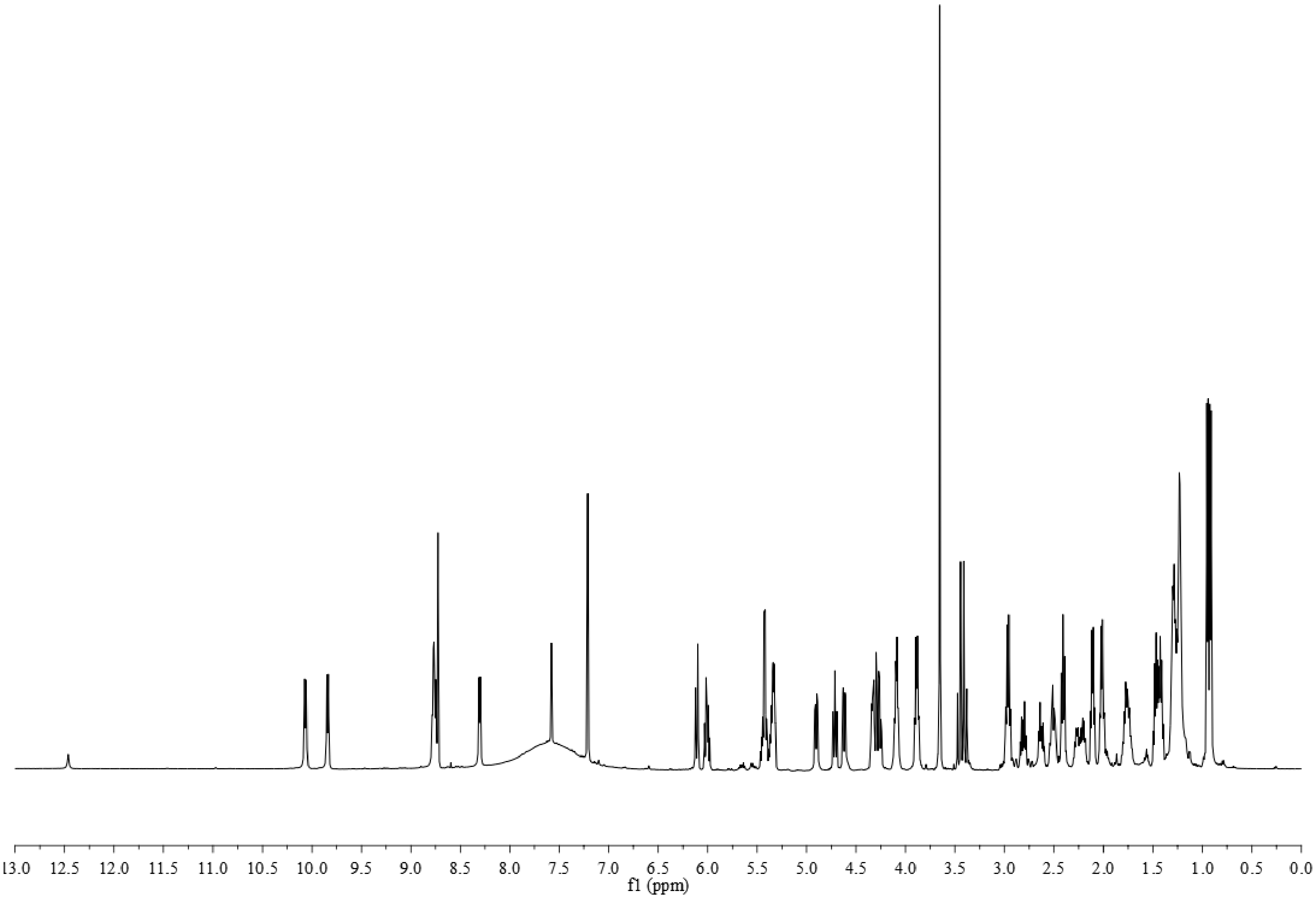


**Figure S26.** [1]H NMR spectrum of dihydro-gladiochelins A in pyridine-$d_5$.

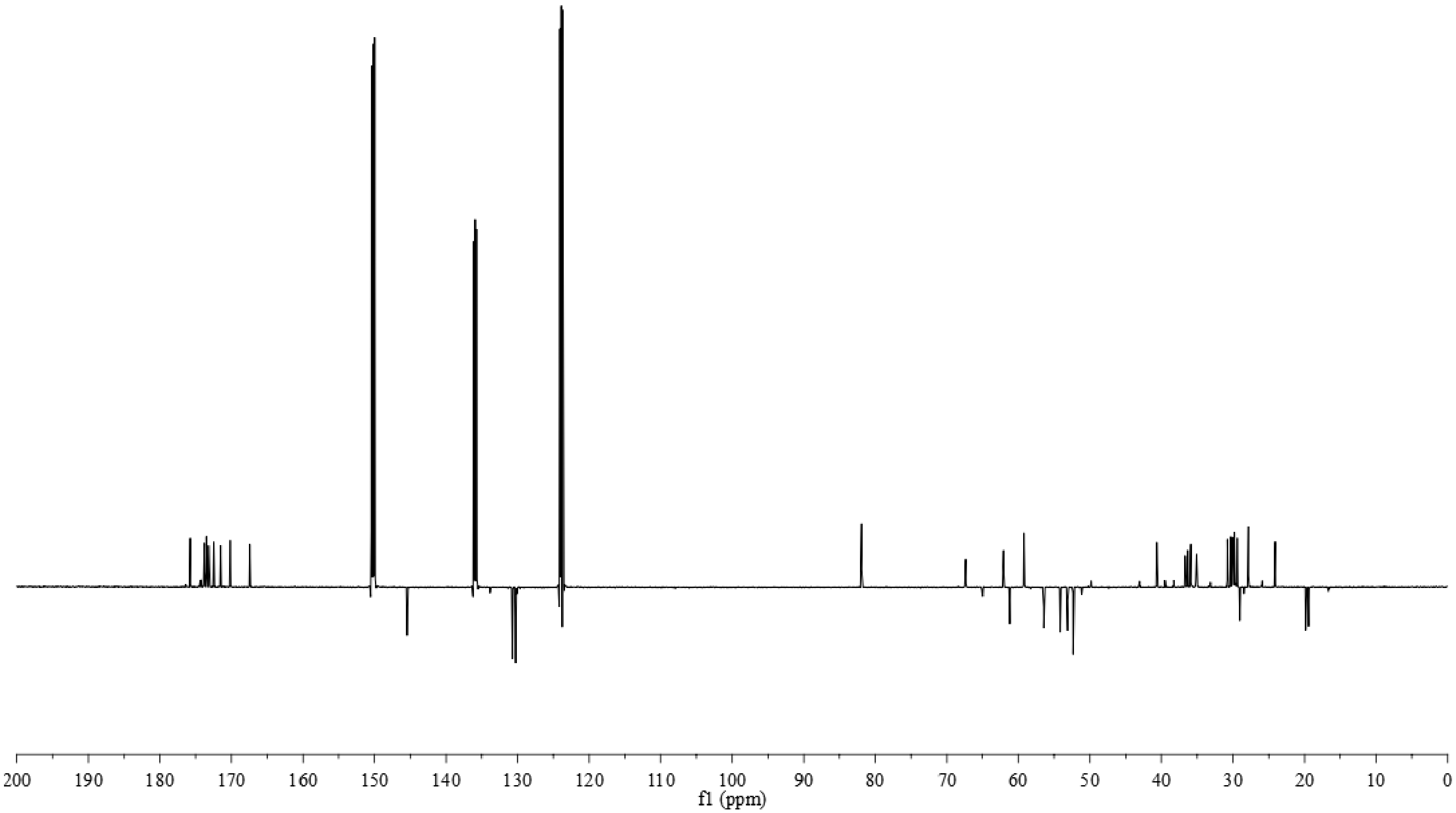


**Figure S27.** $^{13}$C NMR spectrum of dihydro-gladiochelins A in pyridine-$d_5$.

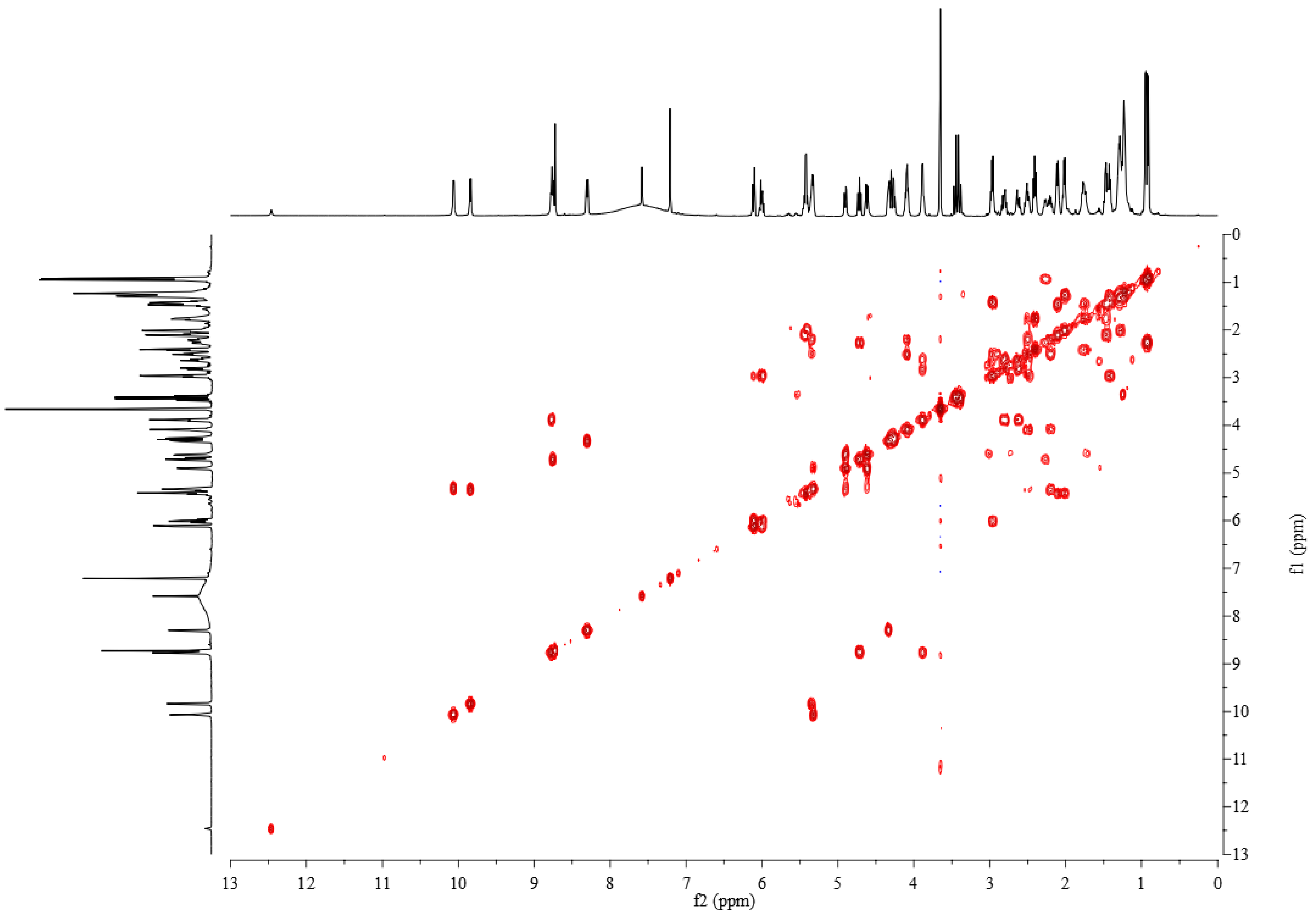


**Figure S28.** COSY spectrum of dihydro-gladiochelins A in pyridine-$d_5$.

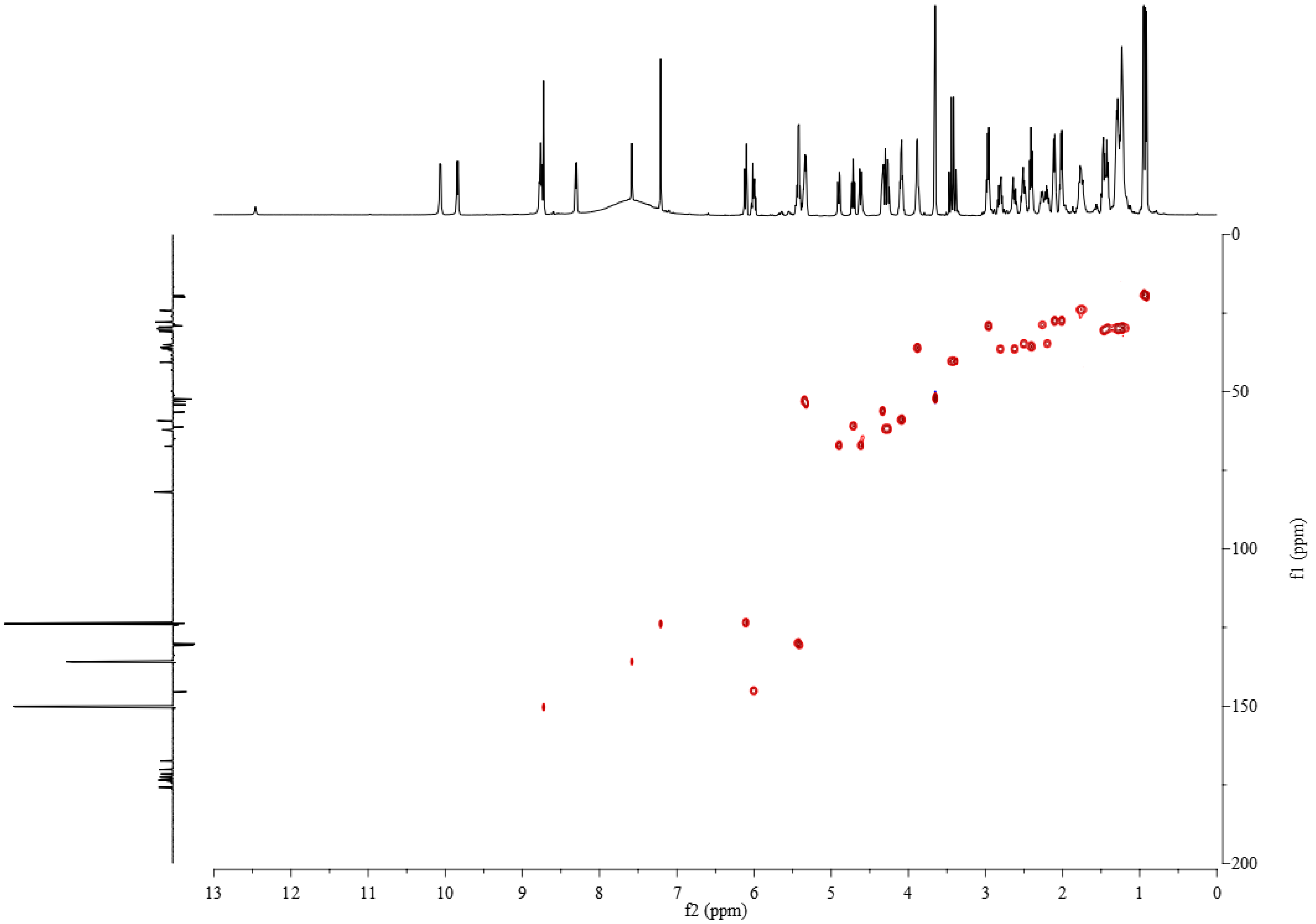

**Figure S29.** HSQC spectrum of dihydro-gladiochelins A in pyridine-$d_5$.

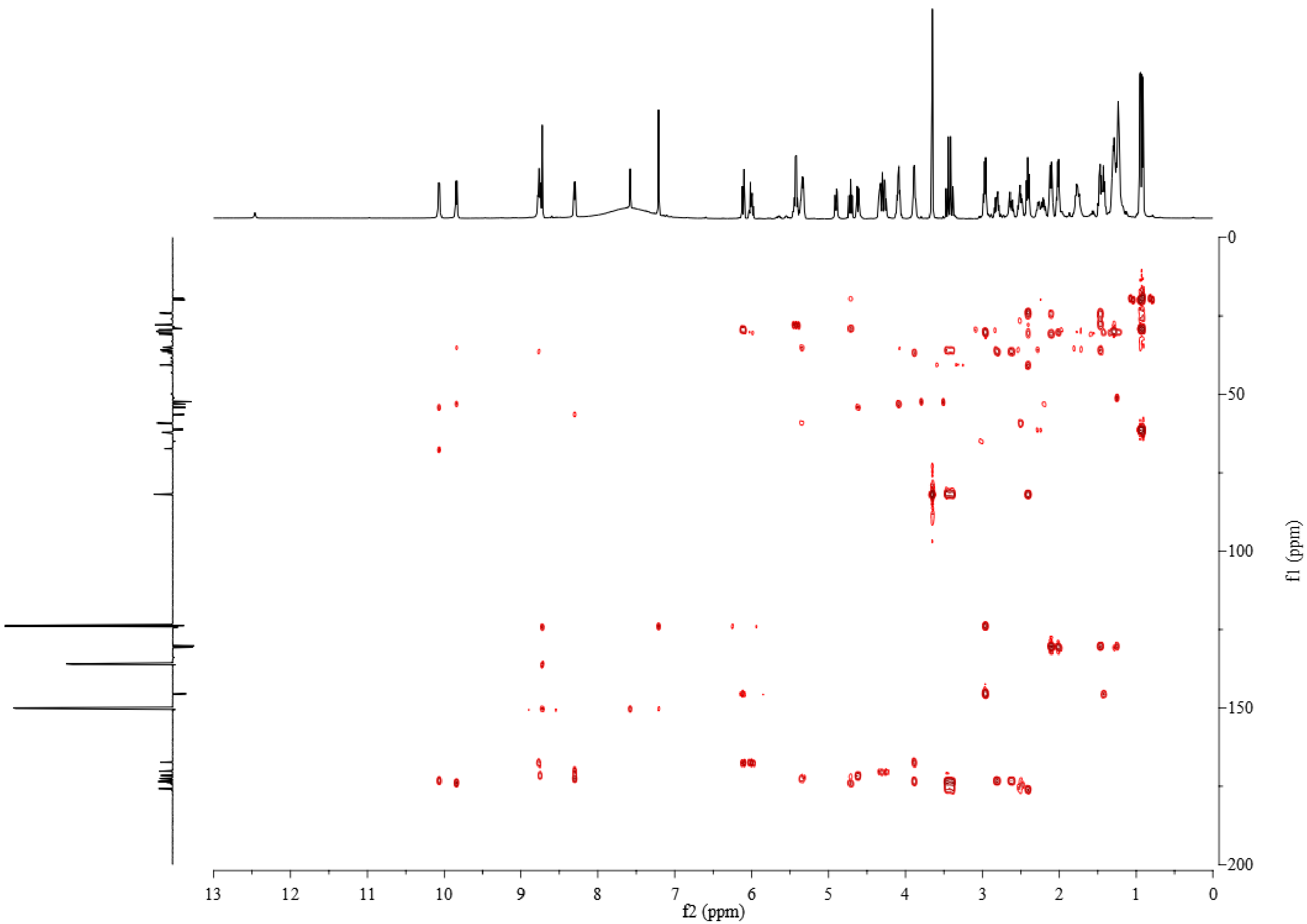


**Figure S30.** HMBC spectrum of dihydro-gladiochelins A in pyridine-$d_5$.

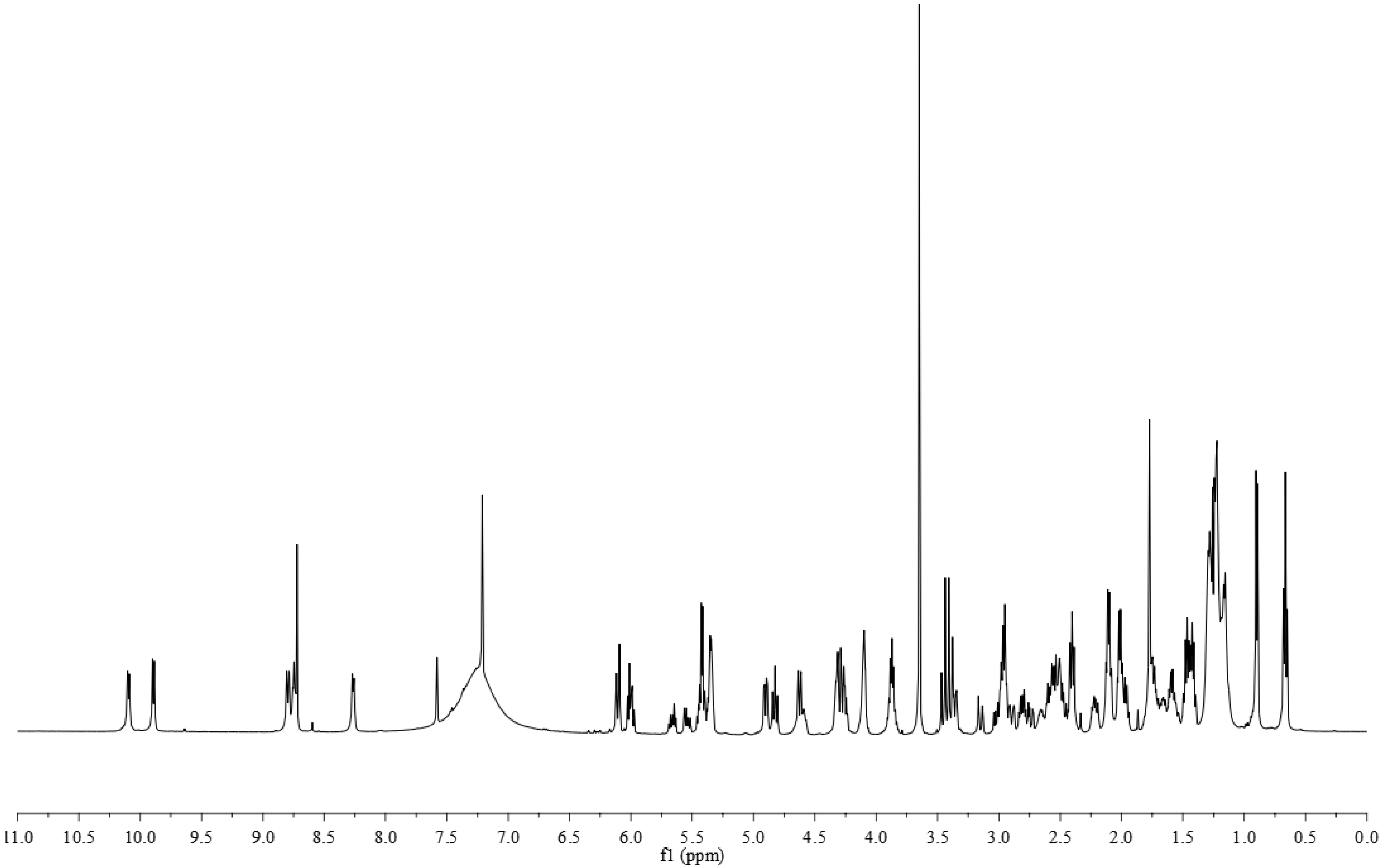


**Figure S31.** $^{1}H$ NMR spectrum of dihydro-gladiochelins B in pyridine-$d_5$.

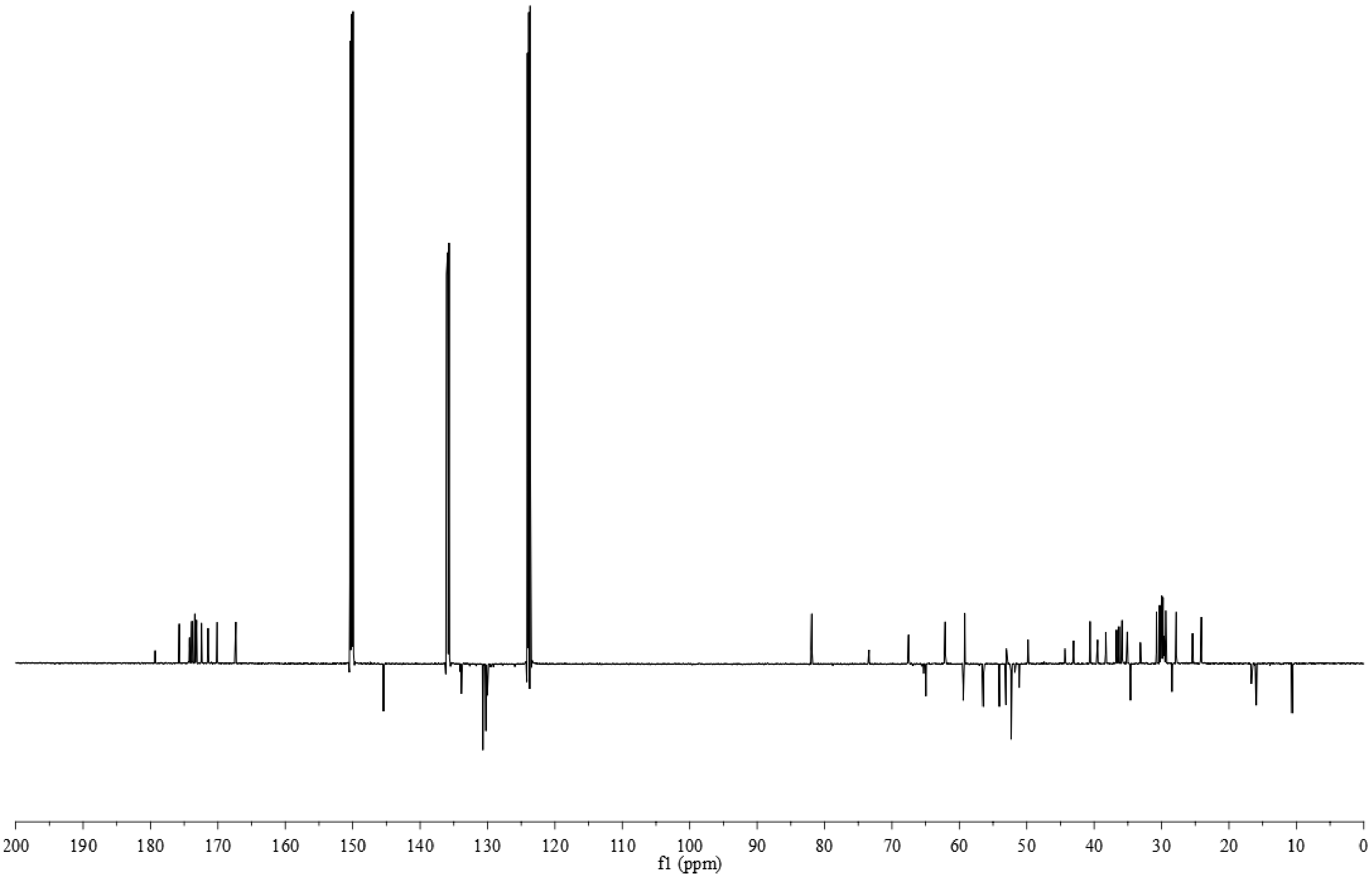


**Figure S32.** $^{13}$C NMR spectrum of dihydro-gladiochelins B in pyridine-$d_5$.

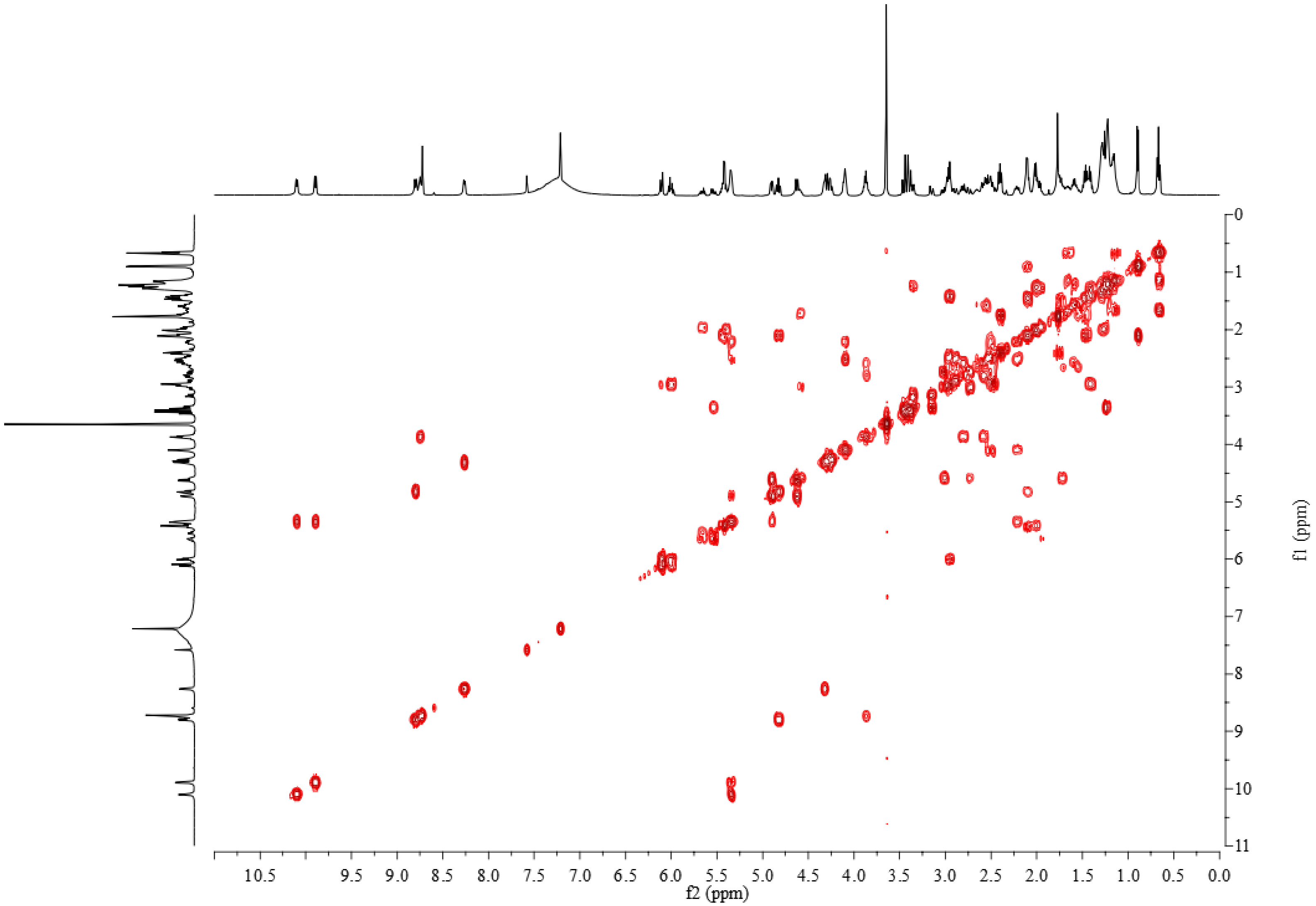


**Figure S33.** COSY spectrum of dihydro-gladiochelins B in pyridine-$d_5$.

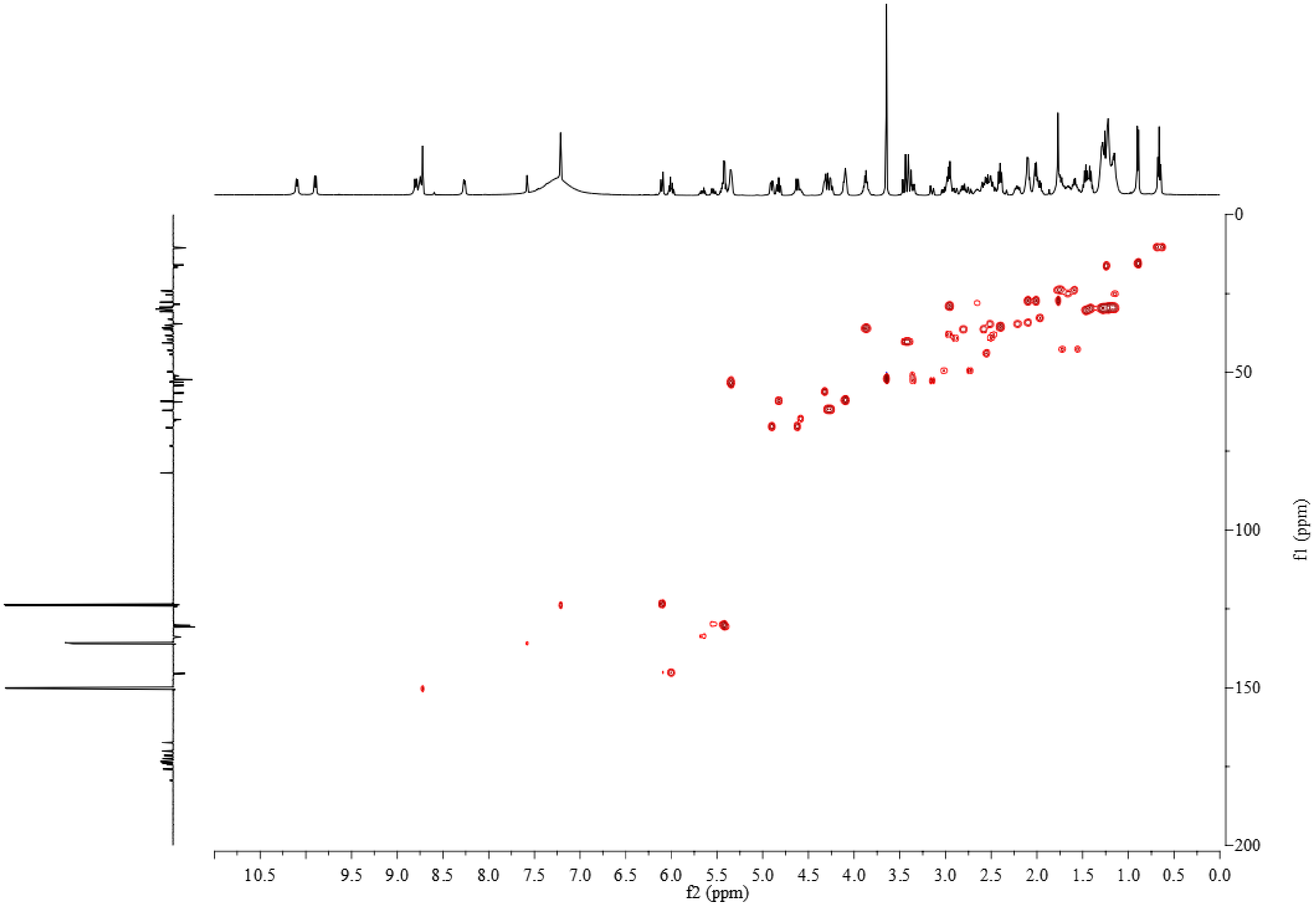


**Figure S34.** HSQC spectrum of dihydro-gladiochelins B in pyridine-$d_5$.

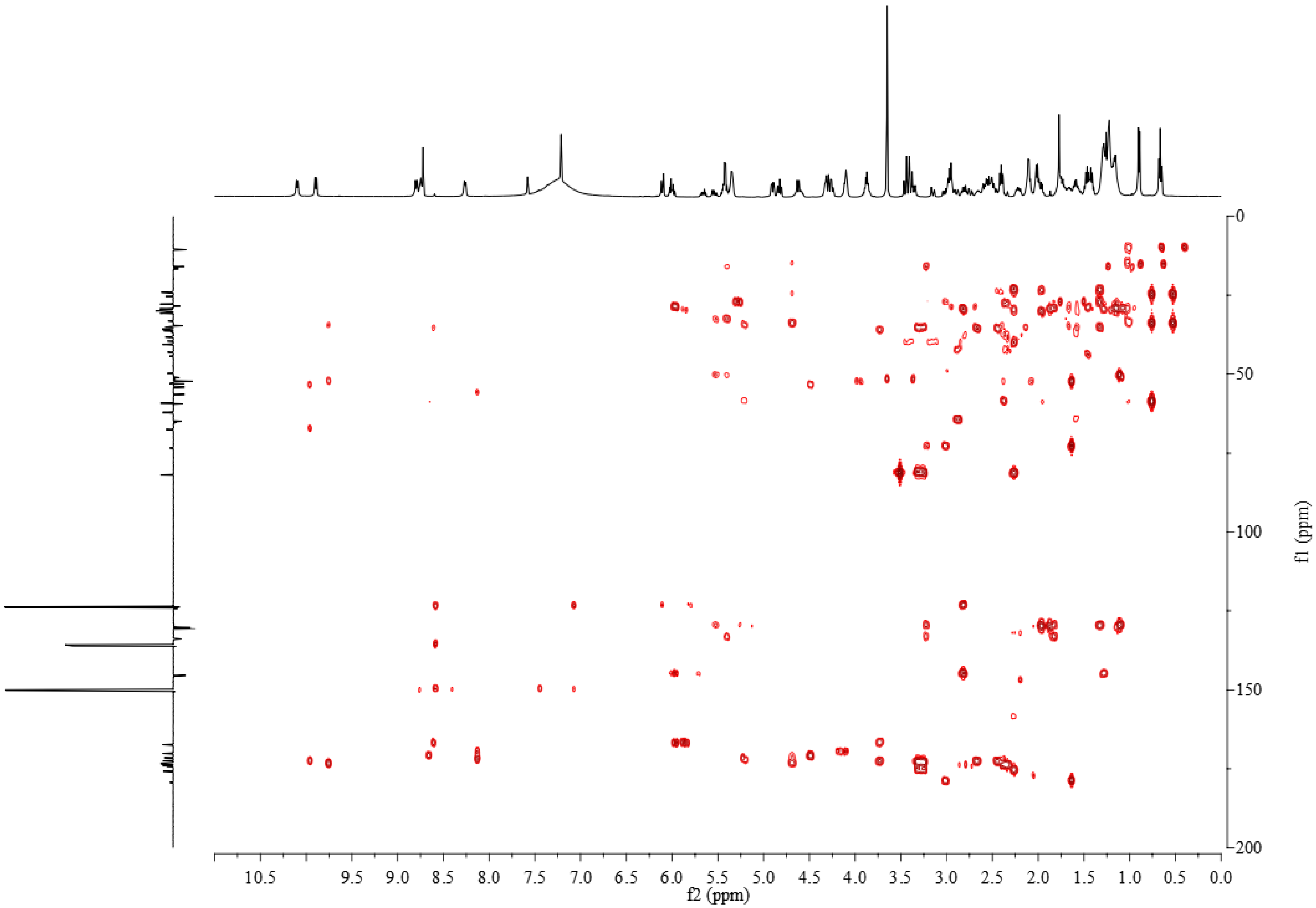


**Figure S35.** HMBC spectrum of dihydro-gladiochelins B in pyridine-$d_5$.

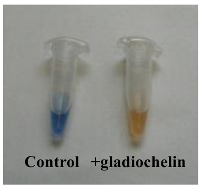


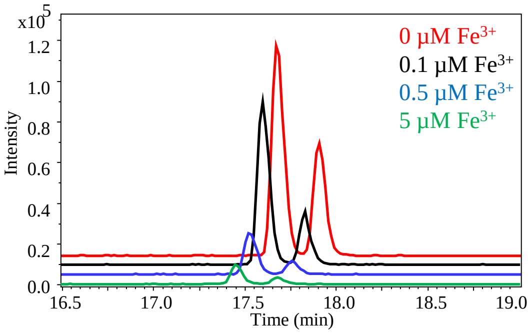


**Figure S36.** Results of CAS assay with gladiochelin A (top) and LC-MS chromatograms showing gladiochelin production is suppressed by addition of ferric iron to the growth medium (bottom).

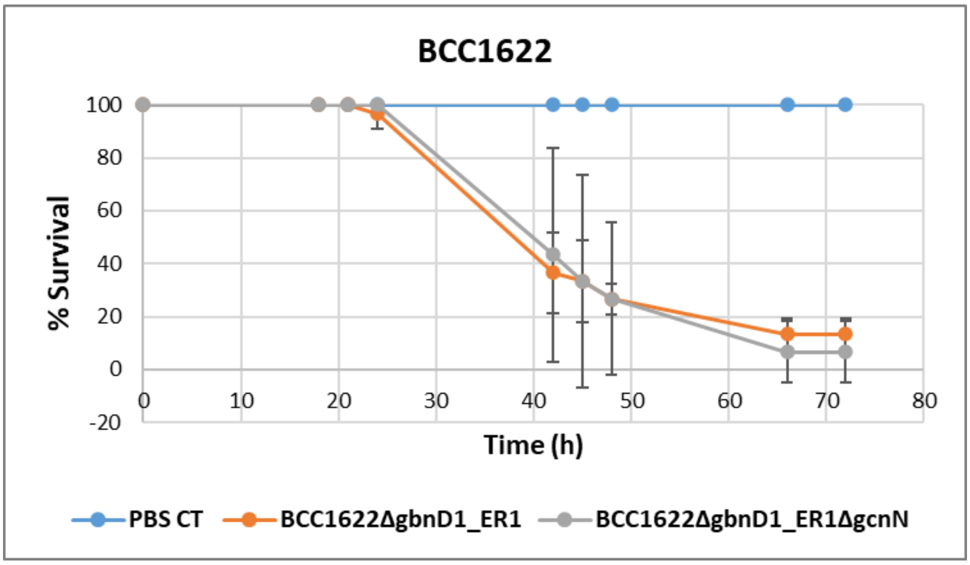


**Figure S37.** Results of virulence assay of *B. gladioli* BCC1622 and *B. gladioli* BCC1622 Δ*gbnD1_ER1*-Δ*gcnN* using the *Galleria* wax moth larvae model. Experiments were performed in triplicate using 10 larvae per treatment.
